# Expansion Sequencing: Spatially Precise *In Situ* Transcriptomics in Intact Biological Systems

**DOI:** 10.1101/2020.05.13.094268

**Authors:** Shahar Alon, Daniel R Goodwin, Anubhav Sinha, Asmamaw T Wassie, Fei Chen, Evan R Daugharthy, Yosuke Bando, Atsushi Kajita, Andrew G Xue, Karl Marrett, Robert Prior, Yi Cui, Andrew C Payne, Chun-Chen Yao, Ho-Jun Suk, Ru Wang, Chih-Chieh (Jay) Yu, Paul Tillberg, Paul Reginato, Nikita Pak, Songlei Liu, Sukanya Punthambaker, Eswar P. R. Iyer, Richie E Kohman, Jeremy A Miller, Ed S Lein, Ana Lako, Nicole Cullen, Scott Rodig, Karla Helvie, Daniel L Abravanel, Nikhil Wagle, Bruce E Johnson, Johanna Klughammer, Michal Slyper, Julia Waldman, Judit Jané-Valbuena, Orit Rozenblatt-Rosen, Aviv Regev, IMAXT Consortium, George M Church, Adam H Marblestone, Edward S Boyden

## Abstract

Methods for highly multiplexed RNA imaging are limited in spatial resolution, and thus in their ability to localize transcripts to nanoscale and subcellular compartments. We adapt expansion microscopy, which physically expands biological specimens, for long-read untargeted and targeted *in situ* RNA sequencing. We applied untargeted expansion sequencing (ExSeq) to mouse brain, yielding readout of thousands of genes, including splice variants and novel transcripts. Targeted ExSeq yielded nanoscale-resolution maps of RNAs throughout dendrites and spines in neurons of the mouse hippocampus, revealing patterns across multiple cell types; layer-specific cell types across mouse visual cortex; and the organization and position-dependent states of tumor and immune cells in a human metastatic breast cancer biopsy. Thus ExSeq enables highly multiplexed mapping of RNAs, from nanoscale to system scale.

**One Sentence Summary:** *In situ* sequencing of physically expanded specimens enables multiplexed mapping of RNAs at nanoscale, subcellular resolution.

## Main Text

Tissues are made of cells of many different types and states, whose spatial organization governs the interactions that yield healthy functions and disease states. This spatial architecture is being widely explored by multiplexed measurements of the locations and identities of RNA molecules within cells, in the context of three-dimensional cellular morphology (*1–11*). Mapping the subcellular locations of RNAs in intact tissues is important for understanding diverse biological processes (*12, 13*), such as in the nervous system, where axons and dendrites can extend macroscopic distances, but RNAs can be localized to nanoscale compartments like dendritic spines, where they regulate synaptic function (*14–17*). Imaging RNAs within such compartments, and throughout detailed cellular morphologies, requires nanoscale precision, which is not easily achieved within tissues with the optical methods commonly used for multiplexed imaging of RNA. Indeed, no method is currently available for the multiplexed imaging of RNA within tissues in the context of nanoscale cellular morphology (note that although seqFISH+ allows high resolution imaging of RNA molecules using Gaussian centroid fitting and rounds of barcode stratification, the detailed cellular and tissue context, i.e. protein distribution and cellular morphology, is not itself resolved with nanoscale precision (*18*)). Ideally one would be able to perform the enzymatic reactions of sequencing *in situ* with practically unlimited multiplexing capacity, while simultaneously providing for scalable, and multi-modal, nanoscale imaging of cellular and tissue context. We here present a toolbox for the untargeted (i.e., not restricted to a pre-defined list of gene targets) and targeted *in situ* sequencing of RNAs within intact tissues, which enable transcripts to be resolved within the context of nanoscale cellular morphology.

We first created an untargeted *in situ* sequencing technology that enables the sequencing of arbitrary RNAs, or parts of RNAs, within detailed cellular and tissue contexts. Untargeted approaches have the potential to discover spatially localized sequence variants, such as splice variants and functional or retained introns (*19*). Fluorescent *in situ* sequencing (FISSEQ) enables such data to be acquired from cultured cells (*20*), but full untargeted sequencing was not demonstrated in tissues (*20*), perhaps because the dense molecular environment hampers reagent access for *in situ* sequencing. That idea motivated us to adapt the chemistry of expansion microscopy (ExM; (*21, 22*)) to decrowd RNAs from each other, and from other nearby molecules like proteins. We reasoned that this could facilitate the chemical access needed for *in situ* sequencing to work well in tissues, while anticipating that the resolution boost from ExM would enable high spatial resolution mapping of the RNAs within cellular and tissue context on conventional microscopes.

In FISSEQ, untargeted *in situ* sequencing of RNA is performed by first enzymatically amplifying mRNA locally (via reverse transcription of RNA to cDNA, cDNA circularization, and rolling circle amplification) to form ‘nanoballs’ of cDNA (termed ‘amplicons’), which contain many copies of the original mRNA sequence (*20, 23*). These sequences can be interrogated *in situ* with standard next-generation sequencing chemistries (e.g. sequencing by ligation, SOLiD) on a standard fluorescence microscope. In ExM (*21*) we physically magnify tissues in an isotropic fashion by infusing a preserved tissue specimen with monomers that polymerize into a dense mesh of highly swellable hydrogel (e.g., sodium polyacrylate), which binds via anchoring molecules to endogenous biomolecules such as proteins and RNA, or to applied labels like antibodies. The resulting tissue-polymer composite is then chemically softened so that it expands by ∼4x in linear dimension upon immersion in water. The result is isotropic decrowding of gel-anchored biomolecules of interest, which facilitates both nanoscale imaging with conventional optics, and better chemical access to the decrowded biomolecules (*22*). It has already been observed that ExM enables individual RNA transcripts, normally densely packed, to be better resolved for amplified and/or multiplexed *in situ* hybridization imaging (*24, 25*). Expanding a specimen would potentially benefit FISSEQ in the same way, by dividing the effective size of the FISSEQ amplicon (200-400 nm; (*20*)) by the expansion factor, reducing the packing density of observed amplicons and facilitating their tracking over many rounds of sequencing, important for sequence reconstruction.

We discovered that, with several critical innovations, ExM chemistry could be adapted to enable FISSEQ in expanded tissues. In particular, the traditional anchoring (**Fig. 1A**), polymerization (**Fig. 1B**), and expansion (**Fig. 1C**) steps, while effective at isotropically separating RNAs for nanoscale imaging (*24*), result in an environment full of highly charged carboxylic acid groups, which suppress multiple enzymatic reactions required for FISSEQ (**Fig. S1**). We thus developed a strategy for neutralizing these charges without causing shrinkage of the gel, by first stabilizing the specimen in the expanded state by re-embedding it in an uncharged gel (*24*), and then chemically passivating the carboxylate groups so that they become charge neutral (**Fig. S1**). We reasoned that this would allow FISSEQ signal amplification (**Fig. 1D**) and readout (**Fig. 1E, 1F, 1G**) steps to proceed. Since sequencing involves many rounds of adding fluorescent oligonucleotides one by one, to read out the biological information, we accordingly established an automated system for fluid handling and spinning disk confocal imaging, using only off-the-shelf parts for end user simplicity (see **Methods** for details). Because the resultant datasets consist of a series of 3D images of the same fields of view, one for each successive base sequenced, we also created a software pipeline (**Fig. S2**) capable of aligning, across images, the puncta for each expressed gene to within 1 pixel, over many imaging rounds, as validated by normalized cross-correlation analysis (**Fig. S3** and **Fig. S4**), followed by puncta segmentation (**Fig. 1Giii**) and base calling.

**Fig. 1.**
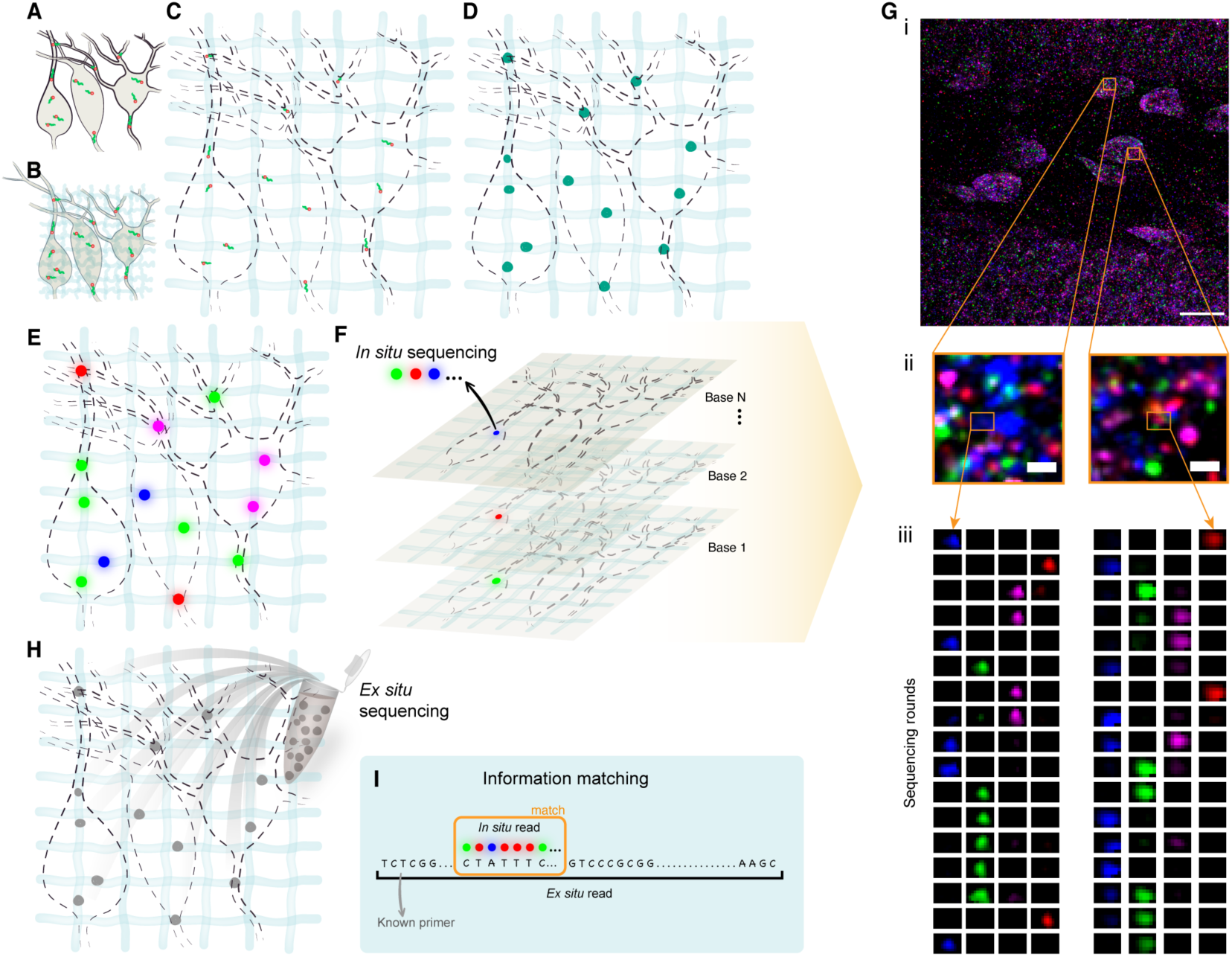
Untargeted expansion sequencing (ExSeq) concept and workflow. **(A)** A specimen is fixed, and RNA molecules (green) bound by LabelX anchor (orange). **(B)** The specimen is embedded in a swellable gel material (light blue, not to scale), mechanically softened using protease, and then expanded with water **(C)**. The RNA molecules are anchored to the swellable gel material. **(D)** RNA molecules present in the sample are reverse transcribed and amplified using FISSEQ, with the sequencing occurring via standard chemistries (e.g., SOLiD sequencing). **(E)**. Colored dots indicate the four colors used in SOLiD. **(F)** In each sequencing round different colors (blue, magenta, green, and red) reveal the current base of the amplified cDNA. **(G)** Example of ExSeq from a 50 micron thick slice of mouse hippocampus dentate gyrus. (i) One sequencing round, with two zoomed-in regions (ii), and puncta histories obtained over 17 rounds of *in situ* sequencing (iii). **(H)** After completion of *in situ* sequencing, cDNA amplicons are eluted from the sample, and resequenced *ex situ* with classical next-gen sequencing. **(I)** *In situ* reads are matched to their longer *ex situ* counterparts, with only unique matches being retained, augmenting the effective *in situ* read length. Scale bars: Gi, 17 microns (in biological, i.e. pre-expansion, units used throughout, unless otherwise indicated), Gii, 700 nanometers.

*In situ* sequencing has previously only been demonstrated with short reads (5-30 bases; (*10, 11, 20*)), due to laser-induced damage during imaging (*23*) and dependence of the signal for a given cycle on the signal from the previous cycle (known as “phasing”), caused by incomplete enzymatic reactions (*26*). This is undesirable, as alignment of short reads to the genome is extremely challenging (*27*). Moreover, short reads do not allow effective exploration of *in situ* mRNA complexity, as required, for example, for the study of alternative splicing. Accordingly, we developed a way to extend the *in situ* sequencing reads with a follow-on round of *ex situ* sequencing, using classical “next-gen” sequencing. We digest the expanded gel after *in situ* sequencing, extracting the amplified cDNA from the sample, and then use Illumina chemistry to re-sequence the same amplified cDNA molecules that were sequenced *in situ* (**Fig. 1H**).

Importantly, the random nature of untargeted reverse transcription priming, cDNA post processing that limits the size of the cDNA to ∼100 bases (see **Methods**), and the circularization of the cDNA, result in the creation of unique molecular identifiers in the *in situ* sequenced region of the amplified cDNA (**Fig. S5**); this allows us to use the *ex situ* information as a dictionary to align and directly interpret the *in situ* sequencing reads (**Fig. 1I**, **Fig. S5A** bottom panel). 92% of all matches and 97% of the 16,253 matches that aligned against non-rRNA were strictly unique in the sense that one *in situ* read matched precisely one annotated *ex situ* library entry (**Fig. S5C**). To prevent ambiguity, we removed the handful of *in situ* reads that matched to more than one annotated *ex situ* library entry (see **Methods** section ‘*Ex situ* and *in situ* sequence matching’); this allows us to explore sequence variations in mRNA, such as alternative splicing, using the longer *ex situ* matched reads. Leveraging this *ex situ* information enables a 1-2 order of magnitude increase in effective *in situ* read length compared previous *in situ* sequencing (**Fig. S6**).

We found that this expansion sequencing (ExSeq) chemistry could produce data from a variety of samples, including mouse brain (**Fig. 1G**), intact C. elegans **(Fig. S7A**), Drosophila embryos (**Fig. S7B**), and HeLa cells (**Fig. S7C**). To quantitatively validate the performance of ExSeq, we used the following mouse specimens: cultured hippocampal neurons (**Fig. 2A-B** and **Fig. S8**), a 15 micron thick hippocampal slice (**Fig. 2C-D**), and a 50 micron thick hippocampal slice (**Fig. 2E-F**); see primers and data for these experiments in **Tables S1-S4**, and summary statistics for the 50 micron thick hippocampal slice in **Table S5**. The length of the *ex situ* sequencing reads was 300 paired-end bases for the 15 micron thick hippocampal slice (using an Illumina MiSeq), and 150 paired-end bases for the two other samples (using an Illumina NextSeq); to improve the efficiency of cDNA circularization we restricted the size of the cDNA fragments to be ∼100 bases long, and as a result the Illumina reads typically contained several repeats of a given cDNA fragment, with average actual sequence length of 76 bases (**Fig. S6**).

**Fig. 2.**
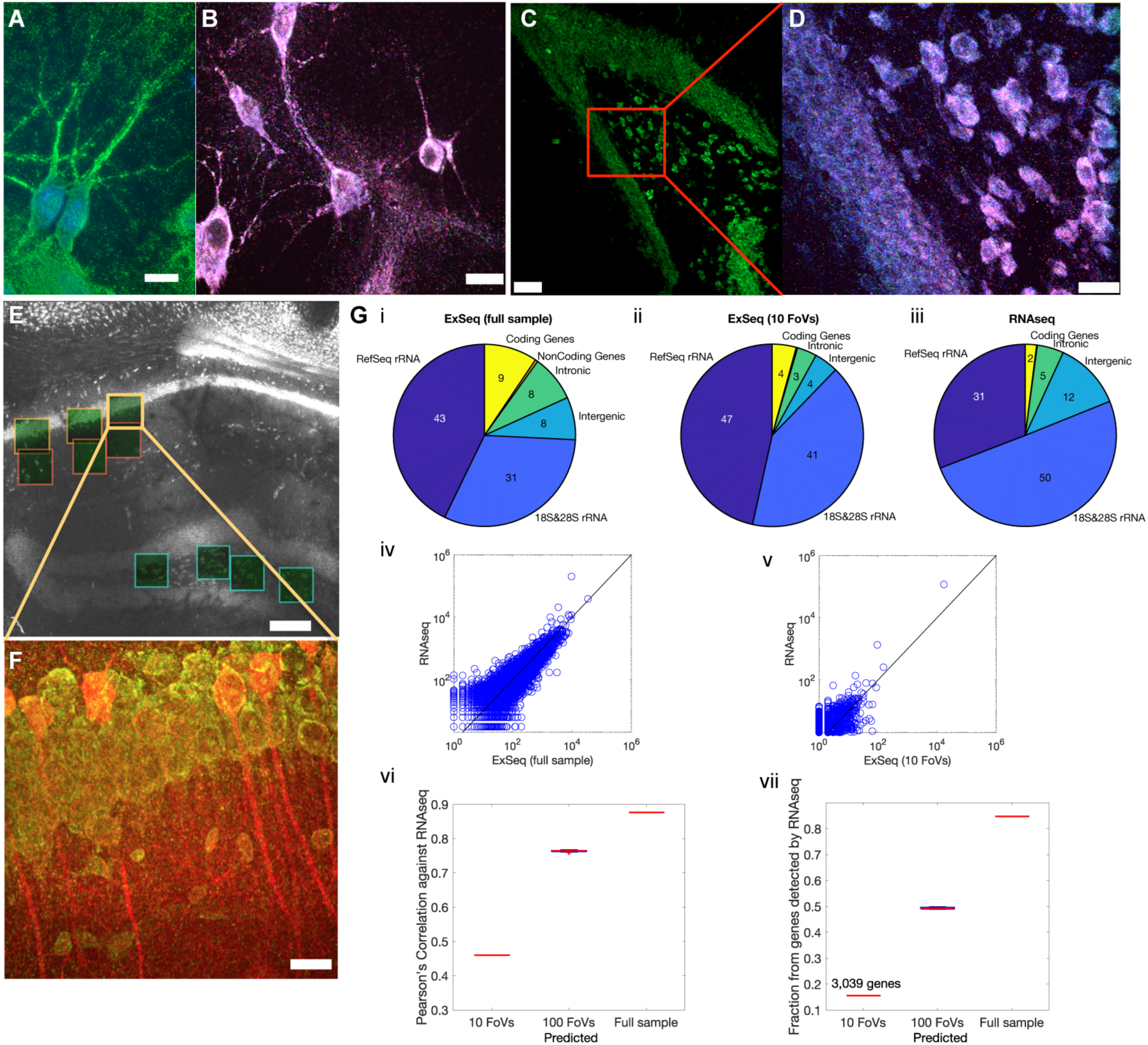
*In situ* sequencing in cells and tissues with untargeted ExSeq. **(A)** Example of ExSeq library preparation in hippocampal culture (green, hybridization probe against amplified cDNA; blue, DAPI). **(B)** Maximum intensity projection of one sequencing round in hippocampal culture; color scheme as in Fig. 1G. **(C)** Low magnification image of ExSeq library preparation in a 15 *μ*m slice of mouse hippocampus (green, hybridization probe against amplified cDNA). **(D)** Maximum intensity projection of a higher magnification image of the specimen in C, focusing on one sequencing round; color scheme as in Fig. 1G. **(E)** Low magnification image of ExSeq library preparation in a 50 *μ*m slice of mouse hippocampus. Fields of view (FoVs) acquired with a higher magnification objective are shown as green squares. White, hybridization probe against amplified cDNA. **(F)** Maximum intensity projection of one FoV of panel E, with antibody staining shown post *in situ* sequencing (red, antibody against YFP (the specimen is a Thy1-YFP mouse); green, hybridization probe against amplified cDNA). **(G)** Sequence analysis of ExSeq specimen shown in (E). (i-iii) The overall RNA content obtained with ExSeq, either with full *ex situ* sequencing data from the entire slice used (i) or using *ex situ* data reads that exactly correspond to *in situ* reads observed within the 10 indicated FoVs in panel E (ii), is in agreement with the overall RNA content of an adjacent slice obtained with standard RNAseq using random primers (iii). The numbers inside the pie chart represent the percentage of the total. (iv) Agreement between the normalized expression levels of all well-annotated genes (RefSeq genes) using RNAseq and ExSeq with full *ex situ* sequencing data as in i. (v) Same as (iv), but using the 10 acquired FoVs, as in ii. (vi) Pearson’s correlation between the log-transformed expression of RefSeq genes using ExSeq and using RNAseq, as a function of the number of acquired FoVs (estimated by sampling from the full *ex situ* sequencing data to simulate the number of expected reads for 100 FoVs; see **Methods** for details). The value for the 100 FoVs is plotted using the MATLAB boxplot function: on each box, the central mark indicates the median, and the bottom and top edges of the box indicate the 25th and 75th percentiles, respectively. (vii) Fraction of RefSeq genes detected using ExSeq compared to RNAseq, as a function of the number of acquired FoVs (estimated by sampling from the full *ex situ* sequencing data to simulate the number of expected reads for 100 FoVs). Scale bars: A-D&F, 13*μ*m; E, 130*μ*m. Note that deconvolution was used in panels D and F, see **Methods**.

The mouse specimens were prepared with two different fixation procedures (fresh frozen followed by slicing and paraformaldehyde (PFA) fixation, and transcardial PFA perfusion, for the 15 and 50 micron thick slices, respectively), and produced similar *in situ* amplified libraries. Antibody staining after *in situ* sequencing, as with previous ExM-related protocols (*28*), can enable visualization of specific proteins; staining with antibodies against YFP in a Thy1-YFP mouse line (*29*) allows for neurons in the hippocampus to be visualized in conjunction with *in situ* sequencing reads (**Fig. 2F**).

To gauge the accuracy of the results obtained with ExSeq, we performed standard RNA sequencing (RNAseq), with random primers, on a 50 micron thick hippocampal slice adjacent to the 50 micron thick ExSeq specimen displayed in **Fig. 2E-F**. As expected for total RNA analysis, most of the RNA detected with both ExSeq and RNAseq was ribosomal RNA, with overall agreement between the RNA types obtained with both methods (**Fig. 2Gi-iii**; albeit with a higher percentage of coding RNA, 4-9% with ExSeq vs 2% with RNAseq). Gene ontology analysis on the genes detected using both methods revealed the expected, and common, function enrichments for this brain region, including ‘synapse’, ‘neuron projection’ and ‘hippocampus’ (**Fig. S9** and **Table S6**). Notably, whereas in FISSEQ highly abundant genes, for example genes involved in translation and splicing, were underrepresented (*20*), we did not observe such a detection bias with ExSeq (see **Methods**). Accordingly, high correlation was observed between the expression levels of all well-annotated genes (RefSeq genes) using RNAseq and ExSeq (Pearson’s r = 0.89), using the full *ex situ* sequencing dataset of the rolling circle amplicon library produced *in situ* during the ExSeq library preparation process (**Fig. 2Giv** and **Fig. S8C**). The correlation was a function of the number of volumes acquired with the confocal microscope, with 10 volumes (each from a field of view of 350×350×100 microns in size, post-expansion) resulting in Pearson’s r = 0.47 (**Fig. 2Gv**; p-value 9 x 10^-164^), with the correlation increasing with the number of volumes analyzed (**Fig. 2Gvi**), estimated by simulating these larger datasets by sampling from the full *ex situ* sequencing data (see **Methods**). The correlation obtained, even with only 10 volumes, is at the high end of the values obtained using the original FISSEQ protocol, in which the Pearson’s r was between 0.2 to 0.5 in the different cell cultures studied. Moreover, the observed correlation is comparable to recent targeted *in situ* sequencing methods; for example, Svedlund et al. report Pearson’s r values ranging from 0.46 to 0.69 (*30*). With 10 volumes, 3,039 genes were detected comprising ∼16% of all the genes expressed in the sample, with the number of genes detected again increasing with the volume sampled (**Fig. 1Gvii**). Thus, ExSeq is able to report on genome-wide expression *in situ*, in an untargeted and highly multiplexed way.

We next sought to utilize the improved spatial resolution of ExSeq, enabled by the physical tissue expansion of ExM, and the ability to support antibody labeling, to pinpoint the locations of transcripts within neural architecture. We manually traced hippocampal CA1 pyramidal neurons, YFP-expressing and antibody labeled as described above (n = 13 hippocampal CA1 pyramidal neurons, across three volumes taken on a confocal microscope, from one mouse). Then, we analyzed the locations of individual *in situ* sequenced RNA transcripts inside the cell bodies and the dendrites of individual identified neurons, visualized with a custom 3D viewer (**Fig. 3** and **Fig. S10**, see **Methods**). The average number of sequencing reads per neuron was 229 ± 74 (mean ± standard deviation used throughout) including rRNA, and 30 ± 14 for non-rRNA, with the dendrites partly imaged, up to ∼100 microns from the cell body. Not including rRNA genes, 326 different RefSeq genes were observed in the imaged parts of these 13 neurons. These numbers are comparable to the original FISSEQ protocol (200-500 reads per cell (*23*)), including rRNA, while the latter was performed in cultured cell lines rather than intact brain tissue.

**Fig. 3.**
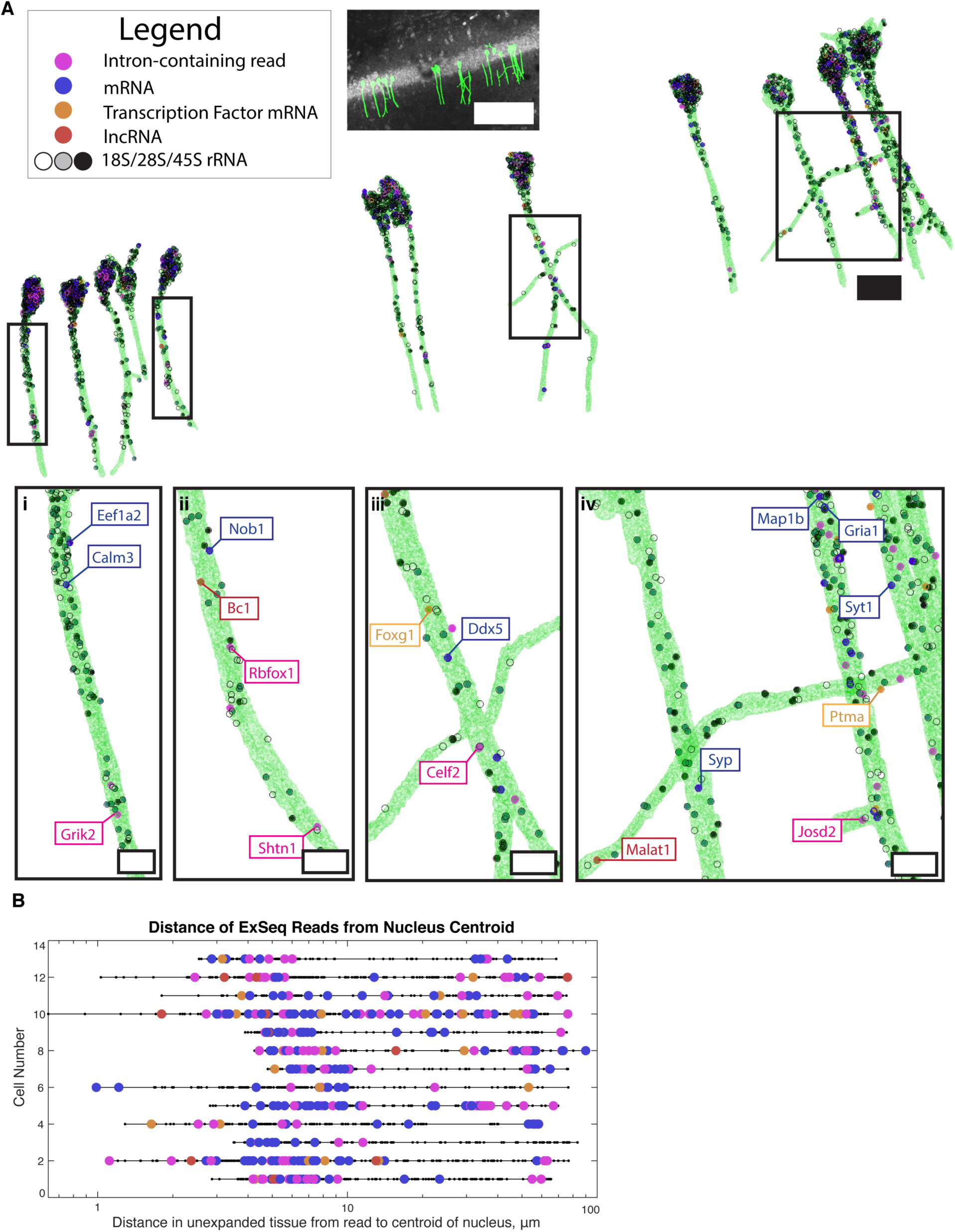
Untargeted ExSeq enables mapping of RNAs and their variants in dendrites of neurons. **(A)** 3D render of Thy1-YFP CA1 neuronal morphology as determined by YFP antibody staining, containing multiple RNA types (intron-containing non-transcription factor mRNAs; intron-free non-transcription factor mRNA; intron-free transcription factor mRNAs; lncRNAs; various rRNA sequences). The zoomed-in regions (boxed above, zoomed-in below) highlight RNA localization in dendrites. Scale bars: top, middle and bottom, 100, 20, and 5 microns, respectively. **(B)** Euclidean distance, relative to the center of the cell body, of sequencing reads for the neurons in A. Color code is as in A.

Splicing of RNA is known to play a role in synapse specification (*31*), and neurons contain one nucleus vs. thousands of potentially molecularly distinct synapses, raising the question of whether splicing of expressed mRNAs, including those that contribute to synaptic function, is regulated in a spatially dependent manner along dendritic trees (*32*). We examined reads that corresponded to intron-containing regions, and found that, while most (70%) of reads that mapped to intronic regions were located inside the soma (as is the case for most genes), we also observed introns in YFP-containing dendritic projections up to 100 microns away from the cell body layer, consistent with previous reports of cytoplasmic intron-retaining transcripts throughout neural processes (*33, 34*). For example, Glutamate ionotropic receptor kainate type subunit 2 (Grik2), which encodes a receptor subunit involved in excitatory glutamatergic neurotransmission that is implicated in autosomal recessive cognitive disability, appears in our data as a dendritic mRNA with a retained intron (**Fig. 3Ai**); the Grik1 subunit has been previously identified as a dendritically-targeted intron-retaining sequence (*33, 35*). This suggests that splicing of multiple glutamate receptor subunit RNAs may occur inside dendrites in intact hippocampal neural circuits, potentially contributing to a spatially regulated diversification of excitatory neurotransmission responses; note that splicing in dendrites was found and characterized previously in cultured neurons (*36*).

The long sequencing reads **(Fig. S6**) and untargeted nature of ExSeq also allows for direct observation of alternative splicing complexity and diversity *in situ.* We first quantified the expression of known alternative splicing isoforms with ExSeq compared to standard RNAseq, revealing a high correlation between the two methods (Pearson’s r = 0.944; **Fig. S11A**). Using only 10 fields of view taken on the confocal microscope, we detected 112 sequencing reads that correspond to known alternative splicing of cassette exon events; 67% of them revealed the identity of the expressed alternative splicing isoforms, for example the genes Ribosomal protein S24 (Rps24) and Microtubule-associated protein 2 (Map2) (**Fig. S11B**). In some cases the sequencing reads revealed possibly new isoforms, for example for the gene Spectrin Beta (Sptbn1) (**Fig. S11B**). Importantly, ExSeq provides the ability to locate these alternative splicing events in space; for example, isoforms of Map2, which is involved in determining and stabilizing dendritic shape (*37*), and the transcription factor Cux1, which is involved in dendrite branching and spine formation (*38*), could be precisely localized to the neuronal soma, outside of the nucleus, of their respective traced neurons (**Fig. S11B** and **Fig. S10**, Neurons 9 and 7, respectively). Thus, ExSeq may be useful in the future for mapping out the locations of splice variants throughout neural morphology in defined cell types in intact brain circuitry, for generating new hypotheses about the role of splicing in neuronal communication and plasticity.

Many genes with known or unknown functions may have previously unappreciated connections to neuronal signaling inside dendritic trees. Previous reports indicated the presence of mRNAs for specific transcription factors inside dendrites (*39*), for example MAX dimerization protein (Mga; (*40*); **Fig. S10**, Neuron 6), yet the full complement of dendritically localized transcription factors in any given neuron type is unknown. In the hippocampus sample, 914 of the known 1675 mouse transcription factors (RIKEN transcription factor database) were detected by ExSeq, 32 reads of which were localized within YFP-expressing cells in the Thy1-YFP mouse, including 11 reads in the dendrites of these cells. These reads include Forkhead box protein G1 (Foxg1), known to be involved with neural development (*41*), and prothymosin alpha (Ptma), involved in learning and memory and neurogenesis (*42*) (**Fig. 3Aiii** and **3** Aiv). We also found in dendrites long non-coding RNAs (lncRNAs) and protein coding genes with unknown function (**Fig. 3A**), raising the question of whether considering dendrite-specific roles may be relevant to understanding the functions of these gene products. For example, BC1 (**Fig. 3Aii**) is a ncRNA from an RNA polymerase III transcript that complexes with proteins to form a ribonucleoprotein particle, long known to be expressed almost uniquely in the brain and dendritically localized (*43*), and involved with activity-dependent synaptic regulation (*44*).

MALAT1, which we observed in a distal dendrite (**Fig. 3Aiv**), has been studied for its role in neural growth, polarization and synaptogenesis, but its localization has not been studied in intact hippocampal tissue (*45, 46*). We localized genes that were shown to be located in dendrites of CA1 pyramidal cells at the protein level, but which had not been mapped at the mRNA level such as Gamma-Aminobutyric Acid (GABA) type A receptor gamma2 subunit (Gabrg2; **Fig. S10**, Neuron 2) (*47*). Thus, ExSeq allows us to rapidly expand our knowledge of dendritically localized genes of known function, such as certain transcription factors, which may point to possible novel hypotheses for these gene products, e.g. synaptic activity dependent gene regulation; we can also identify transcripts encoding for genes of function unknown in the hippocampus (e.g, Nob1, **Fig. 3Aii**) (*48*), which may contribute to their functional analysis.

To more systematically understand how the types and identities of transcripts varied with location along a dendrite, we measured the distance from each read to the centroid of its corresponding neuron’s cell body (**Fig. 3B**), revealing the positions of RNAs encoding for transcription factors, intron-containing reads, and lncRNAs up to 100 microns from the soma.

This untargeted genome-wide analysis can be followed by an in-depth examination of specific genes through other technologies -- for example microtubule-associated protein 1B (Map1b), involved in microtubule assembly, and glutamate receptor 1 (Gria1) (**Fig. 3Aiv**), an excitatory neurotransmitter receptor, are further analyzed through our targeted ExSeq methodology across a complete section of the mouse hippocampus, later in this paper. Thus, ExSeq allows spatial and cellular subcompartment-based analysis of *in situ* sequencing reads in intact 3D brain tissue, which may be useful for generating new hypotheses about subcellular signaling relevant to brain circuit computation and disease.

Although untargeted sequencing enables transcriptome-wide exploration and discovery of localized RNAs, including rare variants and those of unknown function, the vast diversity of possible reads generated by untargeted methods naturally leads to a lower per-gene copy number of detected molecules, and a larger number of required biochemical and imaging cycles to distinguish among reads, compared with targeted methods that must only distinguish a smaller predefined set of genes which can be indexed by short barcodes. Targeted methods are applicable to key questions such as mapping cell types and states, and their spatial relationships *in situ*, and visualizing subcellular gene regulation of known sequences. For the targeted single-molecule multiplexed interrogation of RNA, an ideal technology would satisfy the following list of criteria. First, the method should possess sufficient yield (probability of detecting a present molecule) in order to detect low copy number transcripts such as transcription factors or sparse RNA molecules in subcellular compartments. Second, the technology should have resolution below the diffraction limit both laterally and axially to resolve nanoscale morphological features, such as dendritic spines in neurons or close 3D appositions of individual cells in dense tissues.

Third, to take advantage of the improved resolution, the method should provide the multimodal ability to image both RNAs and proteins, and to work with 3D tissues (e.g., multiple cell layers), in order to localize RNAs in their native biological context. Finally, the method should work with various tissue types, including human tissues. Given the ability of expansion microscopy to facilitate high resolution imaging and multimodal imaging, we thus decided to build from the untargeted version to develop a targeted version of ExSeq (performance summarized in **Tables S7-S8**).

In targeted ExSeq, oligonucleotide padlock probes bearing barcodes are used to directly hybridize to specific transcripts (*11, 49*), and generate amplicons for subsequent readout through *in situ* sequencing of the barcodes, the latter performed in a manner similar to that of untargeted ExSeq (**Fig. 4A**, **Fig. S12)**. The requirement for the inefficient (*20*) reverse transcription step of most *in situ* sequencing technologies (*11, 20, 50*) is circumvented by the direct binding and ligation of multiple padlock probes on each targeted transcript using PBCV-1 DNA Ligase (also known as SplintR ligase), which can ligate DNA on an RNA template a hundred-fold faster than the commonly used enzyme for ligation, T4 DNA Ligase (*49, 51–54*). After circularization and rolling circle amplification, the barcodes are sequenced *in situ*. As the barcodes are sequenced across multiple rounds of imaging, the number of resolvable molecular targets scales exponentially with the number of imaging rounds. We can use both Illumina sequencing-by-synthesis and SOLiD sequencing-by-ligation, and expect other sequencing chemistries to be usable as well (*10, 11, 50*). We thus sought to explore the performance of the method, and to apply it to multiple contexts in neuroscience and cancer biology, including resolving RNA amidst cellular morphology and mapping spatially dependent cell relationships in a cancer sample (summarized in **Table S5**).

**Fig. 4.**
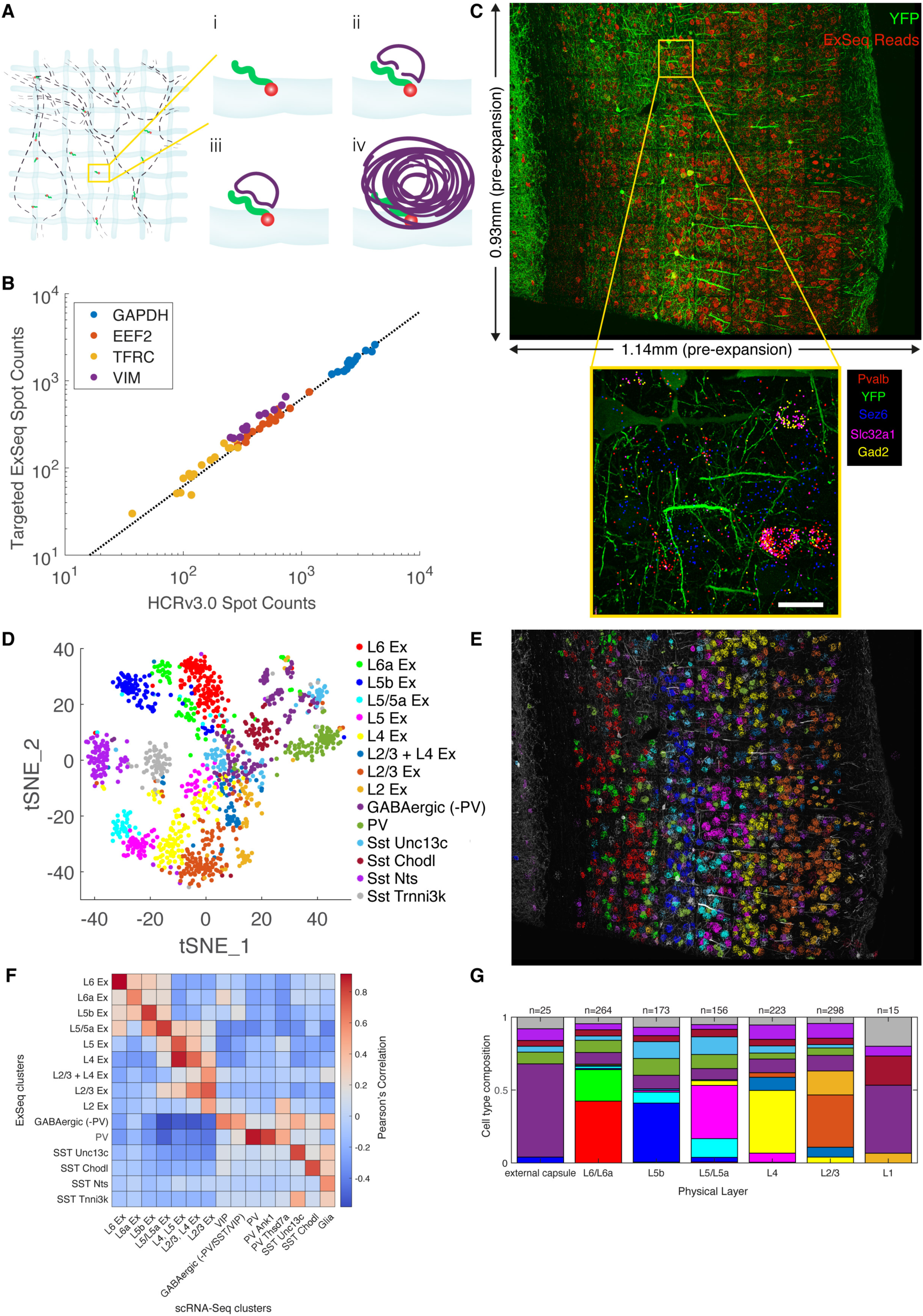
Targeted ExSeq efficiently detects transcripts, recapitulating the organization of neuron types of mouse primary visual cortex. **(A)** Targeted ExSeq library preparation: (i) RNA anchoring and expansion; (ii) padlock probe hybridization; (iii) probe ligation; (iv) rolling circle amplification. **(B)** Amplicon counts for targeted ExSeq vs. HCRv3.0-amplified ExFISH for the same transcript in the same HeLa cells (60 cells in total); slope is 0.62 (Pearson’s r = 0.991). **(C)** Targeted ExSeq of 42 cell type marker genes in Thy1-YFP mouse visual cortex. Top, maximum intensity projection image showing all targeted ExSeq reads (red) and YFP protein (green). Bottom, localization of canonical marker genes Pvalb (red), Sez6 (cyan), Slc32a1 (magenta), and Gad2 (yellow), with YFP (green). **(D)** Targeted ExSeq gene expression profiles of 1154 cells clustered into 15 cell types. Cluster legend and colors apply to panels D, E, and G. **(E)** Spatial organization of cell types identified in (D). Cell-segmented reads are shown, colored by cluster assignment, and overlaid on the YFP morphological marker (white) indicating a subset of neurons of the primary visual cortex. **(F)** Heatmap showing Pearson’s correlation between clusters identified in the targeted ExSeq dataset and a prior scRNA-seq study (*58*). **(G)** Layer-by-layer fractional cell-type composition across segmented cortical layers. Scale bars: (C) bottom, 20 microns (pre-expansion).

To validate the yield of the method, hybridization chain reaction (HCR)v3.0-amplified expansion FISH (ExFISH) and targeted ExSeq were sequentially performed on the same expanded HeLa cells, for the same genes (*24, 55*). The number of ExFISH puncta vs. targeted ExSeq amplicons were quantified (**Fig. 4B**, barcodes and probes are in **Tables S9-S10**, and raw data in **Table S11)**, indicating that targeted ExSeq has an RNA amplification and detection yield of ∼62% (Pearson r = 0.991) relative to HCRv3.0-ExFISH, which itself has a detection efficiency of ∼70% in tissue (*24*). These numbers are remarkably high considering that single-cell RNA sequencing captures about 10% of mRNA, and the probability of recovery of any individual transcript is low (*56, 57*).

We then assessed the performance of targeted ExSeq by mapping the cell types of a tissue with existing single cell RNA sequencing (scRNA-Seq) data-based classification of cell types, the mouse primary visual cortex. Based on scRNA-Seq data classifying the cell types of the visual cortex (*58*), we designed a panel of probes targeting 42 genes (sequences in **Tables S9-S10**) that mark key excitatory and inhibitory neuronal cell types. We performed targeted ExSeq of these 42 genes across a coronal section of the entire primary visual cortex of a Thy1-YFP mouse over a volume of 0.933 mm x 1.140 mm x 0.02 mm, sequencing 265,347 reads throughout this volume (**Fig. 4C**, top; counts in **Table S12**). The spatial distribution of ExSeq reads for known marker genes recapitulated the spatial distributions in the Allen *in situ* hybridization (ISH) atlas (**Fig. S13**). Transcripts known to be co-expressed in the same cell type also appeared in similar relative positions -- for example, in one field of view, the Parvalbumin (Pvalb), Vesicular inhibitory amino acid transporter (Slc32a1), and Glutamate decarboxylase 2 (Gad2) transcripts, all known to be co-expressed in parvalbumin-positive (Pvalb^+^) interneurons (PV interneurons), were co-located within the same cells while Seizure protein 6 homolog (Sez6) transcripts, associated with excitatory neurons in deeper cortical layers (as well as vasoactive intestinal peptide (VIP^+^) interneurons) did not co-localize with the the Pvalb, Slc32a1, and Gad2 transcripts (**Fig. 4C**, inset).

To ascribe reads to cells, we developed a cell segmentation pipeline (see **Methods**, **Fig. S14**) that yielded a total of 1915 cells containing a total of 220,783 reads, out of which 1154 cells with at least 50 reads (177 ± 127 reads/cell) were used for subsequent analysis. To identify cell types in targeted ExSeq data, we clustered cell expression profiles using *k*-means and subsequently embedded them into a low-dimensional space using t-Stochastic Neighbor Embedding (t-SNE) (*59*) (**Fig. 4D**). Two broad types of clusters were identified using previously known markers (see **Methods**), corresponding to excitatory neurons (“Ex”, and sub-annotated by their layer location) and inhibitory neurons (annotated with relevant cell type marker). The expression of key marker genes in detected clusters (shown in **Fig. S15**) was consistent with prior work (*58*).

We compared our results to those from a previous study in which scRNA-Seq was performed on mouse primary visual cortex (*58*). We clustered the previous single-cell dataset, focusing on the 42 genes from our gene panel, with *k*-means clustering, and compared ExSeq clusters to scRNA-Seq clusters, finding good agreement between the clusters (**Fig. 4F**). When we projected the locations of the ExSeq-derived cell types onto the visual cortex image, we observed the canonical layer-by-layer stratification of excitatory neurons in the visual cortex, with 9 ExSeq clusters mapping onto classical layers 2 through 6 of the cortex (**Fig. 4E**, **Fig. S16**); these 9 clusters corresponded, with slightly different grouping perhaps due to how the clustering algorithm treated subtleties in the statistical properties of each dataset, to 7 scRNA-Seq clusters of excitatory neurons (**Fig. 4F**) -- for example, the ExSeq cluster L4 Ex mapped primarily onto two scRNA-Seq clusters, L4, L5 Ex and L2/3, L4 Ex. Other ExSeq clusters of excitatory neurons corresponded to mixtures of multiple scRNA-Seq clusters as well, often at the boundaries between cortical layers, which may represent real cell-type biological variability (**Fig. 4E**).

Among inhibitory neurons, we found some cell types to match in 1-to-1 correspondence between ExSeq and scRNA-Seq clusters -- for example, two somatostatin interneuron clusters, the SST cluster expressing Unc-13 homolog C (cluster SST Unc13c) and the SST cluster expressing Chondrolectin (cluster SST Chodl) appeared prominently in both datasets; both sets of neurons were scattered throughout cortical layers L1-L6. As with the excitatory neurons, some ExSeq clusters of inhibitory neurons mapped onto multiple scRNA-Seq clusters. For example, two ExSeq clusters, which we denoted PV and GABAergic (-PV), mapped onto multiple scRNA-Seq clusters, the former onto three scRNA-Seq cluster that corresponded to subclasses of PV cells, and the latter onto VIP neurons and non-PV/VIP/SST GABAergic neurons, with some cells falling into other clusters.

Such poolings of scRNA-Seq clusters into ExSeq clusters (and vice versa) are likely due to the smaller number of cells analyzed with ExSeq vs. scRNA-seq, the small number of markers interrogated, and the use of a simple *k*-means algorithm for clustering. Some substructure is visible in the t-SNE plot for the cluster GABAergic (-PV) (**Fig. 4D**), for example, suggesting that alternative clustering approaches, for instance utilizing additional morphological criteria or protein markers, or more sophisticated segmentation algorithms that take into account priors on the transcriptome from scRNA-Seq, could be devised in the future to yield more precise delineations of cell types. We tested the robustness of the approach by varying the parameters for cell segmentation of the ExSeq dataset, and by varying the genes used for clustering the single-cell dataset, and found our conclusions to be robust to these algorithmic choices (**Fig. S17-S18**). As non-neuronal cells (including microglia, astrocytes, and oligodendrocytes) did not highly express the interrogated markers, they were inefficiently detected, and were non-specifically clustered with diverse interneuron types. For cell types whose markers were well represented, targeted ExSeq was able to classify cells with high agreement with established non-spatial methods for cell-type classification, with the benefit of providing information on the spatial organization of the cells.

As described previously (*58, 60*), the canonical layer-specific excitatory neuron transcription factor marker genes homeobox protein cut-like 2 (Cux2), RAR-related orphan receptor beta (Rorb), Fasciculation and elongation protein zeta-2 (Fezf2) and Forkhead box protein P2 (Foxp2) were expressed in L2/3, L4, L5b, and L6 respectively (**Fig. 4E-F**, **Fig. S15**). The clusters corresponded to excitatory cells localized to a given layer, and were used to segment the cortex into layers (**Fig. S14D**), so that the cell types within each layer could be quantified (**Fig. 4G**; raw counts, **Fig. S19**). Each of the clusters of inhibitory neurons was found to be dispersed across layers L2 to L6 (**Fig. 4G**, **Fig. S19**), consistent with prior work in which each of the canonical GABAergic cell types (PV, VIP, SST) were found to exhibit a fairly even layer distribution (*58, 60*). Thus, targeted ExSeq enables sensitive RNA detection across circuit-relevant volumes of intact tissue, enabling cell types to be systematically mapped while preserving their spatial context.

We next used the high spatial resolution of targeted ExSeq to explore nanoscale RNA compartmentalization within neurons of the mouse hippocampus, where local RNA translation and the dendritic locations of specific RNAs are implicated in mechanisms of synaptic plasticity and learning (*61–63*). We applied targeted ExSeq to Thy1-YFP mouse hippocampal brain circuitry, tracing the YFP signal to identify dendrites and spines, and targeting for sequencing 34 transcripts previously found in dendrites of CA1 neurons (*64*). We note that spines could not be observed in the untargeted hippocampus data because the antibody staining was performed post-sequencing, resulting in a weaker staining, whereas here the antibody staining was performed pre-expansion (see **Methods**). We performed four rounds of *in situ* sequencing to localize these transcripts on 170 fields of view (1.7mm x 1mm x 0.02mm, **Table S5**) spanning a coronal section containing the major subfields of the hippocampus, yielding 1.2 million reads, 90,000 of which localized within YFP expressing neurons (**Fig. 5A**, counts in **Table S13**). The overall cellular distributions of expressed genes were similar across the hippocampus to the patterns reported in the Allen Brain Atlas *in situ* hybridization dataset (**Fig. S20**). Using the YFP signal as a mask, we segmented the cell bodies, dendrites, axons, and spines of CA1 Pyramidal neurons, as well as the dendrites and cell bodies of dentate gyrus granule cells (the spines could not be automatically segmented because of low signal-to-noise), to localize transcripts within these subcellular and nanoscale compartments. We found transcripts within dendrites (CA1, DG), spines (CA1), and to a much smaller extent, in axons (CA1) (**Fig. 5B**). Out of 106,000 spines examined, we found 730 reads contained inside dendritic spines (one RNA per spine, except for one spine that had two). By simulating possible sources of bias **(Fig. S21)**, we concluded that our technique was not preventing detection of multiple reads per spine.

**Fig. 5.**
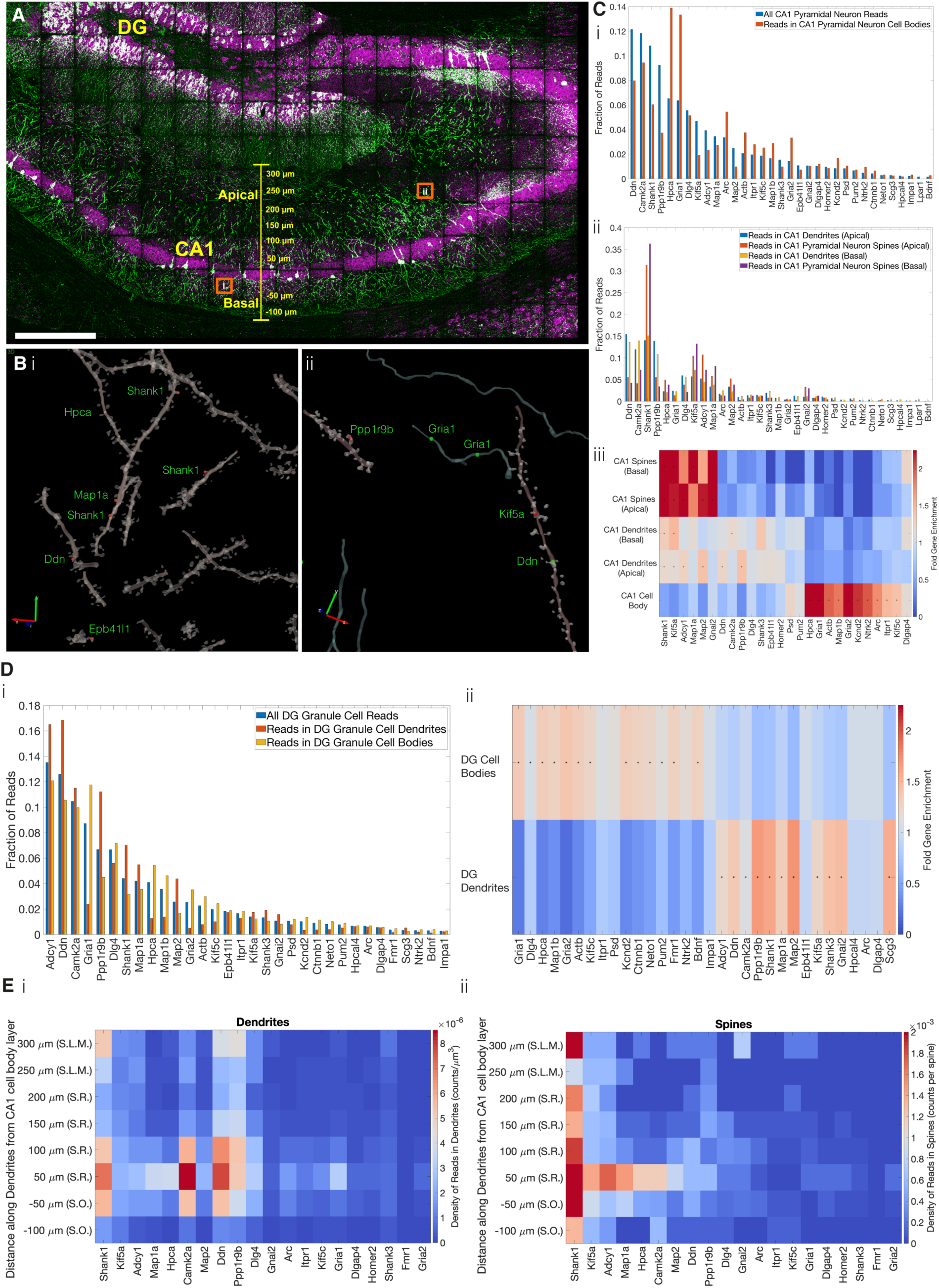
Targeted ExSeq characterization of nanoscale transcriptomic compartmentalization in mouse hippocampal neuron dendrites and spines. **(A)** Confocal image showing targeted ExSeq of a 34-panel gene set across a slice of mouse hippocampus. Green, YFP expressing cells; magenta, reads identified via ExSeq; white, reads localized within YFP expressing cells. DG, dentate gyrus; CA1, CA1 region of hippocampus. **(B)** 3D reconstruction of dendrites, spines, and axons showing reads localized in spines (red dots) and processes (green dots) for regions i and ii indicated by orange boxes in A. **(C)** The abundance and enrichment of transcripts in cellular compartments of CA1 Pyramidal Neurons: (i) abundance of transcripts in all cellular compartments vs. cell bodies; (ii) abundance of transcripts in apical and basal dendrites and spines; (iii) heat map showing the enrichment of transcripts in the different apical and basal dendritic and spine compartments of CA1 pyramidal neurons, vs. cell bodies; (*) indicates statistically significant enrichment (bootstrapped p-value <0.001). **(D)** The abundance and enrichment of transcripts in cellular compartments of dentate gyrus (DG) granule cells; (i) abundance of transcripts in the cell bodies and dendrites of DG granule cells; (ii) heat map showing the enrichment of transcripts in the compartments of DG granule cells; (*) indicates statistically significant enrichment (bootstrapped P-value <0.001). **(E)** Plots showing the density of transcripts in the dendrites (i) and spines (ii) of CA1 Pyramidal Neurons along the apical-basal axis (Euclidean distance) of hippocampal area CA1, including the CA1 regions S.R. (stratum radiatum), S.O. (stratum oriens), and S.L.M. (stratum lacunosum moleculare). Scale bars: A, 300 microns, Bi and Bii, 2 and 3 microns respectively, red and green arrows (pre-expansion).

We examined the distribution of reads across CA1 neuronal compartments including cell bodies, dendrites, and dendritic spines. As expected, genes such as the postsynaptic density protein Dendrin, the synaptic plasticity-associated gene Camk2a, and the postsynaptic scaffolding protein SH3 and multiple ankyrin repeat domains 1 (Shank1) were prominent among reads found in dendrites, whereas cell bodies exhibited a different distribution, with the neuronal calcium sensor Hpca and the synaptic glutamate receptor Gria1 being amongst the most abundant (**Fig. 5C**). In spines, we found Shank1, Adenylyl cyclase 1 (Adcy1) and kinesin family member 5a (Kif5a), a motor protein whose loss-of-function mutations may contribute to amyotrophic lateral sclerosis (ALS) (*65*), to be amongst the most abundant transcripts. When we compared the distribution of reads between CA1 pyramidal cell bodies, apical dendrites, basal dendrites, apical dendritic spines, and basal dendritic spines, we found that the distribution of reads in each compartment was statistically different from all others (bootstrapped 2-sample Kolomogorov-Smirnov test p-value < 0.001), except for apical versus basal spines which were not statistically different from each other (**Fig. 5Ciii**) -- suggestive of a common set of spine RNAs and spine RNA trafficking principles, at different sites throughout these neurons. To validate the accuracy of our targeted *in situ* sequencing results in this tissue context, we performed bulk RNA sequencing from hippocampal slices adjacent (+/-100 µm coronally) to the section we used for targeted ExSeq, finding a high level of correlation between our *in situ* sequencing results and the bulk RNA sequencing results (Pearson’s r = 0.85, **Fig. S22**).

Importantly, for the genes studied through both untargeted and targeted forms of ExSeq, we found high correlation between the readcounts (Pearson’s r = 0.68, **Fig. S22, Table S13**). Using the genes studied through both forms of ExSeq, we estimated the yield of untargeted ExSeq to be 0.6% compared to targeted ExSeq (**Table S13**).

Certain genes were significantly (bootstrapped p-value <0.001) enriched in specific CA1 neuronal compartments (**Fig. 5C**). In particular, spines and cell bodies of CA1 pyramidal neurons, and to a smaller extent their dendrites, showed distinct and contrasting patterns in the level of gene enrichment. The transcripts for Shank1, Kif5a, Adcy1, Map1a, Map2, and Gnai2, were highly enriched in spines, and these same transcripts were enriched to a smaller extent in both apical and basal dendrites, compared to cell bodies, perhaps pointing to a process through which these transcripts are progressively enriched the closer they get to synapses. Most of these genes serve structural roles in spines and dendrites, with Shank1 being a primary scaffolding protein in the postsynaptic density and Kif5a, Map1a, and Map2 interacting with the cytoskeleton (*66–68*). On the other hand, a distinct set of genes, including Hpca, Gria1, ActB, and Map1b, among others, were highly enriched in cell bodies as compared to dendrites or spines, as had been observed previously in the study that characterized the dendritic existence of these RNAs (*64*). Interestingly, Arc, a key mediator protein of LTP and LTD in synapses, was among the genes uniquely enriched in cell bodies, though this observation is consistent with the highly regulated, activity-related nature of its presence in dendrites (*69, 70*). In addition, a few genes such as Camk2a and Ddn were distinctly enriched in dendrites, consistent with previous studies (*64*), when compared to both spines and cell bodies (*64*). To assess whether these selective enrichments were cell type specific, we examined the enrichment profile of these same transcripts in the dendrites vs. cell bodies of YFP-expressing dentate gyrus granule cells (**Fig. 5D**). In dentate gyrus dendrites, we found transcripts similar to those found in CA1 apical and basal dendrites, such as Shank1, Map2, and Pppr1r9b, and across the entire 34 gene set, a striking similarity between the dendritic localization of RNAs in dentate gyrus granule cells vs. CA1 pyramidal neurons (r = 0.91, **Fig. S23)**. This similarity raises the possibility that there may be general rules governing the selection of specific RNAs for dendritic transport, or for their targeted trafficking to dendrites, and may motivate further investigations of the rules governing dendrite specific mRNA transport across cell types of the brain.

Seeing that specific RNAs were enriched in the spines of CA1 dendrites, we next asked how these spine-localized transcripts were distributed along the lengths of dendrites. Transcripts are known to exhibit varied distributions along dendrites, with some restricted to proximal dendrites and others showing localization distally (*64*). However, it is not known which spatial distributions occur for transcripts that are localized inside spines, because of the high resolution required to map this out in the circuitry of a brain slice. When we examined the density of transcripts within dendrites, we found that most were found close (+/-50 µm) to the cell body layer, and their density decayed rapidly towards distal regions of dendrites, similar to previous observations for these genes (*64*) (**Fig. 5Ei, Fig. S24**). Some transcripts, such as Shank1, Ddn, and Ppp1r9b were present in distal regions of dendrites. When we next quantified the presence of transcripts within spines along dendrites, however, we observed a markedly different distribution (**Fig. 5Eii, Fig. S24**). Most prominently, spine-localized Shank1 transcripts exhibited a strong presence throughput spines in both proximal and distal regions of dendrites in both apical and basal directions. The highest density of Shank1 in spines was found close to the cell body layer, but with a second highest density hundreds of microns away, in the stratum lacunosum-moleculare (SLM). Other transcripts, such as Kif5a and Adcy1 were found in spines of distal dendrites, but not at such a high density as Shank1. For most transcripts found in spines among those in our probe set, their highest density occurred close to the cell body layer. Thus, although spines are directly connected to dendritic branches, they can exhibit strikingly different distributions of mRNAs, here revealed by the high spatial resolution and multiplexing capability of targeted ExSeq across length scales relevant to neural circuitry.

High-resolution spatial transcriptome mapping has clear utility in neuroscience, as described above. We next asked if cancer biology and immunology might be able to utilize such techniques as well, and whether such techniques can be used in human tissues such as biopsies. One key question, for example, is to understand how tumor microenvironments, such as the state of local immune cells, govern tumor initiation, progression, metastasis, and treatment resistance. Multiplexed spatial mapping of RNA performed in human tissues to date has not achieved high enough resolution for single cell quantification, let alone subcellular resolution.

For example, Svedlund et al. (*30*) analyze the spatial sequencing data of human breast cancer tissues in 100-500 µm bins, due to low read counts. Single cell observation is also not possible in Spatial Transcriptomics (ST) and Slide-Seq (*71, 72*) (**Table S7**).

A core biopsy was taken from a patient with metastatic breast cancer infiltration into the liver, and 297 tumor-related genes of interest (selected based on single-cell and single-nucleus RNA-seq data, see **Methods**) were profiled in the intact tissue, resolving 1.15 million reads, including 771,904 reads in 2,395 DAPI-segmented cells (**Fig. 6A**, counts in **Table S14**). In addition, the nanoscale and 3D spatial resolution of ExSeq allowed the detection of 516 RNA reads inside nuclear structures that were under one micron in size, that are possibly nucleoplasmic bridges, a challenging structure to resolve in intact tissue (*73*) (**Fig. S25**).

**Fig. 6.**
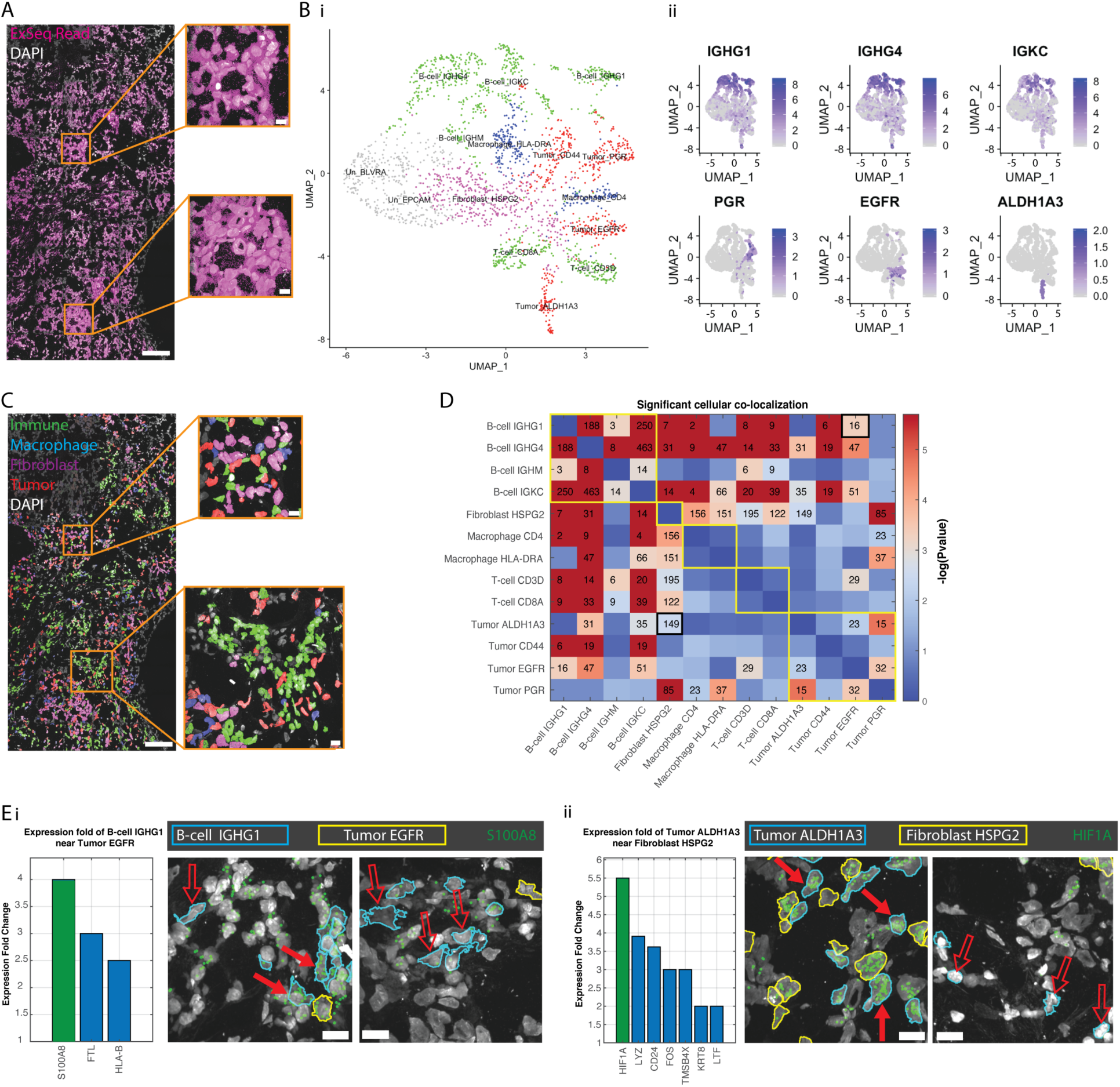
Targeted ExSeq resolves spatial maps of cell types and cell states in a metastatic breast cancer biopsy. **(A)** ExSeq resolves 771,904 reads in 2,395 cells (with at least 100 reads per cell) across 297 genes in metastatic breast cancer. **(B)** Uniform manifold approximation and projection (UMAP) representation of PCA-based expression clustering reveals immune and tumor cell clusters, indicated by different colors: green (T-cells, B-cells), red (tumor cells), blue (macrophage), magenta (fibroblast) and gray (un-annotated clusters, see **Methods**) (i), which express known cell markers for immune cells (ii, top row) and tumor cells (ii, bottom row), expression projected onto UMAP as log_2_(1+counts). **(C)** Transcriptionally-defined cell clusters mapped onto the tissue context, demonstrating regions containing mixed populations of fibroblasts, tumor cells, and immune cells (T-cells and B-cells, same colors as in B(i)) (top) and one region with a low density of tumor cells and high density of immune cells (bottom). **(D)** Spatial colocalization analysis of cell clusters. Adjacency matrix text values, number of cell pairs of indicated type that are in close proximity (nucleus centroid distance of <20 µm; robustness analysis in **Fig. S26**). Adjacency matrix heatmap, p-value (500,000 bootstrapping iterations) relative to obtaining the same or higher number of cells in close proximity by chance. Adjacency matrix entries with text values are statistically significant (Bejamini Hochberg false-discovery rate of 1.5%). Yellow borders along the diagonal illustrate major cell type categories (B-cell, Fibroblast, Macrophage, T-cell, Tumor), and two black-bordered entries correspond to the pairs shown in (E). **(E)** ExSeq resolves cell states that are a function of physical proximity, measured by calculating differential expression when cells of different kinds are spatially adjacent (<20µm) vs. farther apart. The gene with the largest fold change in a specific cell type when adjacent versus non-adjacent to another specific cell type is shown in green in the histogram (p-value = 1e-4 using 100,000 bootstrapping iterations, all other genes shown in the histogram have p-value < 0.05), as well as in the image showing the gene’s read locations in the original sample. Examples show (i) expression of S100A8 is elevated in B cells of a specific state (IGHG1-positive) when they are in close proximity to a specific kind of tumor cell (EGFR-positive), and (ii) expression of hypoxia-inducible factor 1 alpha (HIF1A) is inducted in tumor cells of a specific kind (ALDH1A3-positive) when in proximity to fibroblasts. Solid red arrow, cells in close proximity; hollow arrow, cells not in close proximity. In (i) the B cells and tumor cells are shown with blue and yellow boundaries, respectively, and in (ii) tumor and fibroblast cells are shown with blue and yellow boundaries, respectively; the boundaries of other cell types are not colored. Scale bar: (A) and (C) 100 microns, and 10 microns for the insets, (E) 10 microns (pre-expansion).

Expression clustering of the DAPI-segmented cells (using the R package Seurat (*74*), as we reasoned that a PCA-based approach would be appropriate due to the high number of targeted genes in this experiment) revealed the expected mixture of cell types in breast cancer metastasis, including tumor, immune (T-cell, B-cell, and macrophage), and fibroblast cell clusters, as characterized using known biomarkers such as members of the immunoglobulin family (IGHG1, IGHG4, IGKC) found in B-cells, and genes known to be expressed in metastatic breast cancer (Progesterone receptor, PGR (*75*); Epidermal growth factor receptor, EGFR (*76*); Aldehyde dehydrogenase 1 family member A3, ALDH1A3 (*77*)) (**Fig. 6B**). Displaying the resulting cell clusters in the tissue context revealed high levels of mixing between tumor and non-tumor cells (**Fig. 6C**) and allowed analysis of spatial colocalizations (here defined as proximity within 20 microns) between putative cell types (**Fig. 6D**), which were robust across various choices of distance parameter **(Fig. S26**). Specifically, we found that the different B-cell clusters tended to co-localize in space, consistent with previous observations (*78*). Interestingly, we found statistically significant co-localizations between most of the B-cell clusters and all the other cell clusters, including T-cell, macrophage, fibroblast and tumor clusters (**Fig. 6D**), except for one tumor cluster which specifically expresses the gene marker PGR (Tumor PGR). This is consistent with recent findings showing that B-cells directly interact with tumor cells and macrophages, and that physical interactions of B-cells with T-cells are important for humoral responses in the microenvironment (*79*). For example, Garaud et al. analyzed a region of breast cancer aggregates and identified 41% and 29% of B-cells touching tumor cells and T-cells, respectively (*78*). Our analysis also indicates other non-random co-localizations between the different clusters, for example between the fibroblast clusters and the macrophage, T-cell and tumor clusters (**Fig. 6D**). This is consistent with the role of fibroblasts in supporting leukocyte migration and aggregation at sites of cancer (*80*), and the suggestion that different fibroblast cell types have distinct spatial locations in breast cancer (*81*).

We next analyzed whether there are cell clusters that express genes differently as a function of physical proximity to other cell clusters. These might represent cellular states which are triggered by a physical contact or close proximity to a different cell cluster, or alternatively, cell states that induce such proximity to occur. For each pair of cell clusters that had non-random colocalization, we designated the cells in each cluster as being in close (<20 µm) vs. not close proximity, to cells of the other cluster, and then searching for statistically significant gene expression differences between the two groups (using bootstrapping, see **Methods**). For example, we found that hypoxia-inducible factor (HIF1A) was more than 5-fold overexpressed in one tumor cell cluster whenever these cells were in close proximity to a fibroblast cell cluster (**Fig. 6Eii)**. This is of particular interest as the protein level of HIF1A serves as a proxy of hypoxic environments, and has been previously described as an essential micro-environmental cue for tumor cell maintenance (*82*). Furthermore, the mRNA level of HIF1A might also be indicative of resistance to radiotherapy in tumors (*83*). We also found that the gene S100A8, an important regulator of inflammatory processes and immune response which was recently suggested as a biomarker for relapse and progression in breast cancer patients (*84–86*), was 4-fold overexpressed in a B-cell cluster whenever the cells were close to a tumor cell cluster (**Fig. 6Ei**).

In summary, ExSeq enables subcellular RNA mapping of 3D biological systems and can be applied to specimens of multiple organ systems and species, including human specimens. Beyond spatial genomics, we expect ExSeq to be useful for *in situ* sequencing of lineage (*87*) and/or connectome (*88–90*) indexing RNA barcodes, which incorporate designed or randomized base-level variation that is not naturally addressed by a FISH approach with a fixed set of tags and targets. More generally, the approaches for re-embedding, passivation, many-round sequential probing, image analysis and *ex situ* sequence matching in expanded samples that we have developed for ExSeq should be broadly applicable to other kinds of *in situ* enzymatic readouts, such as for the multiplexed readout of endogenous DNA or of antibody-attached tags, which may benefit from nanoscale spatial resolution in intact tissues.

## Acknowledgments

We acknowledge Allen Lin for the help with Illumina sequencing, Demian Park for providing cultured neurons, Evan Murray for providing cultured HeLa cells, Kiryl Piatkevich for performing transcardial perfusions, Reza Kalhor for helpful discussions, and Eftychios A. Pnevmatikakis for helpful discussions on image processing. We also acknowledge the SpaceTx analysis working group, for help in clustering: Trygve Bakken, Zizhen Yao, Peter Kharchenko, and in gene selection: Eeshit Dhaval Vaishnav, Brian Aevermann, Richard Scheuermann, Kenneth Harris. **Funding:** R.E.K. and A.H.M. acknowledge IARPA MICrONS (D16PC0008); E.R.D. acknowledges government support under HG005550 and HG008525 awarded by the National Institutes of Health and under DGE1144152 awarded by the National Science Foundation; S.A. is a HHMI fellow of the LSRF; D.G. is NSF GRFP fellow; F.C. is supported by the Schmidt Fellows Program at the Broad Institute; A.T.W. was supported by the Hertz Foundation Fellowship and the Siebel Scholarship; A.S. was supported by the NIH Neuroimaging Training Program T32 grant 5T32EB001680; J.M. and E.L. thank the Chan Zuckerberg Initiative, an advised fund of Silicon Valley Community Foundation for their support (Award No.: 2017-0525). E.S.B. was supported by Lisa Yang, John Doerr, the Open Philanthropy Project, Cancer Research UK Grand Challenge grant C31893/A25050, U. S. Army Research Laboratory and the U. S. Army Research Office under contract/grant number W911NF1510548, NSF Grant 1734870, NIH UF1NS107697, NIH IDIQ17X149, NIH 1RM1HG008525, NIH 1R01MH103910, NIH 1R01MH114031, NIH 1R01MH110932, NIH 1R01EB024261, NIH 1R01DA045549, IARPA D16PC00008, NIH 1U19MH114821, NIH 1R01NS102727, the Chan Zuckerberg Initiative Human Cell Atlas pilot program, the Ludwig Foundation, and the HHMI-Simons Faculty Scholars Program. This project has been funded in part with Federal funds from the National Cancer Institute, National Institutes of Health, Task Order No. HHSN261100039 under Contract No. HHSN261201500003I.

## Author contributions

S.A., D.G., A.S., A.T.W., F.C., E.R.D., G.M.C., A.H.M. and E.S.B. contributed key ideas and designed experiments; S.A., A.T.W., A.S., and F.C. performed experiments; S.A., D.G., A.T.W., A.S., F.C., A.H.M. and E.S.B. analyzed data and wrote the paper; D.G., F.C., Y.B., A.K., A.G.X. conceived and implemented image analysis pipeline with assistance from S.A. and initial discussions with E.R.D; D.G., Y.B., A.K., A.G.X., K.M. and R.P. analyzed the imaging data; A.T.W., A.S., F.C., A.H.M., conceived and implemented targeted ExSeq; D.G. designed and implemented 3D visualization tool; D.G. analyzed cDNA movement experiments; S.A. and D.G. performed 3D Tracing; Y.C., A.C.P., C.C.Yao., P.T., P.R., and R.E.K. contributed to protocol optimization; S.A., F.C., H.-J.S., R.W. automated the *in situ* sequencing; F.C., E.R.D., A.H.M. conceived passivation; S.A., F.C. implemented passivation; S.A., F.C. conceived and implemented *ex situ* sequencing with initial discussions with E.R.D and A.C.P.; S.A., F.C. implemented *ex situ* and *in situ* sequencing matching with assistance from D.G.; S.A., A.T.W., A.S., F.C. conceived and implemented antibody staining; E.R.D. conceived and implemented cDNA fragmentation protocol and performed experiments on earlier preliminary protocols (data not shown); S.A., F.C. and E.R.D. optimized FISSEQ enzymatics for the expanded gel; S.A. and F.C. conceived and implemented cDNA anchoring with initial discussions with E.R.D.; C.-C.Yu, performed C. elegans fixation, cuticle reduction, and permeabilization; N.P. carried out mouse surgeries; J.A.M. and E.S.L. designed the gene list for the visual cortex experiment; A.L., N.C., S.R., K.H., D.L.A., N.W., B.E.J., J.K., M.S., J.W., J.J-V, O.R.R., A.R. designed the gene list and provided the human sample for the cancer experiment; S.L, S.P., E.I., contributed to targeted ExSeq; A.H.M., E.S.B., G.M.C. initiated the project; E.S.B. supervised the project.

## Data and materials availability

All the raw Illumina sequencing data will be deposited to NCBI Sequence Read Archive (SRA) upon publication. The entire MATLAB pipeline to process ExSeq datasets from the microscope to spatial analysis of gene expression is publicly accessible at GitHub http://www.github.com/dgoodwin208/ExSeqProcessing. The GitHub repository includes a tutorial wiki with step-by-step instructions on how to run the pipeline, and a tutorial set of targeted ExSeq data from the mouse visual cortex. The 3D visualization tool is hosted on GitHub at https://github.com/dgoodwin208/ExSeqOF3DVisualizer.

## Supplementary Materials

### Overview of fixation and use of mice

The first step of ExSeq is tissue fixation. For mouse tissues, we have demonstrated ExSeq using three different fixation conditions: hippocampal culture fixation (Fig. 2A), fixation of fresh frozen brain slices (Fig. 2C), and transcardially paraformaldehyde (PFA) perfused mouse brain (Fig. 2E). All methods for animal care and use were approved by the Massachusetts Institute of Technology Committee on Animal Care and were in accordance with the National Institutes of Health Guide for the Care and Use of Laboratory Animals. All solutions below were prepared from nuclease-free reagents. The mice used for Fig. 2 and Fig. 3 (excluding Fig. 2A**-B)** were Thy1–YFP (Tg(Thy1–YFP)16Jrs) male mice in the age range 6–8 weeks. Hippocampal neurons (Fig. 2A**-B**) were prepared from postnatal day 0 or day 1 Swiss Webster (Taconic) mice. No sample-size estimate was performed, since the goal was to demonstrate a technology. As noted in ref. Dell, R.B., Holleran, S. & Ramakrishnan, R., Sample size determination, ILAR J.43, 207– 213 (2002), “in experiments based on the success or failure of a desired goal, the number of animals required is difficult to estimate.” As was also noted in this paper, “the number of animals required is usually estimated by experience instead of by any formal statistical calculation, although the procedures will be terminated [when the goal is achieved].” No exclusion, randomization, or blinding of samples was performed.

### Fixation of hippocampal culture (Fig. 2A)

Hippocampal neurons were prepared as previously described (*91, 92*). ∼1000 hippocampal neurons were cultured per coverslip. Two weeks after the culture was prepared, the neurons were fixed using 10% formalin in 1X PBS for 15 min at 25°C, then washed with 1X PBS three times and finally stored in 70% ethanol at 4°C until use.

### Fixation of fresh frozen brain slice (Fig. 2C)

Mice were terminally anesthetized with isoflurane, then decapitated, and the brain dissected out into a cryomold with OCT. The cryomold was then placed in a dry ice/isopentane bath.

Overall, freezing of the brain was completed within 5 min after euthanasia. 15 µm slices were sliced on a Cryotome (Leica) and then immediately fixed with ice cold 10% formalin in 1X PBS for 12 min. Slices were washed 3 times for 5 minutes each with ice cold 1X PBS, treated with ice cold 4% SDS in 1X PBS for 2 minutes, washed again with ice cold 1X PBS three times, and finally stored at 4°C in 70% ethanol until use.

### Fixation via transcardial perfusion (Fig. 2E)

Mice were terminally anesthetized with isoflurane and perfused transcardially with ice-cold 4% paraformaldehyde, and the brain dissected out, and left to post-fix in 4% paraformaldehyde at 4 °C for one day, before moving it into 1X PBS containing 100 mM glycine. 50 µm slices were sliced on a vibratome (Leica VT1000S) and stored at 4°C in 70% ethanol until use.

### RNA anchoring

RNAs were anchored to the hydrogel network using the chemical LabelX as previously described (*24*). Briefly, LabelX is the commercial chemical reagent Label IT® (Mirus Bio LLC) modified, by simple mixing, with the hydrogel-anchorable group acryloyl-X. In more detail, specimens were washed with 1x PBS three times to remove residual ethanol, then once with 1X MOPS buffer (20 mM MOPS pH 7.7) for 45 min (15 min for neuronal culture samples), and finally reacted overnight at 37°C with 0.018 g/L LabelX in 1X MOPS buffer.

### Gelling, digestion and expansion

To allow expansion, the specimens were gelled as previously described (*93*). Briefly, tissue specimens were incubated in gelling solution (1X PBS, 2 M NaCl, 8.625% (w/v) sodium acrylate, 2.5% (w/v) acrylamide, 0.15% (w/v) N,N′-methylenebisacrylamide, 0.01% 4-hydroxy-TEMPO, 0.2% (w/v) ammonium persulfate (APS) and 0.2% (w/v) tetramethylethylenediamine (TEMED)) for 10 min at 4°C (for neuronal culture samples this incubation step is not needed).

The specimens were then placed in a gel chamber, constructed by sandwiching the sample between a slide and a coverglass, with #0 coverglass spacers on either side of the sample to prevent contact between the biological specimen and the coverglass. Specimens were incubated with gelling solution at 37°C for 1.5-2 hours until gelling was completed. Next, the specimens were incubated overnight at 37°C in digestion buffer (8 units/mL Proteinase K (New England Biolabs, cat. no. P8107S), 50 mM Tris pH 8.0, 1 mM EDTA, 0.5% Triton X-100, and 5 mM 2-Amino-5-methoxybenzoic acid (for formaldehyde adduct removal (*94*)). Specimens were then expanded by incubation in excess volumes of ddH2O 3 times, for 45 mins each time.

### Re-embedding

To prevent gel conformational changes during sequencing, expanded gels were re-embedded into non-expanding polyacrylamide gels, as previously described (*24*), resulting in a final expansion factor of ∼3.3x. Briefly, the specimens were incubated while rocking for 30 min at 25°C with re-embedding monomer solution (acrylamide and N,N-Methylenebisacrylamide (3% and 0.15% (w/v), respectively), 5 mM Tris base, 0.075% (w/v) TEMED, 0.075% (w/v) APS). Specimens were then placed in gel chambers, constructed by sandwiching the sample between slides and coverglasses, with two #1.5 coverglass spacers (placed one on top of the other) on either side of the sample to prevent contact. Gel chambers were placed inside a closed Tupperware-like container which was filled with nitrogen for about 5 min. Gel chambers were then incubated for 1.5 hr at 37°C.

### Passivation

We discovered that the carboxylic acid groups of sodium acrylate, used in the polymer of the expanding gel, could inhibit downstream enzymatic reactions important for *in situ* sequencing (**Fig. S1**). This might have been due either to a chelation effect or to interaction between the negative charges of the carboxylic group and the enzymes tested. A passivation procedure was designed to block the negative charges of carboxylic groups after expansion. The samples were treated with 1-Ethyl-3-(3-dimethylaminopropyl)carbodiimide (EDC), N-Hydroxysuccinimide (NHS) to covalently react ethanolamine to the carboxylic groups, converting them to amides with no charge. This reaction was performed in two steps (*95*): the first was incubation at 25°C for 2 hours with 2 M ethanolamine hydrochloride, 150 mM EDC, and 150 mM NHS in 100 mM 2-(N-morpholino)ethanesulfonic acid (MES) buffer at pH 6.5; the second step was incubation at 25°C for 40 min with 2 M ethanolamine hydrochloride and 60 mM sodium borate (SB) buffer at pH 8.5. The samples were then washed three times with 1X PBS for at least an hour overall. We note that as EDC can react with guanine in RNA (*96*), therefore higher concentrations of EDC, compared to the concentration used here, might have undesired side effects.

### Library preparation for *in situ* sequencing

Following expansion, re-embedding and passivation, several enzymatic steps were carried out to prepare the samples for *in situ* sequencing. Some of these steps were adapted from the FISSEQ protocol (*23*), but modified to better work with expanded gels. Briefly, anchored RNA underwent reverse transcription, cDNA circularization, and rolling circle amplification with phi29 polymerase. The details of the reactions are below.

First, endogenous DNA was removed to allow for later *ex situ* sequencing of cDNA without DNA contamination after library preparation was complete. The specimens were incubated with DNase I (Roche; MilliporeSigma cat. no. 4716728001) at 0.5 U/µL and RNase inhibitor (Enzymatics, cat. no. Y9240L) at 0.4 U/µL in 1X DNase I buffer for 2 hours at 25°C. The reaction underwent heat inactivation at 75°C for 5 min, and finally washed for 1 hour with 1X PBS.

Next, anchored RNAs were further biochemically modified into *in situ* sequence-able form via reverse transcription. Specimens were incubated with 10 U/µl SuperScript IV (SSIV) reverse transcriptase (ThermoFisher, cat. no. 18090050), 0.4 U/µL RNase Inhibitor (Enzymatics), 5 mM DTT, 250 µM dNTP, 10 µM inosine, 1X SSIV RT Buffer and 2.5 µM random octamer reverse transcription primer. The primer sequence was/5Phos/ACTTCAGCTGCCCCGGGTGAAGANNNNNNNN for the 15 micron thick hippocampal slice (Fig. 2C), and /5Phos/TCTCGGGAACGCTGAAGANNNNNNNN for the 50 micron thick hippocampal slice and the hippocampal culture (Fig. 2E and 2A, respectively). (The latter primer is shorter, so less self-circularization is anticipated.) The inclusion of inosine allowed generation of ∼100 base long cDNA fragments in a subsequent step using Endonuclease V cutting. We performed this step because CircLigase has lower efficiency for longer cDNA strands, and therefore the overall yield was improved by including inosine. The reverse transcription reaction was done overnight at 37°C after 15 min of 4°C incubation. For the 15 micron thick hippocampal slice, to save time, the reverse transcription reaction was done with thermo-cycling using the following program: (a) 8°C for 12 min, (b) 8°C for 1 min, (c) 37°C for 4 min, (d) back to (b) 70 times.

For the hippocampal culture and the 50 micron thick hippocampal slice (Fig. 2A and 2E), 40 µM aminoallyl-dUTP was also included in the reverse transcription mixture. These samples were formalin fixed after the reverse transcription to anchor aminoallyl-dUTP-modified cDNAs to the acrylamide moieties in the gel, to ensure that cDNAs would not move during subsequent steps (**Fig. S4**). The samples were washed with 1X PBS twice for 15 min, then incubated with 4% formaldehyde in 1X PBS for 1 hour at 25°C, and finally washed three times with 1X PBS for 15 min. For the 15 micron slice, Fig. 2C, no cDNA anchoring was performed; some small amount of motion of resultant cDNAs may have occurred (**Fig. S4**). (We note that since cDNA is not generated in targeted ExSeq, and the padlock probes are expected to remain stationary around the anchored RNA transcript (*97*), no post-fixation was performed in these samples; see below).

As single-stranded cDNA is required for the circularization step, the samples were treated with RNase. The specimens were incubated for 2 hours at 37°C with 0.01 U/µL Riboshredder RNase blend (Epicentre, cat. no. 12500), 0.25 U/µL RNase H (Enzymatics, cat. no. Y9220L), 0.05 U/µL Endonuclease V (New England Biolabs, cat. no. M0305S) and 1X NEB 4 buffer. For the 15 micron thick hippocampal slice (Fig. 2C), 1 mg/mL RNase DNase-free (Roche; MilliporeSigma cat. no. 11579681001) was used instead of Riboshredder RNase blend, as the latter was temporarily out of stock.

Next, the specimens were rinsed with nuclease-free water twice to remove traces of phosphate and the cDNA underwent circularization. The circularization reaction ran for 3 hours at 60°C with 3 U/µl CircLigase II (Epicentre, cat. no. CL9025K), 1 M betaine, 2.5 mM MnCl_2_ and 1X CircLigase buffer.

The specimens were then hybridized with the rolling circle amplification primer, which is reverse complement to the reverse transcription primers mentioned above: TCTTCAGCGTTCCCGA*G*A (* is phosphorothioate to block the exonuclease activity of phi29) for the hippocampal culture and the 50 micron thick hippocampal slice (Fig. 2A and 2E), and TCTTCACCCGGGGCAGCTGAA*G*T (* is phosphorothioate) for the 15 micron thick hippocampal slice (Fig. 2C). The hybridization was done for 2 hours at 37°C with 0.5 µM of the rolling circle amplification primer, 30% formamide and 2X SSC buffer. The specimens were then washed for 30 min at 37°C with 30% formamide and 2X SSC buffer, and then with 2X SSC, 1X SSC and 1X PBS, each for 5 min at 25°C.

Next, rolling circle amplification was performed overnight at 30°C with 1 U/µl phi29 DNA polymerase (Enzymatics, cat. no. P7020-HC-L), 250 µM dNTP, 40 µM aminoallyl dUTP and 1X Phi29 buffer.

To cross-link the amplified cDNA molecules containing aminoallyl dUTP, the specimens were washed with 1X PBS, and then incubated for 2 hours at 25°C with 5 mM BS(PEG)9 in 1X PBS. Then the samples were washed with 1X PBS and the reaction was quenched with 1 M Tris pH 8.0 for 45 min at 25°C.

Finally, to generate Fig. 2A, **C** and **E**, the specimens were incubated with the ATTO 565-labeled hybridization probe, matching the reverse transcription primers mentioned above: /5ATTO565N/TCTCGGGAACGCTGAAGA for the hippocampal culture and the 50 micron thick hippocampal slice (Fig. 2A and 2E), and /5ATTO565N/ACTTCAGCTGCCCCGGGTGAAGA for the 15 micron thick hippocampal slice (Fig. 2C). The hybridization was done for 45 min at 25°C with 0.1 µM hybridization probe, 10% formamide, 4X SSC and then the samples were washed with 2X SSC, 1X SSC and 1X PBS at 25°C for 5 min each.

### Automated *in situ* sequencing

SOLiD chemistry was utilized to sequence the cDNA amplicons (*23*). We automated the sequencing procedure by combining a dedicated spinning disk confocal microscope with computer-controlled fluidics. The spinning disk used was a Yokogawa CSU-W1 coupled with a Nikon Ti-E inverted microscope with Borealis modification. The fluidics system was composed of a FCS2 flow cell (Bioptechs), modular valve positioner with HVXM 8-5 valve (Hamilton) and PTFE laboratory tubing (S1810-12, Finemech). The flow was controlled with a syringe pump for the 15 micron thick hippocampal slice (Fig. 2C), and peristaltic pump for the hippocampal culture and the 50 micron thick hippocampal slice (Fig. 2A and 2E). (The peristaltic pump is accurate enough for high flow rates, and less prone to stop pumping in the middle of an experiment, than the syringe pump.) In both cases, a National Instruments Data Acquisition (NI-DAQ) card was used to connect the pump and the valve positioner to the scope computer. To mitigate movement during sequencing, the specimens were re-embedded again onto a Bind-silane (GE17-1330-01, GE Healthcare) treated coverslip, and the coverslip with the specimen then placed inside the flowcell. The treatment of coverslips with Bind-silane is described in (*24*), and the re-embedding was performed using the same protocol as in the ‘Re-embedding’ section above with the addition of fluorescent beads (Tetraspeck 0.2 µm, Life Technologies (T7280), diluted 1:100 into the re-embedding solution), to allow color correction as described below. The hippocampal culture and the 50 micron thick hippocampal slice (Fig. 2A and 2E) were re-embedded on the same coverslip and sequenced together, whereas the 15 micron thick hippocampal slice (Fig. 2C) was re-embedded on a different coverslip and sequenced separately. The lines of the valve positioner were populated with the chemicals required to perform the SOLiD sequencing by ligation *in situ*. The chemicals that required temperature other than 25°C were temperature controlled with Mini Dry Baths (Fisher Scientific).

For SOLiD sequencing by ligation chemistry, the fluidics setup was utilized to perform the following sequence of reactions; for each one of the 5 sequencing primers (**Table S1**), the specimen was first stripped for 30 min to remove the hybridization probe or the previous sequencing primer with strip solution (80% formamide and 0.01% Triton-X in water). Next, the specimen was washed with 1X instrument buffer (SOLiD Buffer F, 1:10 diluted) for 10 min, and incubated for 20 min with 2.5 µM of sequencing primer in 5X SASC (0.75 M sodium acetate, 75 mM tri-sodium citrate, pH 7.5). After a 5 min wash with 1X instrument buffer, the specimen was reacted for 1 hour with T4 DNA ligation mixture (6 U/µl T4 DNA ligase (Enzymatics, cat. no. L6030-HC-L) and 1:40 diluted SOLiD sequencing oligos in 1X T4 DNA ligase buffer). The specimen was then washed for 1 hour with 1X instrument buffer and imaged with SOLiD imaging buffer. To acquire the next base a two-step cleave reaction was performed, first with SOLiD buffer C for 30 min (part #4458932) and then SOLiD buffer B for 15 min (part #4463021). The cleave reaction was followed by a 10 min wash with 1X instrument buffer, 1 hour with T4 DNA ligation mixture, 1 hour wash with 1X instrument buffer and finally imaging with SOLiD imaging buffer (SOLiD buffer A, part #4463024). The cleave-ligation-wash-imaging cycle was repeated 3 times for each one of the 5 sequencing primers. For the hippocampal culture and the 50 micron thick hippocampal slice, a dephosphorylation reaction was performed before each cleave reaction to reduce phasing. (The 15 micron slice experiment was done earlier, before we realized that phasing could be an issue; we recommend the phosphatase for routine work.) The reaction was for 30 min with 1:20 dilution of Quick CIP (NEB, cat. no. M0508L) in 1X CutSmart buffer. All reactions were done at room temperature; the strip solution and the sequencing primers were kept at 80°C; the ligation mixture, the imaging buffer, and SOLiD buffer B were kept at 4°C.

The following optical configuration was used for *in situ* sequencing: (a) laser lines - 150 mW solid state OPSL 488 laser, 100 mW solid state OPSL 560 laser, 100 mW solid state DPSS 594 laser, 110 mW solid state OPSL 642 laser; (b) emission filters - 525/50, 582/15, 624/40, 685/40; (c) camera: Zyla sCMOS plus 4.2 megapixel with 100 msec exposure time; (d) objective: Nikon 40X CFI Apo, water immersion with long working distance, NA 1.15; (e) laser power and resolution: for the 15 micron thick hippocampal slice 50% laser power was used for all laser lines and the resolution was set to 0.5 microns in the Z axis and 0.17 microns in X & Y axis. For the hippocampal culture and the 50 micron thick hippocampal slice 100% laser power was used for all laser lines and the resolution was set to 0.4 microns in the Z axis and 0.17 microns in X & Y axis. The imaging time for one 40X field of view with the above configurations and 150 z-sections was ∼7 min using a piezo stage. As SOLiD chemistry takes ∼3.5 hours per sequenced base (see above), the chemistry+imaging time is ∼5hours for 10 fields of view per base and ∼100hours for a 20 base long sequencing experiment. We note that total experiment time scales with the number of bases sequenced, and therefore sequencing of short barcodes is faster, see **Methods** section ‘Targeted ExSeq of Visual Cortex and Hippocampus’ below.

### Morphology (Fig. 2F and Fig. 3)

Following *in situ* sequencing, the 50 micron thick hippocampal slice from the Thy1-YFP mouse was first stripped for 30 min to remove the sequencing primer with strip solution. Next, the specimen was washed with 1X instrument buffer for 10 min, and incubated for 50 min with 2.5 µM of hybridization probe in 4X SASC and 10% formamide. After a 10 min wash with 1X instrument buffer, the specimen was treated with 10 µg/mL primary antibody against GFP (Rabbit Anti-GFP, Invitrogen) in 5X SASC and 0.1% Triton-X for 24 hours. After a 3 hour wash with 1X instrument buffer, the specimen was reacted with secondary antibody, 4 µg/mL donkey anti-rabbit IgG CF®633 dye (Biotium) with 5X SASC and 0.1% Triton-X, for 24 hours. Finally, after an additional 3 hour wash with 1X instrument buffer, the sample was imaged in an imaging buffer. All reactions were done at room temperature, the strip solution was kept at 80°C, and the imaging buffer was kept at 4°C.

### Software overview and open source accessibility

The entire MATLAB library to process ExSeq datasets from the microscope to spatial analysis of gene expression is publicly accessible at http://www.github.com/dgoodwin208/ExSeqProcessing. Specifically, there is a tutorial wiki at http://www.github.com/dgoodwin208/ExSeqProcessing which includes instructions how to run the pipeline and a tutorial set of targeted ExSeq data from the visual cortex. **Fig. S2** is an illustrated overview of the pipeline.

### Image processing - Deconvolution

Deconvolution was used in all imaging channels for the two untargeted ExSeq experiments of the hippocampus (Fig. 2D**,F**) because the morphology signal was low, but was not used for other experiments. Deconvolution was done with Huygens software from Scientific Volume Imaging, using the CMLE mode with 10 iterations and a signal to noise ratio of 10.

### Image processing - Color Correction

For color correction, each 3D image volume from a round of SOLiD sequencing was acquired in four fluorescence channels. In the case of rigid offsets for the images between fluorescence channels, which can occur depending on the acquisition order on the microscope, color correction was done using the fluorescent beads as a reference (see **Methods** section ‘Automated *in situ* sequencing’). This function is included in the software pipeline repository published with this paper (see **Fig. S2**).

### Image processing - Feature-based 3D Registration

Feature-based registration pipelines have three fundamental parts: keypoint detection, feature construction at a keypoint, and feature matching. Matched features then create corresponding points between different image volumes, which can then be used to calculate a warp for one image into the coordinate space of another image. We implemented this pipeline, as described below. We selected the round with the best signal to noise ratio (typically one of the early SOLiD sequencing rounds) as the reference round for registration, and warped the other rounds of imaging to match the reference. The image data was downsampled for the first four steps, providing matched features that were re-mapped into the original space for the warp calculation. All code is included in the open source repository given with this paper.

a. Normalization: The four fluorescence channels from the SOLiD chemistry were combined into one 3D image before registration. Because each channel has a different fluorescence distribution, quantile normalization was used to normalize each fluorescence image and the adjusted color channels are then summed into one grayscale image.
b. Calculation of 3D keypoints: The Harris Corner Detector (*98*), adapted to 3D as previously applied to 3D hydrogel registration (*99*), identified candidate points in the image that would be identifiable across all sequencing rounds.
c. Calculation of 3D SIFT features: a feature vector was constructed at each keypoint to describe the point and its local spatial context (*100*). For a complete description, please refer to (*101*); briefly, the image gradients were calculated in the immediate neighborhood of each keypoint, at multiple scales and in multiple directions, and a vector was created from the polar histograms of the various gradients.
d. Matching of 3D keypoints: (*100*) presented a matching algorithm for SIFT features in 2D which also is effective in the 3D SIFT features used here. To additionally filter the matches, we applied a RANSAC operation to discard correspondences that are outliers to the resulting affine warp estimated by the majority of keypoint correspondences.
e. Warping to the reference round: once the quality correspondences were identified, a warp was calculated and applied to all SOLiD sequencing channels of the original data. We implemented both an affine transform (a global, linear operation) and a Thin Plate Spline (*102*) (which can produce much finer, localized adjustments at the cost of computational complexity), and ultimately an affine transform was sufficient when used in combination with the second re-embedding of the sample to the coverslip (see **Methods** section ‘Automated *in situ* sequencing’ above). For less than 1% of the fields of view, the affine transform was insufficient as determined by visual inspection and the thin plate spline was then used to improve the registration.

### Image processing - Background Subtraction

When manual inspection using FIJI (*103*) determined the presence of a non-uniform background signal, a background subtraction was performed by applying a morphological opening operation (of size 5 pixels, roughly the radius of an amplicon), then subtracting the opened image from the original image and clamping any negative values at zero.

### Image processing - Puncta segmentation

To identify and extract the pixels associated with each amplified cDNA, we utilized the long-established methodology of a watershed transform. To increase the signal, we first summed the registered image volumes (the grayscale 3D images from the combined normalized channel volumes then applied a background subtraction method) of the *in situ* sequencing rounds into a single volume. This composite image was then interpolated via a shape-preserving piecewise cubic interpolation in Z to achieve isotropic voxel size and the punctate signal was amplified using a Difference of Gaussians filter (similar to the commonly used Laplacian of Gaussians, used in (*10, 104*)). The filtered image was thresholded using the Otsu method, and the binarized image was segmented using the watershed transform. After segmentation, the data was un-interpolated in Z back into the original image size. We note that by design, this segmentation strategy was less likely to generate false negatives (i.e. puncta without segmentation) and more likely to generate false positives (i.e. not existing puncta); the latter are then removed by the following quality control steps. The resulting sections were identified using connected components, and candidate puncta that were too small or too large were removed, based on a manually set threshold (less than 30 pixels, greater than 2000 pixels). Additionally, we enforced the constraint that a puncta was present in all sequencing rounds; because the background subtraction clamps pixel values at 0, the presence of fluorescent signal in any channel for a particular *in situ* sequencing round was readily detected. If a puncta was missing a fluorescent signal in any channel for more than one sequencing round, it was discarded from consideration.

### Data Analysis - Basecalling

After cDNA amplicons were segmented using a 3D watershed approach, the pixels comprising the amplicons were processed for base calling. For each sequencing round, the pixels for the four color channels were quantile normalized to account for differences in fluorophore characteristics. For each channel in each amplicon, the normalized intensities of the channels were sorted and the average of the top 30 pixels were compared between channels, creating a sorted vector V. The highest channel was selected, and the confidence calculation was calculated as follows:

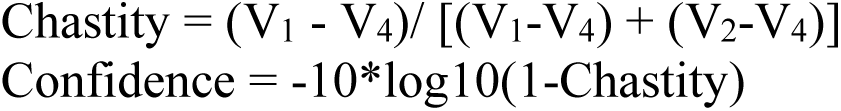

That is, the color intensities for each puncta were shifted to zero according to the dimmest color channel, V_4_, then the chastity was calculated as the ratio of the brightest color to the sum of the top two brightest colors. This is similar to Illumina’s chastity calculation, and the confidence, in rough analogy to a Phred score, is scored logarithmically. We then applied a filter to the candidate *in situ* reads, requiring that no more than 6 bases be called with a confidence less than 5.

### *Ex situ* sequencing

The specimens were first stripped for 30 min to remove the sequencing primer with strip solution, and washed with 1X instrument buffer for 10 min. The hydrogel was then digested with 20 mM sodium meta-periodate in 1X PBS pH 6 at 37°C for 12 hours, followed by 2 min of vortexing. The DNA was extracted and purified from the digested hydrogel with Genomic DNA Clean & Concentrator Kit (Zymo Research), and then the single stranded DNA was transformed into dsDNA with NEBNext (New England Biolabs). After additional purification with the Genomic DNA Clean & Concentrator Kit, the Nextera XT DNA Library Preparation Kit (Illumina) was used to simultaneously fragment and tag the dsDNA with the adapter sequences required for the Illumina MiSeq/NextSeq *in vitro* sequencing. The Nextera XT Index Kit (Illumina) was used to add Illumina sequencing barcodes during the PCR amplification of the extracted DNA. For the 15 micron thick hippocampal slice a MiSeq instrument was used to generate paired-end sequencing reads from the extracted DNA with MiSeq® Reagent Kit v3 (2X300 cycles; Illumina). For the hippocampal culture and the 50 micron thick hippocampal slice a NextSeq instrument was used to generate paired-end sequencing reads from the extracted DNA with NSQ® 500 Mid Output Kit v2 (2X150 cycles; Illumina).

### Data Analysis - Annotation of the *ex situ* sequencing data

The paired-end reads from the *ex situ* sequencing were aligned against both the mouse mRNA database (RefSeq genes, downloaded from the UCSC genome browser on 2018/08/14) and the mouse genome (version GRCm38/mm10). In both cases local alignment was used with Bowtie2 software (*105*) using default settings, and the output format was converted into browser extensible data (bed) format using the samtools package (*106*). For alignment against the genome, to avoid alignments in repetitive regions reads that were given a Bowtie mapping quality score of 10 and below were filtered out. Minimum mapping quality was not set for the alignment against mRNA to allow alignments against genes with several alternative splicing isoforms. To unify the alignments against the mRNA and the genome, the alignment against the mRNA was converted into genomic coordinates. This was done with a custom Perl script, using the alignment of the RefSeq genes against the mouse genome in bed format (downloaded from the UCSC genome browser on 2018/08/14). After converting all alignments to genomic coordinates, reads aligned against genomic regions with low complexity, detected with the WindowMasker software with SDust module (*107*) (coordinates downloaded from the UCSC genome browser on 2018/08/15), were filtered out using the intersect function of BEDTools (*108*); default settings were used with ‘-f 0.2’, i.e. no more than 20% of the read was allowed to overlap with the low complexity regions. In addition, reads aligned against mRNA regions that couldn’t be mapped in the genome were discarded. The aligned reads that passed the above mentioned filters were annotated using UCSC gene information to rRNA (28S and 18S), tRNA, introns, exons of non-coding genes, and exons of coding genes. In addition, we detected all reads that mapped to known alternative splicing events (UCSC alternative splicing events dataset). All the gene information and the known alternative splicing events were downloaded in bed format from the UCSC table browser (on 2018/08/22), and the intersection was determined with BEDTools as described above. RefSeq annotations, i.e. transcripts names (NM and NR accessions) and associated gene symbols, were downloaded from the NCBI website (on 2018/08/20). Cases in which the two mates of a paired-end read were aligned against two different transcript names were allowed if they represented the same gene symbol (as in the case of two splice variants of the same gene); however, paired-end reads were filtered if the two mates were aligned against two different gene symbols.

### Data Analysis - *Ex situ* and *in situ* sequence matching

The *in situ*-generated 20 base long sequences (see Data Analysis - Basecalling), in SOLiD color space (henceforth referred to as *in situ* color vectors), and the annotated *ex situ* sequences were matched as follows; first, as the SOLiD sequencing started from the 5’ of the sequencing primer, this primer sequence was identified in the paired-end *ex situ* reads. The software BLASTn using default settings and e-value cutoff of 0.05 was utilized for primer identification. For each paired-end read, we required that at least one of the paired-end mates matched the sequencing primer and that the match included the primer 5’ end. To generate 20 base long fragments equivalent to the *in situ* sequences, the 17 bases at the 5’ end of the sequencing primer were then collected from the *ex situ* read, and combined with the three 5’ bases of the sequencing primer, which are the first bases read in the SOLiD sequencing chemistry. To minimize sequencing errors, these 17 bases were collected only if each base had a Phred score of 30 or above. Next, the resulting 20 base long sequences from the annotated *ex situ* reads were converted into SOLiD color space (henceforth referred to as the *ex situ* library). We note that converting sequences in base space to SOLiD color space is straightforward and direct, especially as only bases with low probability of sequencing errors were used, and therefore the ambiguity of converting SOLiD color space to base space was avoided (*20, 109*); for example, in converting from SOLiD color space to base space, all bases after a sequencing error would be converted incorrectly (*109*). To avoid non-informative *ex situ* and *in situ* matching, a low-complexity filter was applied both to the *in situ* color vectors and the *ex situ* library, requiring a minimum Shannon entropy of 1. Overall, the resulting annotated *ex situ* library had 710,659 entries.

To match the *in situ* color vectors to the annotated *ex situ* library, we employed a continuous distance function. Briefly, we reshaped each *in situ* color vector and *ex situ* library entry into 68-length vectors (17 *in situ* sequencing rounds x 4 colors) and normalized each sequencing round to 1 by dividing by the brightest channel’s value (setting the max value to one every four entries in the 68-length vector). The *ex situ* library was transformed into a matrix of size 710,659×68, termed G, and the *in situ* color vectors were transformed into a similar matrix of size 394,957×68, termed C. All alignments were done independently for each one of the fields of view. The alignment distance, A, is calculated by A = 17-CG^T^. Each row of A is the alignment distance to each entry of G, so the column index for the minimum distance in each row of A is the index for the matching *ex situ* library entry. This distance calculation requires a threshold under which a putative match could be considered an actual match, and that threshold was determined by setting a target false positive rate. To quantify the false positive rate of the *ex situ* and *in situ* sequence matching, we shuffled the order of all the *in situ* color vectors and re-aligned using the same method as described above. Because there is a monotonic relationship between threshold distance and the false positive rate, we can set the threshold distance based on a desired false positive rate, which was set at 15%. We demanded that the *in situ* color vectors will uniquely match the *ex situ* library. This was achieved by removing all the entries in G (the *ex situ* library) that were aligned to C (the *in situ* color vectors), producing G’. We then realigned the *in situ* color vectors against G’, and counted any entries of C that align to G’ as a non-unique alignment that will be removed. We determined that 92% of all alignments (including rRNA) were unique, and importantly, 97% of the non-rRNA alignments were unique. As expected, this uniqueness is dependent on the number of bases sequenced *in situ* (**Fig. S5C**). All non-unique matches were discarded from further analysis.

Overall, we matched 115,075, 15,709, and 35,835 *in situ* color vectors with a false positive rate of 15%, 5%, and 15%, for the 50 micron thick hippocampal slice (Fig. 2E), 15 micron thick hippocampal slice (Fig. 2C), and the hippocampal culture (Fig. 2A), respectively. We note that reducing the false positive rate (FDR) reduces the number of aligned reads; for example, reducing the false positive rate to 10% and 5% for the 50 micron thick hippocampal slice results in 96,857 and 63,306 aligned reads, respectively. However, reducing the FDR to 10% or even 5% doesn’t change the results presented in this paper. Qualitatively, expected synaptic and cytoskeletal RNAs, such as Kcnq2 and Map1b, are still obtained, as well as almost all the genes discussed in the main text: Map2, Sptbn1, Mga, Ptma, BC1, Malat1, Grik2, Gabrg2, Nob1, Map1b, Gria1, Eef1a2, Calm3, Nob1, Rbfox1, Shtn1, Ddx5, Celf2, Syt1, Syp, Josd2, Rps24 (with FDR of 10%), excluding only Cux1 and Foxg1. Quantitatively, the functional enrichment groups in the lists of genes obtained with the different FDR are very similar to one another, and to the functional enrichment groups obtained with the RNAseq dataset (**Table S6**). Moreover, the correlation between RNAseq and untargeted ExSeq with 10 volumes (Fig. 2Gv) still holds with the different FDR: Pearson’s r of 0.41 (p-value 6.2 x 10^-93^) and 0.39 (p-value 5 x 10^-50^) with 10% and 5% FDR, respectively, compared to Pearson’s r of 0.47 (p-value 9 x 10^-164^) with FDR of 15%. Finally, the agreement between untargeted ExSeq and targeted ExSeq (**Fig. S22D**) still holds with the different FDR: Pearson’s r of 0.57 (p-value 4.1 x 10^-4^) and 0.53 (p-value 1.4 x 10^-3^) with 10% and 5% FDR, respectively, compared to Pearson’s r of 0.68 (p-value = 9.75 × 10^-6^) with FDR of 15%.

The expression levels of all RefSeq genes detected using ExSeq in all these conditions are given in **Table S2-S4**. For the 50 micron thick hippocampal slice, the RNAseq is also included in **Table S2**. All the raw Illumina sequencing data will be deposited to NCBI Sequence Read Archive (SRA) upon publication.

### Data Analysis - *Ex situ* sequencing data sampling

Sampling from the full *ex situ* sequencing data, for Fig. 2G, was performed as follows:

a. Paired-end reads were randomly selected from the full *ex situ* sequencing fastq file. The number of lines selected was such that the total number of counts for all RefSeq genes in the random set was 10 times that of the number in 10 FoVs, to simulate the datasets for 100 FoVs.
b. The random set of paired-end reads was then analyzed similarly to the procedure described in the **Methods** section ‘Annotation of the *ex situ* sequencing data’; briefly, the reads were aligned against the mouse RefSeq genes using local alignment with Bowtie2 software (*105*). The alignments against the mRNA were converted into genomic coordinates, and reads aligned against genomic regions with low complexity were filtered out. Cases in which the two mates of a paired-end read were aligned against two different transcript names were allowed if they represented the same gene symbol (as in the case of two splice variants of the same gene); however, paired-end reads were filtered if the two mates were aligned against two different gene symbols.
c. For each RefSeq gene, the number of aligned paired-end reads was counted, resulting in an expression vector that was compared to the expression vector generated using the RNAseq data; the Pearson’s correlation between the log-transformed expression vectors was calculated. In addition, the fraction of RefSeq genes detected using the random set of paired-end reads, compared to RNAseq, was calculated. (d) Steps (a-c) were repeated 10 times. Box plots are presented in Fig. 2Gvi**-vii**.

### Comparison of untargeted ExSeq to bulk RNAseq (Fig. 2G)

To perform a comparison of untargeted ExSeq to bulk RNAseq of a matching tissue sample, a coronal hippocampal slice was selected that was within 100 µm of the original 50 micron thick hippocampal slice (Fig. 2**-3**) along the antero-posterior axis. The coronal hippocampal slice was fixed with 4% PFA in exactly the same way as the original slice, washed with 1X PBS and was kept in 70% ethanol at 4℃ until RNA extraction; throughout the experimental procedure of sequencing the RNA from a matching sample, we followed the steps in the **Methods** section ‘Library preparation for *in situ* sequencing’ when possible. Accordingly, the DNA was removed using DNase I (Roche). RNA was extracted from these slices using an RNeasy FFPE Kit (Qiagen) following the “Deparaffinization using Melting” protocol indicated in the kit. The extracted RNA was then reverse transcribed using SSIV reverse transcriptase with random primers, following the manufacturer’s protocol. The RNA was removed using RNase DNase-free (Roche) and RNase H. The single stranded cDNA was transformed into ds-cDNA with NEBNext second strand synthesis (New England Biolabs). After purification with the Genomic DNA Clean & Concentrator Kit (Zymo), the Nextera XT DNA Library Preparation Kit (Illumina) was used to simultaneously fragment and tag the dsDNA with the adapter sequences required for the Illumina *in vitro* sequencing. The Nextera XT Index Kit (Illumina) was used to add Illumina sequencing barcodes during the PCR amplification of the extracted DNA. The library was sequenced using a MiSeq instrument to generate paired-end sequencing reads with the MiSeq® Reagent Kit v3. The resulting sequencing reads were annotated as described in the **Methods** section ‘Data Analysis - Annotation of the *ex situ* sequencing data’ above.

To perform the comparison of untargeted ExSeq to bulk RNAseq of a matching hippocampal culture (**Fig. S8**), we followed the same steps as for the hippocampal slice above, with the following differences: (1) whereas one coverslip with hippocampal neurons was used for ExSeq, five such coverslips (cultured and fixed at the same time as the one used for ExSeq, see **Methods** section ‘Fixation of hippocampal culture’) were used for the matching RNAseq experiment, to compensate for the fact that the RNA is not RCA amplified in the RNAseq experiment; (2) the neurons in the five matching coverslips were first treated with ProK to allow detachment from the coverslips and collection in an eppendorf tube.

In FISSEQ, highly abundant genes, such as genes involved in translation and splicing, were underrepresented (*20*). We examined if such a detection bias exists in the untargeted ExSeq dataset by comparing it with the matching RNAseq dataset (**Table S2**). First, we detected genes that were expressed in the RNAseq dataset but were not expressed in the untargeted ExSeq with full *ex situ* sequencing data in a 50 micron thick hippocampus slice; three expression cutoffs were used: 10, 20, or 50 counts, and for each cutoff genes with counts higher or equal to the cutoff in the RNAseq, but not in the untargeted ExSeq dataset (normalized to the RNAseq dataset by the total number of reads), were detected. Gene ontology analysis using the software package DAVID (*110*) was then performed on the detected genes with each cutoff, revealing only 2, 0, and 0 enriched ontologies (Benjamini p-value<0.01) with the expression cutoffs of 10, 20, or 50 counts, respectively. Therefore the underrepresentation of highly expressed genes from specific functional groups, reported in FISSEQ, is not observed with untargeted ExSeq. The two functional enrichment groups (Benjamini p-value<0.01) as obtained by DAVID for the expression cutoff of 10 counts are given below:

### Image Processing - 3D Tracing (Fig. 3)

To be considered a read inside a cell, the pixels of an amplicon must overlap with the annotated neuron morphology. The annotations were created by manually tracing Thy1-YFP antibody signals using Vast Lite (*111*). A cell was annotated as three parts: the soma, the dendrites and the nucleus (visually determined by the high perinuclear density of amplicons, and the reduction of puncta density inside the nucleus).

### 3D Viewer (Fig. 3)

In order to visualize the richness of the ExSeq information, we developed a 3D visualization tool using OpenFrameworks (an open-source platform for graphics and interactivity, v.0.9.8, (*112*)). Code is hosted on GitHub at https://github.com/dgoodwin208/ExSeqOF3DVisualizer.

We used the cytoplasmic YFP antibody data to produce a 3D mesh of the exterior of the neuron. We then loaded each *in situ* read in space, along with its readtype (exon, intron, etc.) and gene symbol (e.g, ‘*Atp2c1*’,’*Camk2a*’). Using the exportTo3DVisualizer.m function included in the GitHub repository, the *in situ* reads were converted from MATLAB objects into a format readable by ExSeq viewer for exploration.

### Targeted ExSeq of Visual Cortex and Hippocampus (Fig. 4-5)

#### Gene selection

For the mouse primary visual cortex, gene panels were selected using a combination of manual and algorithm-based strategies, as described previously (*113*) and below, and required a reference single cell and single nucleus RNAseq data set from the same kind of tissue (in this case, ∼12,000 single cells in mouse primary visual cortex) (*114*). First, cells were re-assigned to a more refined set of 192 types, using the published types as a starting point and a consensus of several computational methods to “over-split” the data. Second, an initial set of high-confidence marker genes were selected through a combination of literature search and analysis of the reference data. These genes were used as input for a greedy algorithm (detailed below). Third, the reference RNAseq data set was filtered to only include genes compatible with smFISH. Retained genes had to be: 1) long enough to allow probe design (> 960 base pairs); 2) expressed highly enough to be detected (FPKM >= 10), but not so high as to overcrowd the signal of other genes in a cell (FPKM < 500); 3) expressed with low expression in off-target cells (FPKM < 50 in non-neuronal cells); and 4) differentially expressed between cell types (top 1000 remaining genes by marker score). To more evenly sample each cell type, the reference data set was also filtered to include a maximum of 50 cells per cluster. We note that these genes were selected to be compatible with both smFISH (*113*) and ExSeq and therefore are more stringent than is required for ExSeq alone.

The main step of gene selection used a greedy algorithm to iteratively add genes to the initial set. To do this, each cell in the filtered reference data set was mapped to a cell type by taking the Pearson’s correlation of its expression levels with each cluster median using the initial gene set of size n, and the cluster corresponding to the maximum value was defined as the “mapped cluster”. The “mapping distance” was then defined as the average cluster distance between the mapped cluster and the originally assigned cluster for each cell. In this case a weighted cluster distance, defined as one minus the Pearson’s correlation between cluster medians calculated across all filtered genes, was used to penalize cases where cells are mapped to very different types, but an unweighted distance, defined as the fraction of cells that are not mapped to their assigned cluster, could also be used. This mapping step was repeated for every possible n+1 gene set in the filtered reference data set, and the set with minimum cluster distance was retained as the new gene set. These steps were repeated using the new gene set (of size n+1) until a gene panel of 42 genes was attained. Code for reproducing this gene selection strategy is available as part of the mfishtools R library (https://github.com/AllenInstitute/mfishtools).

For experiments on the hippocampus, 35 genes of interest that were highly expressed in the synaptic neuropil in rats were selected from a prior study (*64*), and converted to their mouse homologs.

#### Probe Design - Overview

For each gene selected to be interrogated *in situ*, a set of DNA oligonucleotide padlock probes directly targeting the RNA transcript was designed. Each probe had four key parts: (1) a 32 nucleotide (nt) homology region (split into two 16 nt portions spanning the ligation junction at the 5’ and 3’ ends of the padlock probe); (2) a constant backbone region for RCA initiation and *in situ* sequencing primer binding; (3) a barcode 5’ to the constant region for SOLiD sequencing readout; and (4) a barcode 3’ to the constant region for Illumina sequencing readout. The homology regions were linked to the SOLiD and Illumina barcodes with a short linker sequence ‘AAA’. Transcripts were assigned barcodes in a logical barcode space (described below), which was represented in sequence space by two nucleotide sequences, one for readout using the SOLiD chemistry and the other for readout using the Illumina chemistry. This provided flexibility, enabling either chemistry to be used to read out the sequences. A schematic of the probe and amplicon product is shown in **Fig. S12.**

#### Probe Design - Homology and constant region

Sequences of interest were downloaded from RefSeq in GenBank format. A sliding-window sequence walk was performed down the length of a transcript, in which 32-mer regions were serially selected, and tested to identify regions passing our QC criterion. If the region passed, the window was advanced to start 5 nt past the end of the current window. If the region did not pass, the window was advanced 1 nt. As the 32-mer sequence was split between two 16-mer regions on the 5’ and 3’ ends of the probe, part of the QC screened each half individually. Our QC criterion for homology regions excluded any homology region with: (1) repetitive sequences (>4 consecutive identical bases); (2) GC content for either 16-mer half < 40% or > 65%; (3) melting temperature for both 16-mer halves > threshold (for visual cortex probes, the threshold was originally 55°C, then subsequently lowered (to a minimum Tm of 52°C) if fewer than six probes per gene were generated; for hippocampus probes, the threshold was fixed at 50°C); (4) significant hairpin or dimer secondary structure; (5) a BLAST hit of the homology region against the mouse transcriptome (excluding the gene of interest) of >12 nt in length, that spanned the ligation junction by at least three nt on either side of the ligation junction. 32-mers passing the QC criterion were saved.

The constant region on the probe backbone was TCT CGG GAA CGC TGA AGA CGG C, a modified version of the universal primer sequence from FISSEQ (*20*), that was extended to increase its melting temperature for compatibility with Illumina sequencing chemistry.

#### Probe Design - Barcode design

Transcript barcodes uniquely identifying transcripts were designed in a logical space, consisting of four rounds of imaging, in which a logical 0, 1, 2, or 3 was decoded in each round of imaging, i.e. 3021 corresponded to reading out a 3 on the first round of imaging, a 0 in the second round of imaging, and so on. Each logical color corresponded to a physical color when read out using SOLiD or Illumina chemistries. Barcodes for transcripts were designed with the first three rounds R1, R2, R3 being independent, and with the last round being a check-sum round equal to R4 = (R1 + R2 + R3) (mod 4), i.e. 3021 is a valid barcode since 3 + 0 + 2 (mod 4) = 1, while 3022 is not a valid transcript barcode. For the hippocampus probeset, bulk expression data (from the original study (*64*)) was used to assign barcodes such that the barcodes were more evenly distributed in colorspace across all rounds of imaging. For the visual cortex dataset, the barcode sequences were evenly distributed in colorspace without weighting by gene expression.

#### Probe Design - Barcode implementation in SOLiD and Illumina chemistry

SOLiD barcodes were designed to be seven nt long, with two ligations on the first primer (SeqN), and two ligations on the second primer (SeqN-1). Dibase encodings correspond to logical bases such that longer wavelengths correspond to higher logical bases, i.e. dibase encodings for the lowest wavelength fluorophore corresponds to logical 0, while dibase encodings for the longest wavelength fluorophore corresponds to logical 3, except in the first ligation on the SeqN-1 primer, in which case the color-mapping is reversed (due to a technical error in one base of the barcode design; we note that since the Illumina chemistry was used with these probes, this error has no effect on the result). All possible 7 nt barcode sequences were interpreted, and formed a library of 64 sequences per 4-round barcode.

Illumina barcodes were designed to be seven nt long, with the first four bases synthesized directly corresponding to the first four readout rounds, with the mapping between logical bases and sequence bases being ({0, 1, 2, 3} -> {A, C, G, T}) (and with the shortest to longest wavelength order being G, T, A, C). All possible 7 nt barcode sequences were interpreted, and formed a library of 64 sequences per barcode.

#### Probe Design - Probe assembly, negative control probes, ordering and pooling

For the visual cortex probeset, a maximum of 8 probes per transcript were selected, and for the hippocampus probeset, a maximum of 16 probes per transcript were selected (we expect that using 16 probes per gene will result in higher yield and is recommended to the user). To assemble probe sequences, the correct number of homology regions were randomly selected from the set of acceptable homology regions for a particular transcript. Each homology sequence was associated with a SOLiD and an Illumina barcode sequence that corresponds to the logical barcode associated with the transcript. The construction of the probe is described above.

In parallel, three negative control probe sets targeting the mCherry-expressing plasmid pMExt589 (*115*), the *D. melanogaster* gene Vg, and random barcodes from a transcriptome-orthogonal barcode set (*116*) were also designed as described above, with BLAST screening against the both the human and mouse transcriptome. These probes were given distinct barcodes from the transcript barcodes, enabling detection of these negative control probes.

Probes were ordered with 5’ phosphate modifications in 96-well plate format from either Integrated DNA Technologies (IDT) or Eurofins Genomics. Probes for each gene were pooled together to form 100 µM or 200 µM subpool solutions. Subpools were pooled together to form stock solutions containing all probes of a particular probeset. Visual cortex and hippocampus barcodes are in **Table S9**, with probe sequences in **Table S10**. For the hippocampus probeset, one gene, Rgs5, was assigned to a highly repetitive barcode (2222), and therefore was excluded *in silico* (as it was difficult to distinguish this simple repeat from imaging artifacts) after the sequencing data was collected by removing the entry from the barcode library used to match *in situ* reads.

#### Tissue preparation

For experiments involving study of the visual cortex (Fig. 4), one seven week old C57BL/6 Thy1-YFP female mouse was terminally anesthetized with isoflurane and perfused transcardially with ice-cold 4% paraformaldehyde. The brain was dissected out and left in 4% paraformaldehyde at 4°C for 12-16 hrs. After briefly washing the brain with 1X PBS, 50 µm slices were then prepared on a vibratome (Leica VT1000s) and stored in 70% ethanol at 4°C until use. For experiments involving study of the hippocampus (Fig. 5), one 14 week old C57BL/6 Thy1-YFP male mouse was similarly utilized.

#### Targeted ExSeq library preparation

Brain sections stored in 70% ethanol were rehydrated with two 15 minute washes with PBST (1X PBS, 0.1% Triton-X) at room temperature. A coronal slice was selected that contained the primary visual cortex, and another coronal slice from a different mouse (see above) was selected that contained the hippocampus. These were labeled with a primary antibody against GFP (which labels the YFP protein) at a concentration of 10 µg/mL Rabbit Anti-GFP (Thermo Fisher, A-11122), in PBST overnight at 4°C followed by staining with a biotinylated secondary antibody at 10 µg/mL (Thermofisher, B-2770), in PBST overnight at 4°C. We note that antibody staining which is performed pre-expansion (as described in this section) results in a better staining compared to antibody staining which is performed post-sequencing (as described in the **Methods** section ‘Morphology’ above); however, in our hands, *in situ* sequencing with SOLiD chemistry was not successful for slices that were antibody stained pre-expansion.

Therefore, for the samples described in this section, the Illumina chemistry was utilized for *in situ* sequencing instead of the SOLiD chemistry (see **Methods** section ‘Targeted ExSeq *in situ* sequencing by synthesis’ below).

To allow the retention of RNA, the sections were treated with 0.1 mg/mL LabelX overnight in 1X MOPS buffer (20 mM MOPS pH 7.7), as previously described. Gelation and digestion were performed as described above for untargeted ExSeq. Re-embedding was also performed as described above except with the following re-embedding solution composition: Acrylamide and N,N-Methylenebisacrylamide, (4% and 0.2% (w/v), respectively), 5 mM Tris base, 0.05% (w/v) TEMED, 0.05% (w/v) APS. Passivation was carried out as described above.

Probes were pooled together to form 200 µM stock libraries for the visual cortex probeset and 81 µM for the hippocampus probeset (these two probeset were purchased from different vendors with different probe stock concentrations). The gelled and passivated samples were then pre-incubated for 30 minutes at room temperature with wash buffer (20% formamide, 2X SSC buffer), then incubated overnight at 37°C with the respective hybridization mix. For the visual cortex experiments, the hybridization mix consisted of 54 µM pooled visual cortex library probeset (total of 334 probes; ∼162 nM each probe) in 20% formamide, 2X SSC buffer. For the hippocampus experiments, the hybridization mix consisted of 54 µM pooled hippocampus library probeset (total of 540 probes; 100 nM each probe), 4.4 µM negative control probes (total of 44 probes; 100 nM each probe) in 20% formamide, 2X SSC buffer.

The samples were then washed twice with wash buffer at 37°C for 30 minutes each, followed by a wash with 1X PBS at 37°C for 30 minutes, and a pre-incubation with 1X SplintR Ligase Buffer for 30 minutes at room temperature. The samples were then incubated with 1250 U/mL SplintR ligase (NEB, cat. no. M0375L) in 1X SplintR ligase buffer at 4°C for 6 hours. The samples were then incubated with a freshly prepared solution of 1250 U/mL SplintR ligase in 1X SplintR ligase buffer overnight at 37°C. Following ligation, the samples were washed with 2X SSC for 30 minutes at room temperature followed by a pre-hybridization with wash buffer for 15 minutes at room temperature.

The samples were then incubated with 500 nM rolling circle amplification primer (TCT TCA GCG TTC CCG A*G*A, where * denotes phosphorothioate backbone modification) in wash buffer for 2 hours at 37°C, followed by a 30 minute wash with wash buffer at 37°C, and another wash with 1X PBS for 15 minutes at 37°C.

After a pre-incubation with 1X Phi29 buffer for 15 minutes at room temperature, the samples were incubated with 1000 U/mL Phi29 polymerase (Enzymatics) in 1X Phi29 buffer for 6 hours at 4°C. Following this step, the samples were incubated with 1000 U/mL Phi29 polymerase, 250 µM dNTP, 40 µM aminoallyl dUTP in 1X Phi29 buffer at 30°C overnight. The next day, the samples were washed once with 1X PBS at room temperature for 30 minutes, and then treated with 5 mM BS(PEG)9 (ThermoFisher, cat. no. 21582) in 1X PBS for 2 hours at room temperature. The samples were then again washed with 1X PBS for 15 minutes at room temperature followed by another wash with 1 M Tris, pH 8 for 15 minutes at room temperature. After this step, the samples were then washed once with 1X PBS.

#### Targeted ExSeq *in situ* sequencing by synthesis

For the stable, multi-round imaging required for *in situ* sequencing, samples were immobilized to the bottom of a 24-well glass-bottom plate. To prepare the plate, wells were treated with Bind-silane (GE17-1330-01, GE Healthcare) as described previously (*24*).

Subsequently, the gelled samples were re-embedded in individual wells within a re-embedding gel (Acrylamide and N,N-Methylenebisacrylamide (4% and 0.2% (w/v), respectively), 5 mM Tris base, 0.05% (w/v) TEMED, 0.05% (w/v) APS) along with 0.2 µm TetraSpeck beads (Life Technologies, cat. no. T7280), which were diluted 1:100 in the re-embedding solution. A droplet of the re-embedding solution was transferred to a well of a bind-silane treated plate. The gel containing the sample was placed on top of the drop, and oriented so that the tissue was on the top of the sample (farthest from the glass). Another droplet of the re-embedding solution was added on top of the sample, and a 10 mm circular coverglass was placed on top. The remaining volume underneath the coverglass and around the gel was back-filled. The plate was placed into a Tupperware container which was purged with nitrogen gas for 5 min, then gelled at 37°C for 1.5 hours. Following gelation, the sample was washed with 1X PBS for 1 hour at room temperature.

To prevent background from base addition to exposed 3’ DNA ends of cellular DNA during sequencing, the surface-attached samples were blocked with dideoxynucleotides using Terminal deoxynucleotidyl transferase (TdT, New England Biolabs, cat. no. M0315L). First the samples were pre-incubated with 50 µM ddNTP, 250 µM CoCl_2_, in 1X TdT buffer for 20 minute at room temperature. Then, the samples were incubated with 400 U/mL, 50 µM ddNTP, 250 µM CoCl_2_ in 1X TdT buffer for 90 minutes at 37°C. After TdT treatment, the samples were washed with 1X PBS for 30 minutes at room temperature, followed by a wash with 4X SSC for 20 minutes at room temperature. The Illumina sequencing primer (TCT CGG GAA CGC TGA AGA CGG C) was then hybridized to the amplicons at 2.5 µM sequencing primer in 4X SSC at 37°C for one hour. The samples were then washed four times with 4X SSC for 10 minutes each wash at 37°C.

At this step, the samples were ready for *in situ* sequencing via Illumina Sequencing-by-Synthesis. For this purpose, we collected the incorporation mix buffer (IMT), imaging buffer (SRE), and cleavage buffer (EMS) solutions from a MiSeq V2 kit, and aliquoted and stored them at −20°C. Each round of sequencing involves a step of base addition, imaging, and cleavage. For adding a base, samples were first washed three times for 10 minutes at room temperature with PR2 buffer (part of the MiSeq V2 kit), and then treated with MiSeq V2 IMT buffer for 20 minutes at room temperature, followed by another wash with IMT for 10 minutes at room temperature. Then, the plates with the re-embedded samples incubating in IMT were transferred to an incubator heated to 60°C for 10 minutes. Following this step, the samples were washed three times, 15 minutes per wash, with PR2 buffer at 60°C.

After base addition, samples were ready for imaging. The samples were stained with DAPI (1 mg/L) in PR2 buffer, and washed twice for 15 minutes at room temperature with the MiSeq V2 SRE imaging buffer. The samples were washed again with SRE imaging buffer, and were then imaged on an Andor Dragonfly Spinning Disk confocal with laser lines 100 mW solid state 405 laser, 150 mW solid state 488 laser 150 mW solid state 561 laser, 160 mW solid state 633 laser; emission filters, 450/50, 540/30, 631/36, 676/37, 775/140; Andor Zyla sCMOS 4.2 plus with 200 msec exposure time; and a Nikon 40X CFI Apo, water immersion with long working distance, NA 1.15 objective. Tiled images of the entire primary visual cortex and the hippocampus were acquired using the Andor Fusion software. For each tile, 151 Z-sections were imaged at a spacing of 0.4 µm.

After imaging, samples were then incubated with the Illumina cleavage buffer, first with two washes each for 10 minutes at room temperature, and then finally with an incubation with the cleavage buffer at 60°C for 20 minutes. Following this step, the samples were washed three times for 10 minutes with PR2 buffer. At this point, the cleavage step is complete and the process was repeated starting from the base addition step. A total of four rounds of base addition, imaging, and cleavage were performed for the visual cortex and hippocampus samples, though cleavage was not performed after the final round of sequencing in order to retain the signal from the final base addition. After the final round of sequencing was imaged, the samples were washed with PR2 buffer for 30 minutes at room temperature, followed by an overnight incubation at 4°C with 10 µg/mL Alexa 488-labeled streptavidin (ThermoFisher, cat. no. S11223) in PR2 to visualize YFP immunostaining. The samples were washed three times for 30 minutes with PR2 at room temperature, and stained with DAPI and imaging buffer (as described above for sequencing). Afterwards, all the fields of view in the samples were imaged including the streptavidin staining marking YFP as well as the signal from the final round of sequencing. By imaging the YFP morphology along with the final round of sequencing, the YFP protein signal was co-registered to the sequencing data.

#### Image Processing - Color correction, registration, segmentation

Color correction, registration and segmentation were all done as described in the **Methods** sections ‘Image processing’ above, with the sole exception that background subtraction was not used because it was not deemed necessary by manual inspection.

#### Image Processing - Basecalling and alignment to barcodes

Basecalling was done as described for the untargeted ExSeq (see **Methods** section “Data Analysis - Basecalling”) with one exception that candidate puncta were not filtered by confidence. Instead of filtering on confidence as was done in longer reads of untargeted ExSeq, puncta for targeted ExSeq were required to have at least one color channel “present” per sequencing round, where presence for a channel is defined as a signal above a channel’s median value across all puncta in a sequencing round. Puncta that did not have a present signal for each round were discarded. Alignment was done by determining the base for each amplicon per round, then calculating the Hamming distance of that sequence to the list of barcodes in the probeset. Only perfect matches (agreements across all four bases) were kept.

#### Image Processing - Visual cortex cell segmentation

Using the DAPI nuclear stain, 3D centroids of each neuron were determined via a thresholding of the nuclear stain. Briefly, each volumetric image had outlier pixel intensities removed by setting a maximum value at the 99% percentile of nuclear stain values. The image was then blurred to account for nonuniformity of nuclear structure, and then a threshold value was determined using Otsu’s method of all non-zero voxels in the image. This threshold value was then used to binarize the image, and connected components was used to filter out putative nuclear images of insufficient size (set as 50000 voxels, corresponding to a volume of 15.5 µm^3^). As some nuclei were erroneously split, nuclei with centroids within a distance of 5 microns of each other were merged into a single object (shown in **Fig. S14A**). The final set of nuclei then had their centroids calculated, which then were used to estimate each cell as a sphere of radius 12.5 µm. ExSeq reads within a cell’s volume were assigned to that cell. If a read was within the volume of multiple cells, it was assigned to the cell with the closest nuclear centroid (shown in **Fig. S14B**). This segmentation was repeated with a radius of 7.5 µm for the robustness analysis (**Fig. S18**).

#### Image Processing - Localization of reads within spines, dendrites, and cell bodies of the hippocampus (Fig. 5)

The locations of spine heads were identified using the commercially available software, MBF Neurolucida 360. Fields of view of interest were first loaded into the software one at a time. Dendrites of YFP expressing neurons were traced using the software’s semi-automated tracing modality. Following this step, spines on traced dendrites were automatically identified. The output of the software consists of the coordinate of spine heads as well as other physical characteristics which were used for subsequent analysis. Axons were traced using the software’s semi-automated tracing modality.

To localize reads within spine heads, a threshold was applied to the deconvolved images of YFP expressing cells to generate a binary mask. Reads were attributed to spine heads if the centroid of the transcript resided within a positive region of the mask and within 250 nm from the centroid of a spinehead.

Similarly, to localize reads within dendrites, a threshold was applied to the deconvolved images to generate a binary mask to be applied to the sequencing data. To identify dendrites in different regions of the hippocampus, the software VAST was used to annotate regions of the hippocampus on Z-projected stitched images of the entire dataset. Reads were then assigned to dendrites within annotated regions of the hippocampus depending on whether their centroid resided within dendrites. To identify reads within YFP cell bodies, VAST was used to manually create masks for YFP neuronal cell bodies within CA1 and the dentate gyrus. VAST was also used to segment the neuropil region of CA1 into 50 micron segments (Fig. 5E).

#### Data Analysis - ExFISH and ExSeq comparison in cultured cells (Fig. 4B)

HeLa cells were cultured on CultureWell Chambered Coverglasses (Thermo Scientific, C37000) in D10 medium (Cellgro) supplemented with 10% fetal bovine serum (FBS) (Invitrogen), 1% penicillin–streptomycin (Cellgro), and 1% sodium pyruvate (BioWhittaker). Cultured cells were washed once with DPBS (Cellgro), fixed with 10% formalin for 10 min, and washed twice with 1X PBS. Fixed cells were then stored in 70% ethanol at 4°C until use.

Cultured cells were treated with LabelX (0.01 mg/mL), and expanded, as described previously following the tissue gelation/expansion protocol (*24*). Samples were subsequently re-embedded and passivated as described above in the targeted ExSeq library preparation protocol.

A direct comparison of ExFISH to targeted ExSeq was performed by carrying out these processes sequentially within the same expanded cells. HCRv3.0-amplified ExFISH (HCRv3.0-ExFISH) was performed using a modified ExFISH protocol (as described below), with HCRv3.0 probes and reagents (Molecular Instruments) (*24, 55*). Modifications to the original protocol were designed to avoid using dextran sulfate in hybridization/amplification steps, as it was found to partly inhibit downstream enzymatic reactions. All steps were performed with gels in PCR strips, with 200 µL liquid volumes unless otherwise noted. To perform HCRv3.0 labeling, samples were pre-incubated with HCRv3.0 Wash Buffer for 30 minutes at room temperature.

Samples were then hybridized with HCRv3.0 probes targeting a single gene that were diluted in HCRv3.0 Wash Buffer (8 nM total final probe concentration; 1.6 pmol probes into 200 µL total volume) at 37°C overnight. Four samples were prepared, using HCRv3.0 probes against one of GAPDH, EEF2, TFRC, VIM (**Tables S9, S10**). Samples were washed with HCRv3.0 Wash Buffer four times for 30 minutes at 37°C, followed by four 15 minute washes with 5X SSCT (5X SSC buffer with 0.1% Tween-20) at RT. HCR amplification was carried out with the corresponding HCR amplifiers (hairpins) labeled with Alexa 546. Briefly, a pair of HCR amplifiers (i.e. BxH1 and BxH2, where x denotes the initiator type; stock solution 3 µM) were individually snap cooled by heating to 95°C for 90 seconds, then annealed at room temperature for 30 minutes in a dark drawer. The amplification buffer was prepared by mixing 2 µL of each snap cooled HCR amplifier with 96 µL of 5X SSCT, forming a total volume of 100 µL with final concentration 60 nM each hairpin. Samples were incubated in the amplification buffer for four hours at room temperature. After washing four times with 5X SSCT for 30 minutes each, gels were transferred to individual wells of a 24-well glass-bottomed plate. Gels were stained with DAPI (1 mg/L) in 5X SSCT for 10 minutes at room temperature. After staining, samples were imaged using a spinning-disc confocal microscope (see ‘Automated *in situ* sequencing’ above) above with a Z-step of 0.5 µm.

Following imaging, HCRv3.0 probes and amplifiers were stripped, and a targeted ExSeq library was prepared. To strip HCR reagents, samples were pre-incubated with 500 µL strip buffer (80% formamide, 20% water (v/v)) for 15 minutes at RT in the 24-well plate. Samples were then washed with 500 µL of strip buffer, and incubated at 37°C for 2 hours, followed by six 20 minute washes with 500 µL strip buffer at 37°C. After stripping, samples were transferred back to PCR tubes. Targeted ExSeq library preparation was then performed for the same gene that was interrogated with HCRv3.0-ExFISH. The library preparation was carried out as described above, using 8 padlock probes per transcript, at a concentration of 100 nM for each probe. After library preparation (ending with washing with PBS after BS-PEG9 cross-linking), amplicons were visualized by hybridizing an Alexa 546-labeled detection oligo (/5Alex546N/TCTCGGGAACGCTGAAGA, where /5Alex546N/ is the Alexa 546 modification) at 100 nM in 2X SSC with 10% formamide at 37°C for 1 hour. Samples were then washed twice with 2X SSC, 10% formamide for 15 minutes each at 37°C. Finally, the samples were washed with 1X PBS for 15 minutes at RT, transferred to individual wells of a glass-bottom 24-well plate, and stained with DAPI (1 mg/L). The same regions that were previously imaged were identified and imaged again on the same spinning disk confocal microscope, with a Z-step of 0.5 µm.

Analysis for each cell was spot-counting quantification for ExFISH amplicons and for targeted ExSeq amplicons for the same gene in the same cell. Using a custom spot-counting MATLAB code developed by the Raj lab (complete source code and instructions can be found at https://bitbucket.org/arjunrajlaboratory/rajlabimagetools/wiki/Home), spots were identified using manual thresholding, counted, and extracted from pairs of image stacks corresponding to ExFISH and targeted ExSeq for the same gene in the same cell. Pearson’s correlation was computed for the log10-transformed data.

### Data Analysis - Clustering and t-SNE embedding (Fig. 4D)

Analysis was performed using a custom MATLAB script. Segmented cells (as described previously) with greater than 50 ascribed reads were retained for subsequent analysis. The gene expression profiles of the cells were standardized using the Z-transform. Z-score values were used for subsequent analysis unless explicitly noted. *K*-means clustering was performed on the expression profiles using *K*=15 clusters. Briefly, *K*=15 was selected based on a prior study (*58*); the 49 clusters of the original paper were grouped into 15 groups: (1, 2) two groups of non-SST, VIP, PVALB GABAergic neurons; (3, 4) two groups of VIP neurons; (5, 6) two groups of SST neurons; (7, 8) two groups of PV neurons; (9) L2/3 neurons; (10) L4 neurons; (11, 12) two groups of L5 neurons; (13, 14) two groups of L6 neurons; (15) glia. The cells were embedded into a two-dimensional space using t-distributed stochastic neighbor embedding (t-SNE) directly on their gene expression profiles, with coloring of points in the t-SNE plot corresponding to their *K*-means cluster.

Clusters were annotated based on the expression of known and novel marker genes (using the dataset from a prior paper (*58*) or the Allen Mouse Brain ISH Atlas), and physical location of cells in the ExSeq dataset. Excitatory clusters (those with an annotation ending in “Ex”) were identified as those highly expressing at least one of the following marker genes (for corresponding cortical layer(s) indicated in parenthesis): Foxp2 (L6), Sez6 (L5, L6), Galnt14 (L5, L6), Fezf2 (L5), Kcnk2 (L5), Rorb (L4, L5), Lingo2 (L2/3), Cux2 (L2/3), or Pcdh8 (L2).

Excitatory clusters were ordered and named based on the physical ordering of the layers, and the known expression patterns of the marker genes. This resulted in the nine ExSeq clusters L6 Ex, L6a Ex, L5b Ex, L5/L5a Ex, L5 Ex, L4 Ex, L2/3 + L4 Ex, L2/3 Ex, and L2 Ex. Inhibitory clusters were identified by their low expression of the marker genes for excitatory neurons listed above. Of these six clusters, five had clear and unique marker genes. The cluster strongly expressing Pvalb was annotated as PV. The four clusters with marker genes Unc13c, Chodl, Nts, and Tnni3k were annotated as SST Unc13c, SST Chodl, SST Nts, and SST Tnni3k respectively based on the the specific expression of the marker genes in minimally-overlapping subsets of SST neurons in the prior study. The remaining cluster, ultimately annotated as GABAergic (-PV), indicating non-PV GABAergic neurons, expressed a number of marker genes including Gad2, Prox1, Npas1, and Nr2f2. These genes were studied in prior work (*58*). Gad2 was strongly expressed in nearly all interneuron types, Prox1 in VIP interneurons, Npas1 in a subset of VIP interneurons and a subset of non-PV/SST/VIP GABAergic neurons, and Nr2f2 in subsets of VIP interneurons and non-PV/SST/VIP GABAergic neurons (that did not overlap with those expressing Npas1). As Pvalb (and strongly correlated markers Ank1, Slc32a1, Lhx6, Kcnmb2) were not strongly expressed in this ExSeq cluster, the cluster was annotated as GABAergic (-PV) to denote the mixed GABAergic neuron composition, excluding Pvalb^+^ interneurons.

Annotation of the clusters for the robustness analysis was performed similarly. The same set of marker genes as above was used to identify excitatory clusters, which were named and ordered based on the physical ordering of the layers in space and the known expression patterns of the marker genes, resulting in the eight ExSeq clusters L6 Ex, L6a Ex, L5b Ex, L5/5a Ex, L4 Ex, L2/3 Ex, L2/3 + L4 Ex, L2/3 + L5b Ex. The latter two clusters included cells from multiple cortical layers. Annotation of the seven remaining clusters was performed similarly as above. Six of the seven clusters had strong marker genes. The cluster expressing Pvalb was annotated as PV, and clusters expressing Grin3a, Tnni3k, Unc13c, Chodl, and Nts were annotated as SST Grin3a, SST Tnni3k, SST Unc13c, SST Chodl, and SST Nts, respectively, based on the specific expression of the marker genes in minimally overlapping subsets of SST neurons in the prior study. The remaining cluster expressed a number of marker genes including Gad2, Prox1, and Nr2f2. As described above, these encompass multiple interneuron types. As markers of PV neurons (Pvalb, and strongly correlated markers Ank1, Slc32a1, Lhx6, Kcnmb2) and SST neurons (correlated markers Chodl, Cdh9, Thsd7a, Tnni3k, Nts) were not highly expressed in this cluster, the cluster was annotated as GABAergic (-PV, SST).

### Image Processing - Making multicolor images of reads in space (Fig. 4C**, 4E**)

Raw images were generated using a custom MATLAB script and merged using FIJI (*103*). For images showing reads, the relevant set of reads was loaded into a custom MATLAB script that Z-projected the reads into a 2D image, with centroids rounded to the nearest pixel (corresponding to 50 nm pre-expansion). For each read, a 2D circle with radius corresponding to 0.5 microns (pre-expansion) was drawn at the centroid location in the relevant color. The script produced three images representing RGB channels. Images were downsampled by a factor of 2 before saving. For Fig. 4C, all reads were used and colored identically; for **Figs. 4E, S14C-D, S18B**, reads were colored depending on the cluster assignment of the cell or randomly by cell as indicated. Images of the YFP morphology across the entire dataset were maximum intensity Z-projected and stitched together. The RGB images and YFP morphology images were merged in FIJI, with the YFP morphology shown in grey.

### Data Analysis - Correlation to previously known cortical cell types (Fig. 4F)

Data from a previous study was downloaded and RPKM data was imported into a custom MATLAB script (*58*). The 42 genes in the visual cortex dataset were identified and used for subsequent clustering analysis. Gene expression profiles for each cell were standardized using the Z-transform, and clustered using *K*-means with *K*=15 clusters. Clusters identified by performing *k*-means clustering on the previous dataset were annotated by examining the correlation of cells within the cluster to the 49 clusters previously identified. Clusters corresponding to excitatory neurons were annotated by their layer (L6 Ex; L6a Ex; L5b Ex; L5/L5a Ex; L4, L5 Ex; L2/3, L4 Ex; and L2/3 Ex). A glia cluster (annotated Glia) was also identified. The remaining clusters were identified as interneuron clusters. Four clusters had strong marker genes, and were clearly attributable to classical interneuron types, and were annotated by their classical type and marker gene: PV Ank1, PV Thsd7a, SST Unc13c, SST Chodl. Of the three remaining clusters, two corresponded to classical types: VIP (annotated without a marker gene as it was the only strongly VIP-correlated cluster), and PV (annotated without a marker gene as a strong subtype marker gene was not identified. The remaining cluster strongly correlated to non-PV/SST/VIP GABAergic neurons and was annotated as GABAergic (-PV/SST/VIP).

The mean expression profile for each cluster was computed. Mean expression profiles were also computed for clusters in the targeted ExSeq dataset, and the Pearson’s correlation matrix between mean cluster expression profiles was computed. This was also performed for the segmentation robustness analysis in **Fig. S18**.

The data from the prior study was also clustered using the variable genes within the 24,057 genes in the original dataset (*58*) (**Fig. S17**), instead of clustering using the 42 genes interrogated by ExSeq. First, genes that exceeded a minimal level of expression were selected by summing the genes’ RPKM values across all cells, and selecting genes with a sum greater than 1000, resulting in a subset of 13,392 genes. Coefficients of variance (CV) were computed for each one of these genes, and genes with a CV greater than 0.75 were retained, resulting in a final subset of 12,604 genes. Downstream analysis was performed as described above, with a similar annotation approach. Six clusters corresponding to excitatory neurons were annotated by their layer(s) (L6a Ex, L5b Ex, L5a + L6b Ex, L4 Ex(1), L4 Ex(2), L2/3 Ex). The L5a + L6b Ex cluster spanned multiple cortical layers, and two similar L4 clusters were identified (L4 Ex(1) and L4 Ex(2)).

Seven interneuron clusters were identified and annotated by the expression of marker genes (VIP Nr2f2, VIP Dlx4, VIP Kcnip4, PV, SST Npas1, SST Thsd7a, SST Chodl). The remaining interneuron cluster corresponded to a mix of non-PV/VIP/SST interneuron types, and was annotated GABAergic (-PV/VIP/SST). Finally, a cluster corresponding to glial cells was annotated as Glia.

### Data Analysis - Distribution of cell types across cortical layers (Fig. 4G)

Images of the transcriptomically-defined clusters corresponding to excitatory neurons (from the clustering analysis) were exported into RGB images using custom MATLAB scripts and FIJI, as described above. The images were partitioned into layers with VAST (*111*), using the DAPI nuclear staining and coronal section images from the Allen Reference Atlas (https://mouse.brain-map.org/static/atlas)(60) to guide the segmentation boundaries. Seven compartments were identified: (1) external capsule (corresponding to white matter below the visual cortex); (2) L6/L6a; (3) L5b; (4) L5/L5a; (5) L4; (6); L2/3; and (7) L1. The layer segmentations were exported as images and loaded into a custom MATLAB script, which first computed the centroids of cells as the mean of the reads ascribed to a cell, and then assigned cells to layers according to the location of the centroid. Bar plots of the cluster distribution by physical layer and the physical layer distribution by cluster were subsequently generated.

### Data Analysis - Comparison of targeted ExSeq to *ex situ* sequencing and bulk RNAseq

To perform comparison of targeted ExSeq to *ex situ* sequencing of libraries, the expanded hippocampal slice processed with targeted ExSeq was processed for *ex situ* sequencing via digestion, DNA fragmentation, amplification, and MiSeq Illumina sequencing as described above in the Methods section, “*Ex situ* sequencing”. The resulting sequencing reads were aligned against the probe sequences using Bowtie2 (*105*) with local alignment and default settings. The output of Bowtie2 was parsed to calculate the number of sequencing reads aligned against each probe sequence, and this information was then merged to give the number of sequencing reads aligned against each targeted gene (as several probes were used for each targeted gene).

To perform a comparison of targeted ExSeq to bulk RNAseq of a matching tissue sample, two coronal hippocampal slices, from the same mouse, were selected that were within 100 µm of the hippocampal slice along the coronal axis. RNA was extracted from these slices using an RNeasy FFPE Kit (Qiagen) following the “Deparaffinization using Melting” protocol indicated in the kit. The extracted RNA was then reverse transcribed and PCR amplified using the NEBNext Ultra II RNA Library Prep Kit using the universal PCR primers of the kit. The amplified cDNA was then sequenced on a MiSeq system following the instructions for a “4 nM library”. The resulting sequencing reads were aligned against mouse mRNA (RefSeq genes, downloaded from the UCSC genome browser on 2018/08/14), using Bowtie2 (*105*) with local alignment and default settings. The output of Bowtie2 was parsed to calculate the number of sequencing reads aligned against each mouse mRNA.

### Targeted ExSeq of Metastatic Breast Tumor Sample (Fig. 6)

#### Gene selection and probe design

To select a set of 300 genes for spatial profiling in metastatic breast cancer (MBC) samples, a preliminary list of ∼ 600 potentially relevant genes was first assembled based on prior knowledge and literature, as well as MBC single cell and single nucleus datasets (in preparation). Genes were chosen to represent various aspects of breast cancer biology, metastasis, and the tumor-immune-microenvironment, as well as cell types and programs discovered from the single cell and single nucleus RNAseq data. The preliminary list was then filtered down to 300 genes based on expression statistics as measured in the MBC single cell RNAseq data set. During probe design, three of the selected 300 genes were excluded as they did not meet technical criteria (specifically all three transcripts were too short), reducing the final gene set to 297 genes. For each selected gene we designed up to 16 probes and no less than 9 probes. The design of the probes was as described in the **Methods** section ‘Targeted ExSeq of Visual Cortex and Hippocampus’ above, with the following two exceptions: (a) the barcodes were generated using the R package DNABarcodes (*117*); we used barcodes of length 7 that had a hamming distance of 3 between them (with the following command: create.dnabarcodes(7, dist=3, heuristic=“ashlock”,cores=12, filter.gc=FALSE,population=500, iterations=500), therefore allowing correction of one substitution error in the *in situ* sequencing step; (b) we followed the design in **Fig. S12**, with the SOLiD and Illumina barcodes having the same sequence; the SOLiD one was encoded in color space and the Illumina one is in base space. Accession numbers and probe sequences are in **Tables S9** and **S10**, respectively.

#### Tissue preparation

As part of an ongoing research study on metastatic breast cancer, biopsies are collected from patients at Dana Farber Cancer Institute. Prior to any study procedures, the patients provide written informed consent for a research biopsy and subsequent analysis of tumor and normal samples, as approved by the Dana-Farber/Harvard Cancer Center Institutional Review Board (DF/HCC Protocol 05-246). For this study, we used an 18-gauge core needle biopsy (∼6×0.8 mm) of a liver metastasis obtained from a 66-year-old woman with a known diagnosis of hormone receptor positive metastatic breast cancer. Surgically dissected metastatic tumor samples were quickly washed with 1X PBS and placed into a cryomold with OCT. The cryomold was then placed in a dry ice/isopentane bath. 8 µm slices were prepared on a Cryotome (Leica) and adhered to Superfrost Plus glass sides, which were then immediately fixed with ice cold 10% formalin in 1X PBS for 12 min. Slices were washed 3 times for 5 minutes each with ice cold 1X PBS and finally stored at 4°C in 70% ethanol until use.

#### Targeted ExSeq library preparation

To allow the retention of RNA, tumor sections were treated with 0.1 mg/mL LabelX overnight in 1X MOPS buffer (20 mM MOPS pH 7.7). Gelation was performed as described above for untargeted ExSeq of brain slices. A modified Proteinase-K digestion protocol was used, consisting of 1 hour of digestion at 60°C, followed by 24 hours of digestion at 37°C, until the gel came off the glass slide. The digestion buffer was modified to include guanidine hydrochloride; final composition 50 mM Tris pH 8.0, 1 mM EDTA, 0.5% Triton X-100, 0.8 M guanidine hydrochloride, 8 U/mL Proteinase-K (NEB). Re-embedding and passivation were performed as described above for targeted ExSeq of the mouse visual cortex and hippocampus.

Probes were pooled together to form 200 µM libraries for the MBC probeset. The gelled and passivated samples were pre-incubated for 30 minutes at room temperature with wash buffer (20% formamide, 2X SSC), then incubated overnight at 37°C with the hybridization mix, consisting of 140 µM pooled library probeset (total of 4353 probes; ∼32 nM each probe) in 20% formamide, 2X SSC buffer. The samples were then washed three times with wash buffer at 37°C for 30 minutes each wash, followed by one wash with 1X PBS for 30 minutes at 37°C. The samples were then pre-incubated with 1X SplintR ligase buffer for 30 minutes at room temperature. The samples were then incubated with 1250 U/mL SplintR ligase in 1X SplintR ligase buffer at 4°C for 3 hours. The samples were then incubated with a freshly prepared solution of 1250 U/mL SplintR ligase (NEB) in 1X SplintR ligase buffer overnight at 37°C. Following ligation, the samples were washed with 2X SSC for 30 minutes at room temperature followed by a pre-hybridization with wash buffer for 15 minutes at room temperature. The samples were then incubated with 500 nM rolling circle amplification primer (TCT TCA GCG TTC CCG A*G*A, where * denotes phosphorothioate backbone modification) in wash buffer for 2 hours at 37°C, followed by a 30 minute wash with wash buffer at 37°C, and another wash with 1X PBS for 15 minutes at 37°C. After a pre-incubation with 1X Phi29 buffer for 15 minutes at room temperature, the samples were incubated with 1000U/mL Phi29 polymerase (Enzymatics) in 1X Phi29 buffer for 3 hours at 4°C. Following this step, the samples were incubated with 1000 U/mL Phi29 polymerase, 250 µM dNTP, 40 µM aminoallyl dUTP in 1X Phi29 buffer at 30 °C overnight. The next day, the samples were washed once with 1X PBS at room temperature for 30 minutes, and then treated with 5 mM BS(PEG)9 in 1X PBS for 2 hours at room temperature. The samples were then again washed with 1X PBS for 15 minutes at room temperature followed by another wash with 1 M Tris, pH 8 for 15 minutes at room temperature. After this step, the samples were then washed once with 1X PBS.

#### Targeted ExSeq *in situ* SOLiD sequencing

To enable stable, multi-round imaging, gelled samples were re-embedded in individual wells of a 24-well glass-bottom plate as described above for targeted ExSeq of mouse visual cortex and hippocampus.

For SOLiD sequencing by ligation chemistry, for each one of the 5 sequencing primers, the specimen was first stripped for 30 min to remove the hybridization probe or the previous sequencing primer with strip solution (80% formamide and 0.01% Triton-X in water). Next, the specimen was washed with 1X instrument buffer (SOLiD Buffer F, 1:10 diluted) for 10 min, and incubated for 20 min with 2.5 µM of sequencing primer in 5X SASC (0.75 M sodium acetate, 75 mM tri-sodium citrate, pH 7.5). After two washes for 5 min total with 1X instrument buffer, the specimen was reacted for 1 hour with T4 DNA ligation mixture (6 U/µl T4 DNA ligase (Enzymatics) and 1:40 diluted SOLiD sequencing oligos in 1X T4 DNA ligase buffer). The specimen was then washed 4 times for 3 hours overall with 1X instrument buffer and imaged with SOLiD imaging buffer (the extended wash was needed since we didn’t use the flowcell but rather did manual sequencing).

After each ligation, samples were ready for imaging. The samples were stained with DAPI (1 mg/L) in 1X instrument buffer, and washed twice with an imaging buffer. The samples were then imaged on an Andor Dragonfly Spinning Disk confocal with laser lines 100 mW solid state 405 laser, 150 mW solid state 488 laser 150 mW solid state 561 laser, 160 mW solid state 633 laser; emission filters, 525/50, 582/15, 624/40, 685/40; Andor Zyla sCMOS 4.2 plus with 200 msec exposure time; and a Nikon 40X CFI Apo, water immersion with long working distance, NA 1.15 objective. Tiled images of the sample were acquired using the Andor Fusion software. For each tile, 151 Z-sections were imaged at a spacing of 0.4 µm.

Following each imaging round, a dephosphorylation reaction was performed before each cleave reaction to reduce phasing. The reaction was for 30 min with 1:20 dilution of Quick CIP (NEB, M0508L) in 1X CutSmart buffer. To acquire the next base, a two-step cleave reaction was performed, first with SOLiD buffer C twice for 30 min total (part #4458932) and then SOLiD buffer B twice for 15 min total (part #4463021). The cleave reaction was followed by three washes for 20 min total with 1X instrument buffer, 1 hour with T4 DNA ligation mixture, 4 washes for 3 hours total with 1X instrument buffer, and finally washing with SOLiD imaging buffer (SOLiD buffer A, part #4463024) before imaging. The cleave-ligation-wash-imaging cycle was done once for each one of the 5 sequencing primers. All reactions were done at room temperature. During the *in situ* sequencing reactions the strip solution and the sequencing primers were kept at 80°C while the ligation mixture, the imaging buffer, and SOLiD buffer B were kept at 4°C.

#### Image Processing - Color correction, registration, segmentation, basecalling and alignment

Color correction, registration, segmentation basecalling were done as described for the targeted hippocampus and visual cortex experiments with two modifications. Background subtraction, as described in the untargeted hippocampus **Methods** sections, was performed. The second modification is that alignment allowed up to one mismatch in the 7-base barcodes, due to the barcode design (described in the **Methods** section ‘Gene selection and probe design’ above) which allows correction of one substitution error.

#### Image Processing - Cell segmentation (Fig. 6)

Because of the high density of cells of the tumor biopsy, the automated cell segmentation was unreliable. We utilized VAST (*111*) to implement a 2D manual segmentation on a stitched image of maximum intensity projections of the DAPI stain for the 78 fields of view. A single person was able to annotate the cells in one day by utilizing the “conditional” feature of VAST, allowing only pixels of a trained intensity regime to be annotated, which accelerates the speed of annotation. Additionally, because MATLAB’s ‘bwlabel’ function can ascribe unique values to all non-touching binary objects, the speed of manual annotation was increased by only using a single annotation color in VAST, and therefore the manual task was mainly to remove touching annotations of neighboring cells. With this pipeline, we segmented 5,862 cells containing at least one ExSeq read, and retained 2,395 cells with >100 ExSeq reads, containing a total of 771,904 reads. For an ExSeq read to be counted as within a cell, given the sample is 8 microns thick (comparable to the cell size), we demanded that the XY portion of the 3D centroid of the ExSeq read would be within the segmented nucleus of a cell.

Nuclear structures (possibly nucleoplasmic bridges) (**Fig. S25**) were manually identified in 2D using VAST and then extended into 3D using the DAPI stain, creating a 3D mask of the nuclear structures. For an ExSeq read to be considered inside the nuclear structure, we enforced that at least 10% of the puncta voxels overlapped with the 3D mask. We note that the relatively small number of reads (516) obtained in these nanoscale regions limited our ability to systematically classify these reads into specific cell types.

#### Data Analysis - Expression clustering (Fig. 6B and 6C)

In order to identify and cluster the 2,395 cells according to their expression pattern, we utilized the R toolkit Seurat (*74, 118*). Following (*119*), we used a supervised approach of using selected genes for dimension reduction (instead of choosing the genes with the highest variability). These genes were: non-tumor marker genes (*119*)-CD3G, CD68, FOXP3, CD4, CD8A, CD3D, CD3E, HLA-DRA; tumor marker genes (*119*)-EGFR, GRB7, ERBB2, PGR, CD44, CD24, ALDH1A3, EPCAM, KRT19, KRT18, CDH1; B-cells-IGHG1, IGHG4, IGKC, IGHM; Fibroblast-HSPG2. The rest of the analysis was according to the Seurat analysis pipeline, i.e. creating a K-nearest neighbor graph based on the euclidean distance in PCA space, and then finding clusters in the PCA space using the Louvain algorithm. Genes that can ‘mark’ each cluster (i.e. with an expression level which is higher in a given cluster compared to the other clusters) were discovered using the Seurat ‘FindAllMarkers’ function. The ‘FindAllMarkers’ function reports a p-value for each putative gene marker in each cluster; for putative gene markers with p-values less than 1E-10, we assigned the cluster with the known annotation of the marker gene, otherwise we marked the cluster as “unknown”.

#### Data Analysis - Determining adjacent cell clusters (Fig. 6D)

For all 2,395 cells classified by Seurat, cell centroids were calculated as the average position of the reads inside that cell. An adjacency graph was then calculated by counting all instances of two cell centroids, from two different cell clusters, being within 20 microns of each other. Cluster labels were shuffled and the adjacency graph of cell clusters was recalculated 500,000 times. From this bootstrapped data, p-values were calculated for each pair of cell clusters as (1+x)/500,000 where x=number of bootstrapped iterations in which the randomized adjacency graph for a given pair exceeded that of the original adjacency graph. The distance of 20 microns was empirically chosen, and 10 microns and 40 microns (39 microns was used in (*119*)) were also computed at 10,000 iterations each, producing a similar adjacency graph structure (**Fig. S26**).

#### Data Analysis - Detecting upregulated genes (Fig. 6E)

Using the cell clusters and adjacency matrix as described above, for any pair of cell clusters A and B, cluster A was partitioned into two subsets: a subset of A cells that are adjacent to B cells, and a subset of A cells that are not adjacent to B cells. Gene expression change (fold change) per gene was then calculated as the ratio of the median expression in the subset of A near B and the median expression in the subset of A not near B, ignoring any genes with median value of zero. To calculate statistical significance, the A cells were randomly partitioned into two subsets, of the same size as the two original subsets, and the fold changes of all genes between the two subsets were recalculated 100,000 times as described above. From this bootstrapped data, p-values were calculated for each gene as (1+x)/100,000 where x=number of bootstrapped iterations in which the fold-change for any gene exceeded that of the original detected gene.

We note that a discontinuity is clear in the right image in panel Fig. 6Ei -- an horizontal line is evident in the top part of this image; this is a result of imperfect stitching between two fields of view. This discontinuity should not have any effect on the data presented or the analysis of the data.

---

**Fig. S1.**
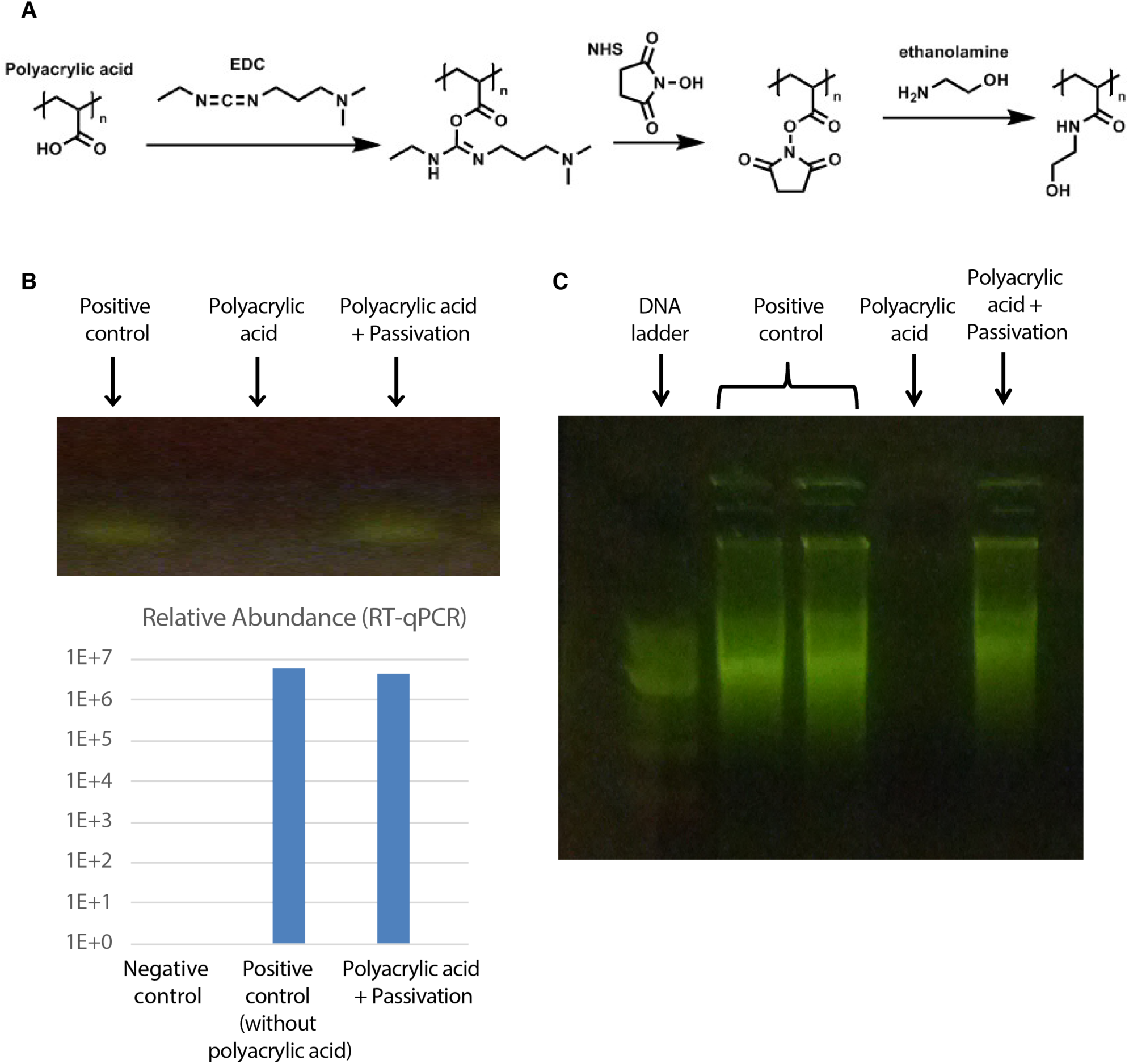
*In vitro* experiments demonstrating that polyacrylic acid, very similar to the polymer used in ExM, inhibits both the reverse transcription step and the rolling circle amplification step, two key enzymatic reactions of FISSEQ, but that the enzymatic activity can be restored with a passivation reaction that cancels out the charge of moieties on the polymer backbone. The passivation reaction (*95*), in which ethanolamine is reacted with carboxylic groups and converts them to amides with no charge (A), restores reverse transcription activity (B) and rolling circle amplification activity (C). These *in vitro* experiments were performed with 0.1-0.5% (w/v) polyacrylic acid, to mimic the concentration inside the expanded gel which is ∼0.2% (8.625% (w/v) sodium acrylate before expansion corresponds to 8.625/3.5^3 or ∼0.2% with expansion factor ∼3.5, and between ∼0.1% to ∼0.5% with expansion factors between 2.5 to 4). The passivation protocol was performed as in **Methods** section ‘Passivation’. We validated the passivation with a range of EDC concentrations, 50-150mM, that are higher than the polyacrylic acid concentration of ∼30mM (∼0.2% polyacrylic acid); however, we didn’t want the EDC concentration to be more than a few fold higher than the polyacrylic acid concentration as EDC can react with guanine in RNA (*96*), and therefore high concentrations of EDC may have undesired side effects. The reverse transcription reaction was performed with M-MuLV (enzymatics) according to the manufacturer’s protocol, with 1.2kb Kanamycin Positive Control RNA (Promega) as a template, and random primers. The rolling circle amplification reaction was performed with phi29 (enzymatics) according to the manufacturer’s protocol. The template used was the 55-base long CircLigase II ssDNA Control Oligo, after circularization with CircLigase II (epicentre) according to the manufacturer’s protocol.

**Fig. S2.**
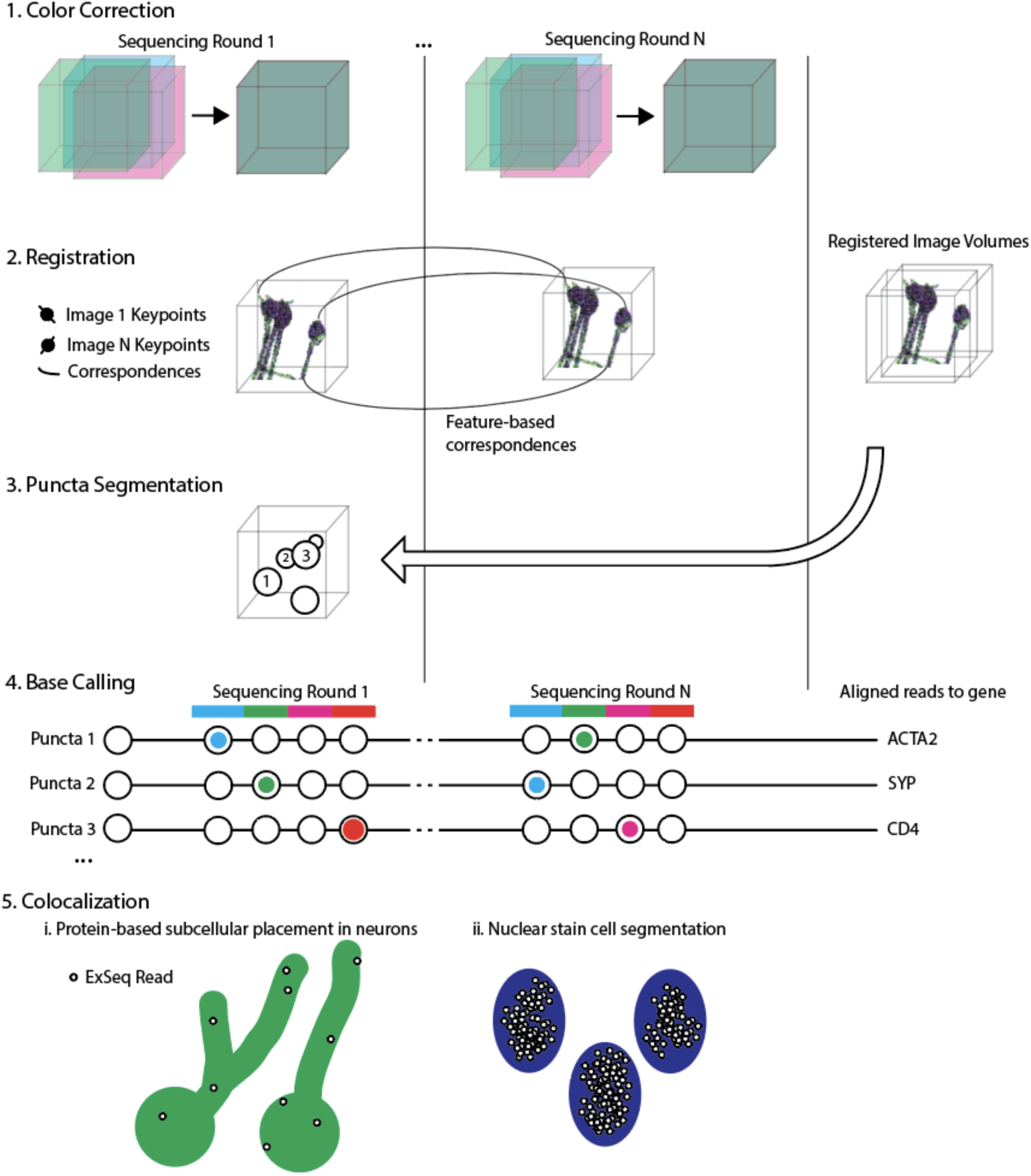
Schematic, for a single field of view, of the multi-step image processing pipeline that automatically processed 362 fields of view, across 6 ExSeq experiments (three untargeted, including 50 and 15 micron thick hippocampal slices, and one hippocampal culture, and three targeted, listed in **Table S5**), from volumetric images to spatially localized RNA reads. Each volumetric image for a given sequencing round is first rigidly color corrected to account for piezo stage drift and optical differences between color channels (1). Then (2), a precise registration (**Fig. S3**) is performed by detecting salient keypoints, describing each keypoint using a 3D SIFT descriptor, and discovering correspondences between keypoints in different sequencing rounds using the SIFT matching algorithm followed by RANSAC. The correspondences are then used to calculate an affine transformation. All the registered images are then combined to detect the amplicons; this puncta segmentation task is performed using watersheding (3). Next, each amplicon is described as a sequence of colors across rounds, by identifying the dominant color channel per round (4). These sequences are then aligned to either an *ex situ* library (untargeted ExSeq) or known barcode library (targeted ExSeq) to convert the color sequence into a gene identity. (5) The reads are then studied in space with the registered morphology or nuclear stain. In (5), all analyses were done in 3D, except for the 8 micron thick tumor tissue (Fig. 6), in which the reads were further studied in 2D. The entire MATLAB library to process ExSeq datasets from the microscope to spatial analysis of gene expression is publicly accessible at http://www.github.com/dgoodwin208/ExSeqProcessing. Specifically, the link includes a tutorial wiki with step-by-step instructions on how to run the pipeline and a tutorial set of targeted ExSeq data from the mouse visual cortex.

**Fig. S3.**
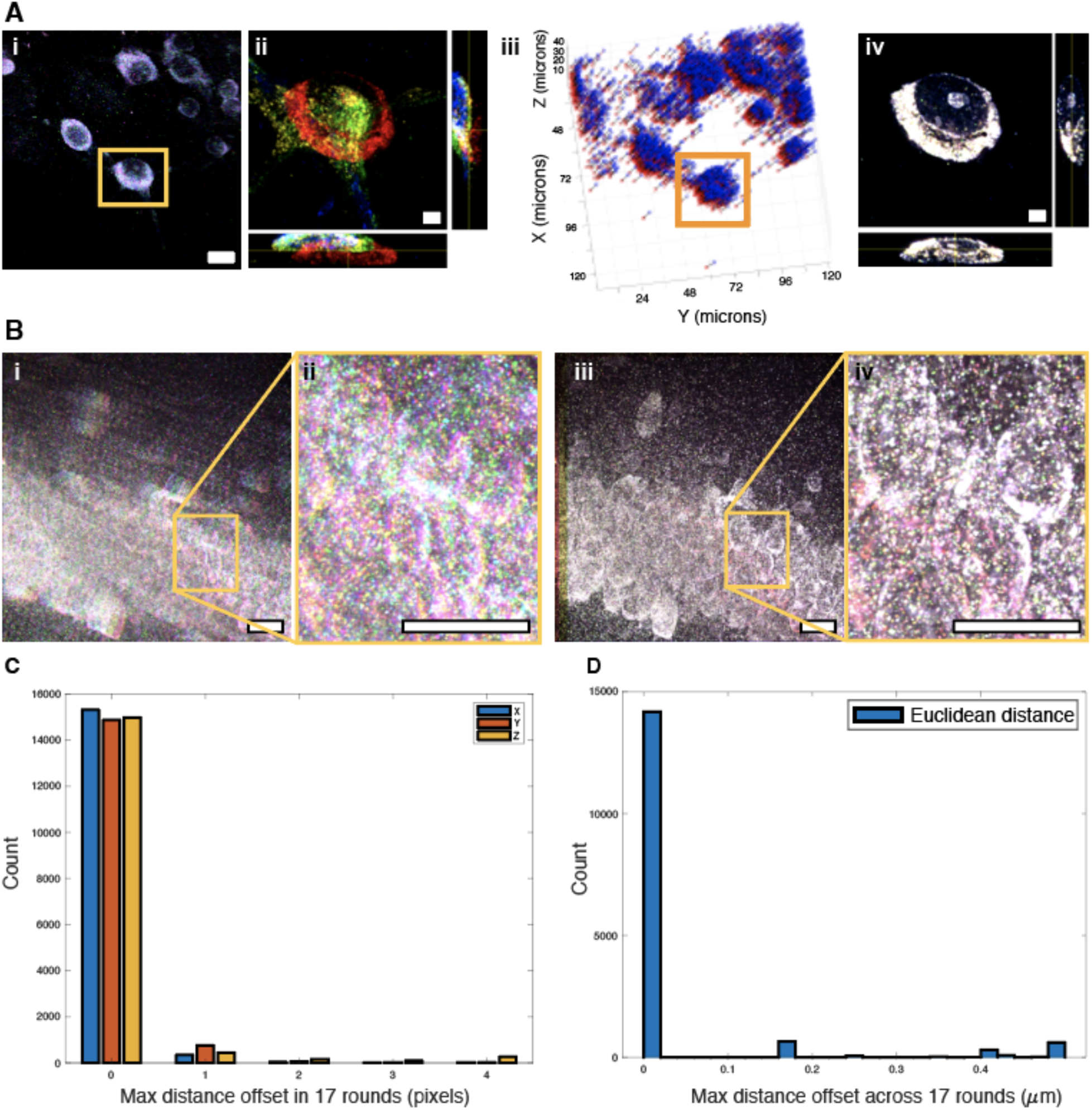
Image registration aligns puncta across 20 *in situ* sequencing rounds for untargeted ExSeq to within 1 pixel on average. Using a sparse neuron culture for demonstration purposes, a single round of sequencing is shown in (Ai), with the four SOLiD sequencing fluorescence channels shown (blue, green, red, magenta). Yellow box, region explored in depth in Aii. (Aii) Combined fluorescence channels of four sequencing rounds, for a set of cultured neurons, are shown as orthogonal views in four different colors: red, green, blue and yellow. Color indicates sequencing round, by summing all fluorescence channels in a given round, and white indicates overlap between all rounds. The image data is not registered. Features (3D SIFT descriptors, see **Methods**) are then calculated in each round and the correspondences to a reference round (the first sequencing round beyond the primer, chosen for its high image quality) are discovered via SIFT matching. (Aiii) 3,311 correspondence points between the reference round (shown in blue) and one of the 19 non-reference rounds (red) were calculated, with the specific neuron shown in (Ai) and (Aii) highlighted in the orange square. Axis labels are pre-expansion distances. (Aiv) After the affine warp is applied, the four sequencing rounds are in high agreement: colors are as in (Aii). (Bi) An example field of view from the intact mouse hippocampus showing six rounds (red, blue, green, cyan, magenta, yellow; all bases are given the same color so that the reader can focus on alignment) without registration, with zoomed in region showing approximately three cell bodies (Bii). White indicates overlap between rounds. (Biii-iv) The same regions as in Bi-ii, after registration. (C-D) To quantify the quality of registration, we calculated a normalized cross correlation of 15,943 subvolumes (each of size 41×41×19 pixels), randomly chosen across the imaged field of view (350×350×100 microns in size, post-expansion). (C) shows the maximum offsets, after registration, across the sequencing rounds for each subvolume in each dimension. The average of these maximum offsets per subvolume was 0.19 ± 1.52 pixels in X, 0.24 ± 1.62 pixels in Y and 0.16 ± 0.81 pixels in Z (mean ± standard deviation)). (D) Histograms of the maximum offsets shown in euclidean distance, with an average distance of 0.10 ± 0.5 microns (SD). Scale bars: (Ai,Bi-Biv) 13µm pre-expansion, (Aii, Aiv) 3µm pre-expansion.

**Fig. S4.**
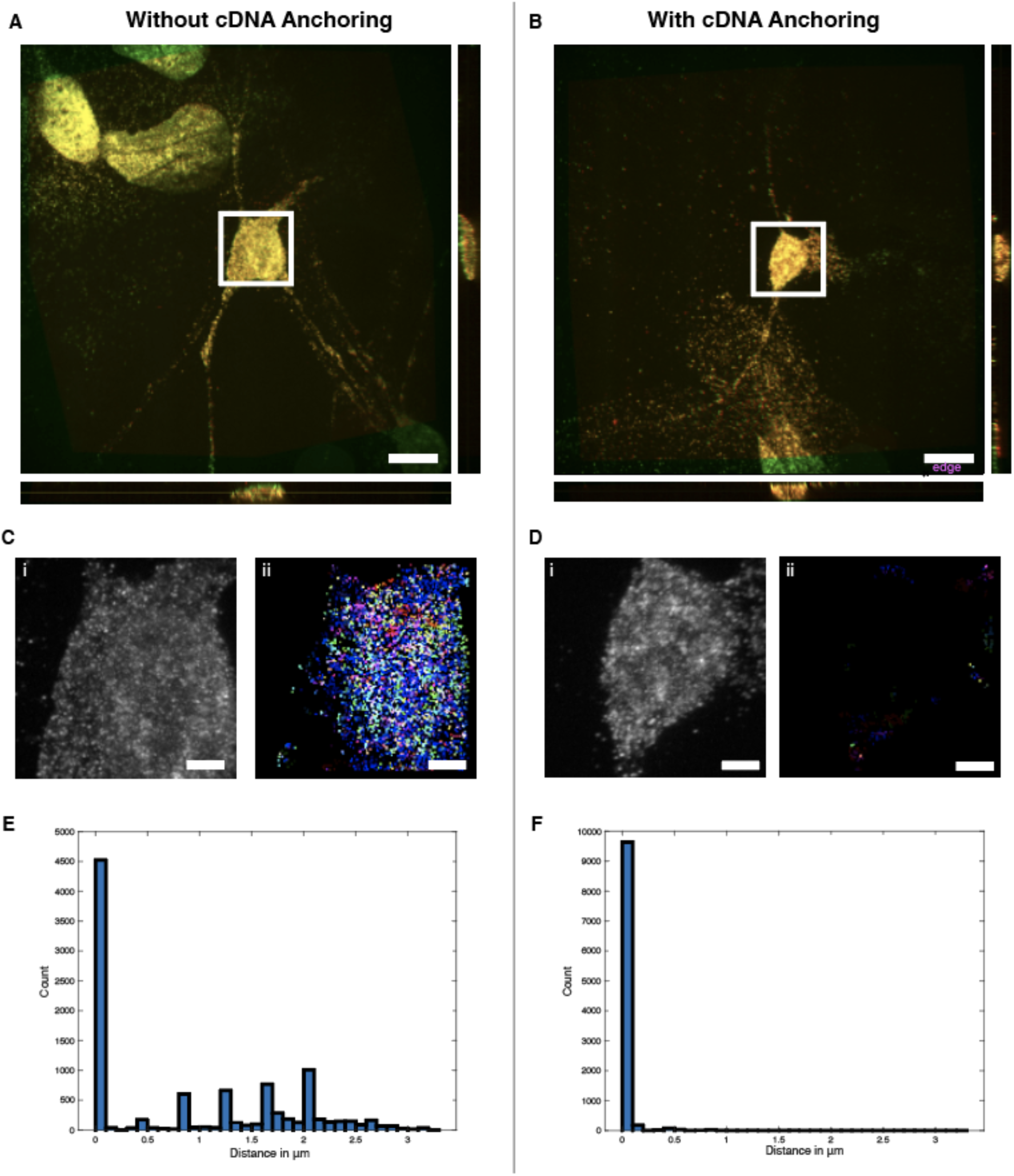
Anchoring of cDNA ensures precise spatial capture of cDNA location. In principle, without cDNA anchoring, the cDNA can move after the RNase digestion, as they are no longer linked to the gel in any way. To quantify this effect, we measured the location of the cDNA before the RNase digestion, and then 12 hours after RNase digestion, to create an extreme scenario in which the cDNA could, in principle, move quite a bit. The experiment was done both with and without cDNA anchoring, and performed in cultured hippocampal neurons. (A) BrdU antibody (see Methods at end of the figure caption, below) against cDNA before the RNAse step (green) and 12 hours after the RNase step (red), registered together (yellow overlap). (B) Same as (A), but the cDNA are anchored to the gel before the RNase step. Focusing just on a single soma, (C) and (D), a normalized cross-correlation method using 21×21×13pixel subvolumes from random locations within the volume shown was used to calculate cDNA drifts between the two imaging times. The locations of the 10,000 21×21×13pixel subvolumes were randomly chosen and required to have a minimum signal intensity to ensure sufficient signal-to-noise ratio for the autocorrelation calculation to calculate a correct offset. As shown in Cii and Dii, the direction and magnitude of the offsets are encoded in an RGB image. For each subvolume, the offset of the peak of the normalized cross-correlation is displayed in red for the X-direction offset (black=zero offset, bright red = offset of 10 pixels or 1.7µm), green for Y-direction offset (black=zero offset, bright green = offset of 10 pixels or 1.7µm) and blue for Z-direction offset (black=zero offset, bright blue = offset of 5 pixels or 2.0µm). (E) Only 52% of the subvolumes for the without-cDNA-anchoring condition are within 0.5µm, whereas 99% of the subvolumes match that criteria when cDNA anchoring was used (F). Scale bar: (A, B) 13µm pre-expansion, (C, D) 3µm pre-expansion. The BrdU staining protocol was performed as follows: 5-bromo-2′-deoxyuridine triphosphate was mixed with G, A, C to a concentration of 5mM. This mix was used instead of dNTPs in the reverse transcription reaction with final concentration of 0.25mM. After the reverse transcription, the RNase treatment, and the 12 hour wash in PBS, 5µg/ml anti-BrdU mouse antibody (Anti-Bromodeoxyuridine from mouse IgG1; Roche) in 1x PBS was incubated for 1 hour at room temperature. After washing with 1x PBS, 10µg/ml of goat anti-mouse cy5 antibody (abcam) in 1x PBS was incubated for 1 hour at room temperature. Finally the sample was washed again with 1x PBS. The cDNA anchoring was done as described in the **Methods** section ‘Library preparation for *in situ* sequencing’.

**Fig. S5.**
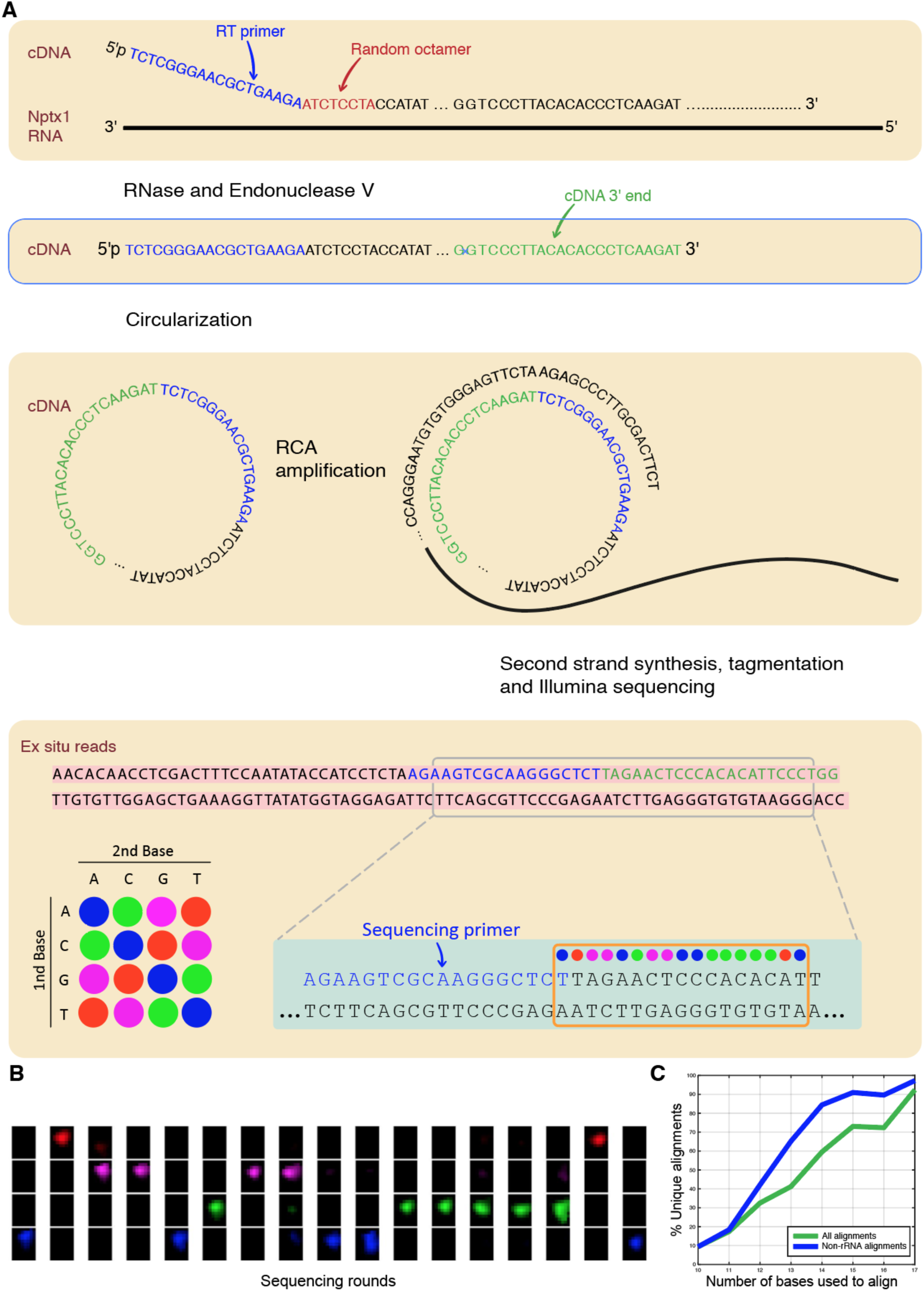
*Ex situ* library construction and *in situ* matching. (A) The random nature of untargeted reverse transcription priming, the cDNA size restriction, and the circularization of the cDNA, create unique molecular identifiers in the *in situ* sequenced region of the amplified cDNA, thus enabling later matching to *ex situ* reads. One sequenced region of the gene Neuronal Pentraxin 1 (*Nptx1*) is presented as an example. First, the cDNA is generated using random priming; the exact location of priming was reconstructed from the actual *ex situ* reads presented below. Inosine is included in the reverse transcription process, allowing generation of ∼100 base long cDNA fragments following Endonuclease V treatment. Next, RNase treatment and Endonuclease V cutting (at inosine) create cDNA fragments which are 95 bases long in this specific example, not including the constant region of the reverse transcription (RT) primer (the exact location of the cDNA 3’ end, reconstructed from the actual *ex situ* reads, is shown). cDNA circularization allows rolling circle amplification (RCA) with a RCA primer which is complementary to the constant region of the RT primer. Note that the cDNA anchoring to the hydrogel is not illustrated, for simplicity. After *in situ* sequencing, the amplified cDNA is extracted from the gel, the ss-cDNA is converted to ds-cDNA, and the ds-cDNA is tagmented and sequenced *ex situ*. The actual resulting *ex situ* reads are presented — note that in this case the paired-end reads give identical reverse complement sequences. To allow *in situ* and *ex situ* sequencing reads to be matched, we reconstruct the *in situ* read from the *ex situ* sequence; first we identify the sequencing primer sequence in the *ex situ* read, and then we convert the bases at the 5’ end of the sequencing primer to SOLiD color space according to the SOLiD 2 base encoding diagram (lower left). The sequences of colors resulting from the *ex situ* information (‘*ex situ* library’) are then used as a dictionary to align and directly match the *in situ* sequencing reads. (B) The sequences of colors resulting from the *ex situ* information (panel A) has one perfect match to the *in situ* sequencing of the gene *Nptx1*, sequenced in a dentate gyrus neuron in a 50 micron mouse hippocampus slice and shown on the left side of Fig. 1Giii. (C) The matching of the *in situ* reads to the *ex situ* library is detailed in the **Methods** section ‘Data Analysis - *Ex situ* and *in situ* sequence matching’. Overall, 92% of the matches, and 97% of the non-rRNA matches, are strictly unique in the sense that if a matching *ex situ* read is removed from *ex situ* library, the *in situ* read does not match to another *ex situ* read. Importantly, all *in situ* reads that are not strictly unique are removed (see **Methods** section ‘Data Analysis - *Ex situ* and *in situ* sequence matching’). This allows to explore sequence variations in mRNA, such as alternative splicing, using the longer *ex situ* matched reads.

**Fig. S6.**
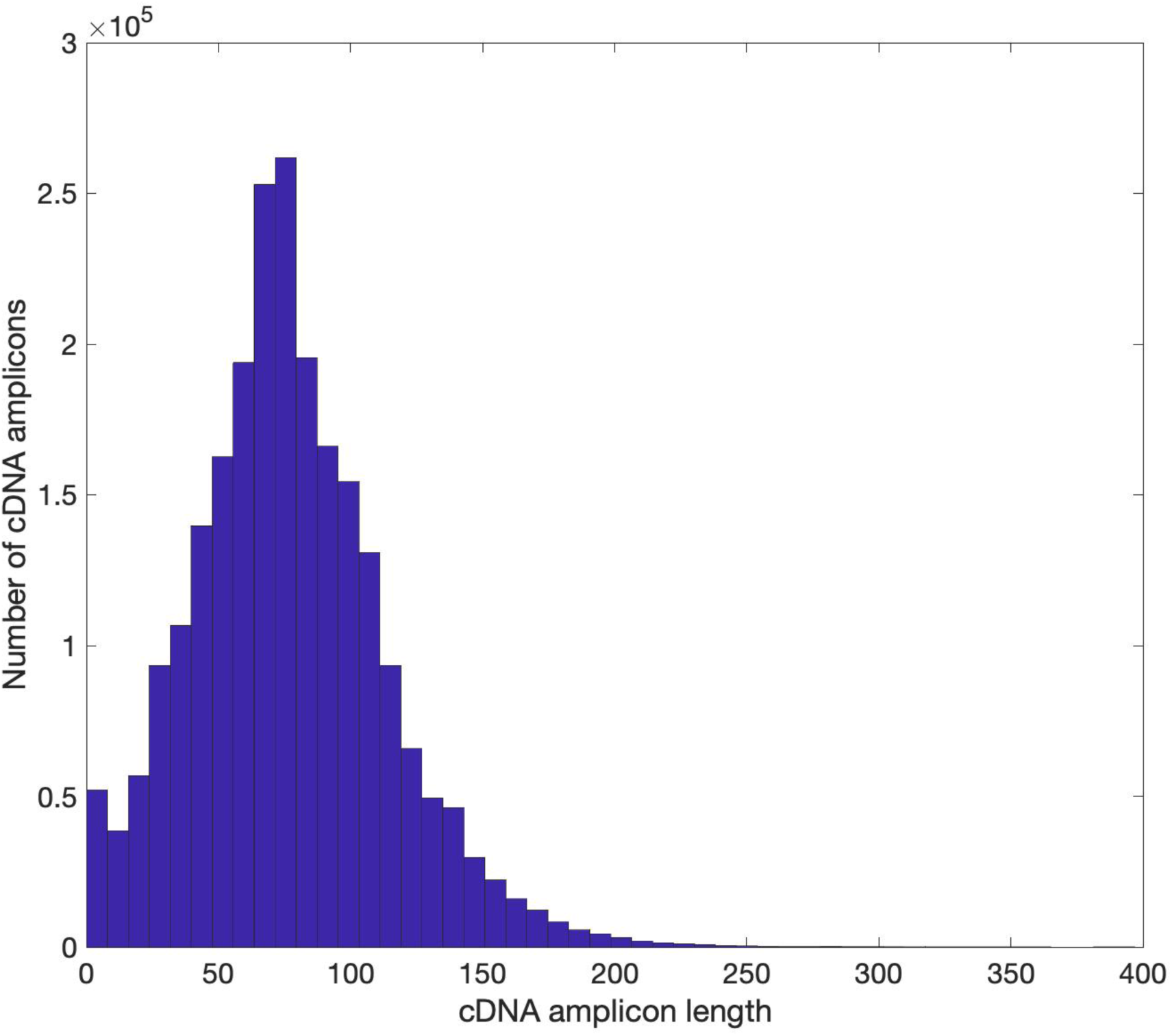
Histogram of cDNA amplicon lengths. The average cDNA amplicon length was 76.6 bases, with standard deviation of 36.6 bases, and maximal length of 397 bases. Methods used to generate this figure: The Illumina reads generated from *ex situ* sequencing typically contain several repeats of the cDNA fragment, as the size of the cDNA fragments was restricted to be ∼100 bases long (see **Methods**) and the Illumina reads were 300-600 bases long. To calculate the length of the cDNA fragments, the 300×2 paired-end reads from the 15 micron thick mouse hippocampal slice were merged using the software Pear (*126*) with default settings, resulting in the successful merge of 92.7% of the paired-end reads. The cDNA fragment repeats are flanked by the sequencing primer (introduced during reverse transcription), and therefore nucleotide-nucleotide BLAST version 2.8.1+ was used to detect all the sequencing primer locations in the merged reads. Default settings were used, with no limit on the number of possible matches. We note that setting a more stringent e-value compared to the default value of 10, for example an e-value of 0.05 which corresponds to a perfect match against the primer, had a minor effect on the resulting cDNA amplicon length distribution. The cDNA amplicon length was defined as the distance between two sequencing primer matches, using all the merged reads that contained more than one sequencing primer match; if more than two sequencing primer matches were present in a merged read, the average distance was calculated. Overall, 23.7% of the merged reads contained more than one sequencing primer match, and therefore were used in the calculation of the cDNA amplicon lengths. The calculation of the cDNA fragment lengths was also performed on the 150×2 paired-end reads from the 50 micron thick mouse hippocampal slice; however, because the Illumina reads were shorter, the merging of the paired-end reads was less successful (only 76.3% of the reads could be merged), and the number of merged reads with more than one sequencing primer match was only 2.2% of the merged reads. Therefore, the cDNA fragment length distribution from the *ex situ* sequencing of the 50 micron thick mouse hippocampal slice is not as representative as the data presented for the 15 micron thick mouse hippocampal slice.

**Fig. S7.**
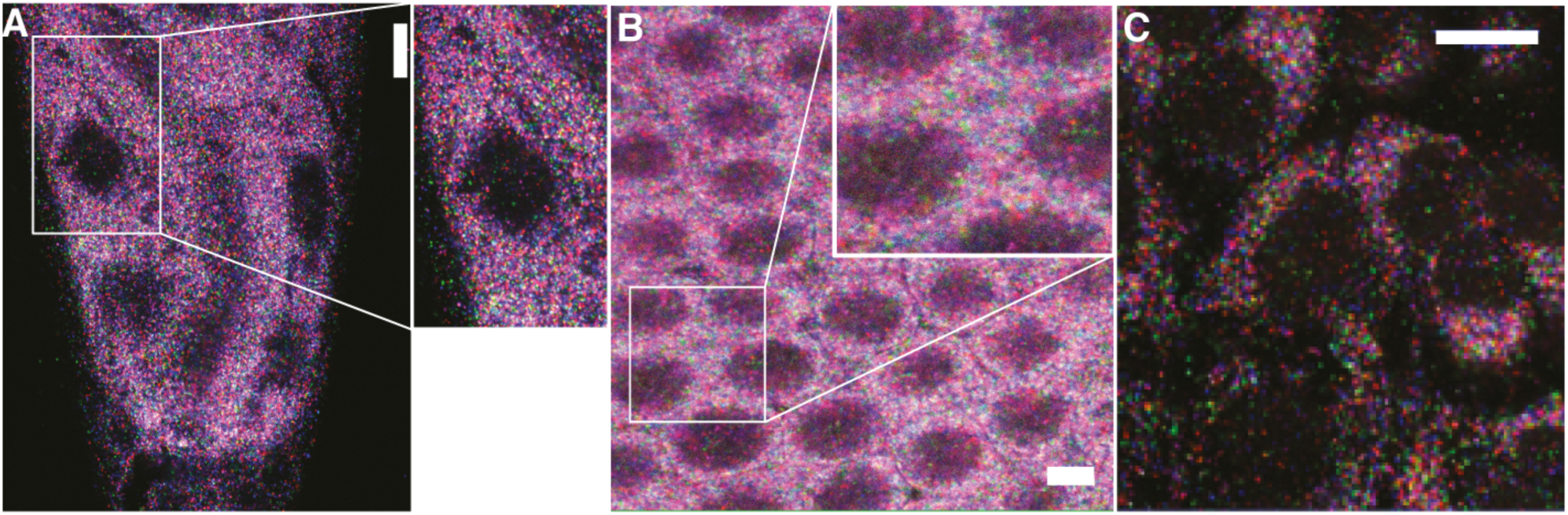
Demonstration of untargeted ExSeq with C. elegans (A), a Drosophila embryo (B) and the HeLa human cell line (C). The first in situ sequencing round is shown and the different colors (blue, magenta, green, and red) reveal the current base of the amplified cDNA (SOLiD sequencing was used). Scale bars: 20, 10 and 30 microns for panels A, B and C, respectively, in post-expansion (e.g., actual size) units. Methods used to generate this figure: Worm fixation and cuticle reduction was adopted from the published Bouin’s tube fixation protocol (*127*). The strain used in the figure was CZ1632: juIs76 [unc-25p::GFP + lin-15(+)] II. The strain was maintained at 20°C under standard conditions (*128*). The worms were collected from agar plates with M9 buffer (3g KH_2_PO_4_, 6g Na_2_HPO_4_, 5g NaCl, 1ml 1M MgSO_4_, water to 1 liter, sterilized by autoclaving) into a 15 mL tube. The tube was spun down at 1000g for 2 min, and the supernatant was replaced with 10 mL of fresh M9. The M9 wash step was repeated 2 more times. The worms were then transferred to a 1.5 mL tube and spun down to remove as much supernatant as possible without disturbing the worm pellet. The worms were placed on ice for 5 min. 1 mL of Bouin’s Fixative (0.46% picric acid, 4.4% paraformaldehyde, 2.4% acetic acid, 50% methanol, 1.2% 2-mercaptoethanol; as prepared in the published protocol), prepared fresh and pre-chilled to 4°C, was then added. The pellet was resuspended and mixed well. The sample was then placed on a tube rotator and mixed vigorously for 30 min at 25°C, followed by 4 hours of incubation at 4°C. The sample was then washed 3 times with 1mL Borate Triton β-mercaptoethanol solution (BTB; see recipe below); each time the sample was spun down, the supernatant was removed, the buffer was added and the sample was mixed thoroughly. BTB was prepared fresh using 1 mL 40x Borate Buffer Stock (3.1g boric acid, 1g NaOH, water to 50 mL), 1 mL 20% Triton X-100, 0.8 mL 2-mercaptoethanol, and 37.2 mL water. The sample was then further incubated three times, for 1 hour each, in 1 mL fresh BTB on a tube rotator at 25°C. Finally, the sample was washed six times: twice with 1 mL BT (1 mL 40x Borate Buffer Stock, 1 mL 20% Triton X-100, 38 mL water), twice with 1 mL 1x PBST (1x PBS, 0.5% Triton X-100), and twice with 1x PBS. The worms were permeabilized for 1 hour with 0.25% Triton X-100 in 1X PBS at 25°C. Drosophila larvae w1118 (BL#5905) were kindly provided by the lab of Aravinthan DT Samuel (Harvard University). Drosophila were raised in vials or bottles with standard yeast-containing medium at 22°C with alternating 12-h cycles of dark and light. HeLa (ATCC CCL-2) cells were cultured on CultureWell Chambered 16 wells Coverglass (Invitrogen) in D10 medium (Cellgro) supplemented with 10% fetal bovine serum (FBS) (Invitrogen), 1% penicillin–streptomycin (Cellgro), and 1% sodium pyruvate (BioWhittaker). Cultured cells were washed once with DPBS (Cellgro), fixed with 10% formalin in PBS for 15 min at 25°C, and washed three times with 1× PBS. Fixed cells were then stored in 70% ethanol at 4°C until use. ExSeq experimental procedures for the worms, Drosophila and HeLa cells were performed according to the following **Methods** sections: ‘RNA anchoring’, ‘Gelling, digestion and expansion’, ‘Re-embedding’, ‘Passivation’, and ‘Library preparation for *in situ* sequencing’. For the reverse transcription, instead of SSIV, M-MuLV (10U/µl; Enzymatics, cat. no. P7040L) was used for the worms and HeLa cells, whereas Maxima (10U/µl; Thermo Scientific, cat. no. EP0741) was used for Drosophila. Aminoallyl-dUTP was not included in the reverse transcription mix (and therefore the cDNA were not formalin-fixed), and the following primer sequence was used: /5Phos/ACTTCAGCTGCCCCGGGTGAAGANNNNNNNN. For the worms and HeLa cells, 2U/µl CircLigase II were used for circularization. To account for possible self-circularization of the primers, control samples with no reverse transcription enzyme were also processed for the worms, Drosophila, and HeLa cells, and as expected produced only a weak signal with hybridization probe after the library preparation. *In situ* sequencing was performed manually using the reagents and the enzymatic reactions outlined in the **Methods** section ‘Automated *in situ* sequencing’. Imaging was performed on a Zeiss Laser Scanning Confocal (LSM710) with Nikon 40X CFI Apo, water immersion with long working distance, NA 1.15 objective, and excitation light sources and emission filters as in (*23*).

**Fig. S8.**
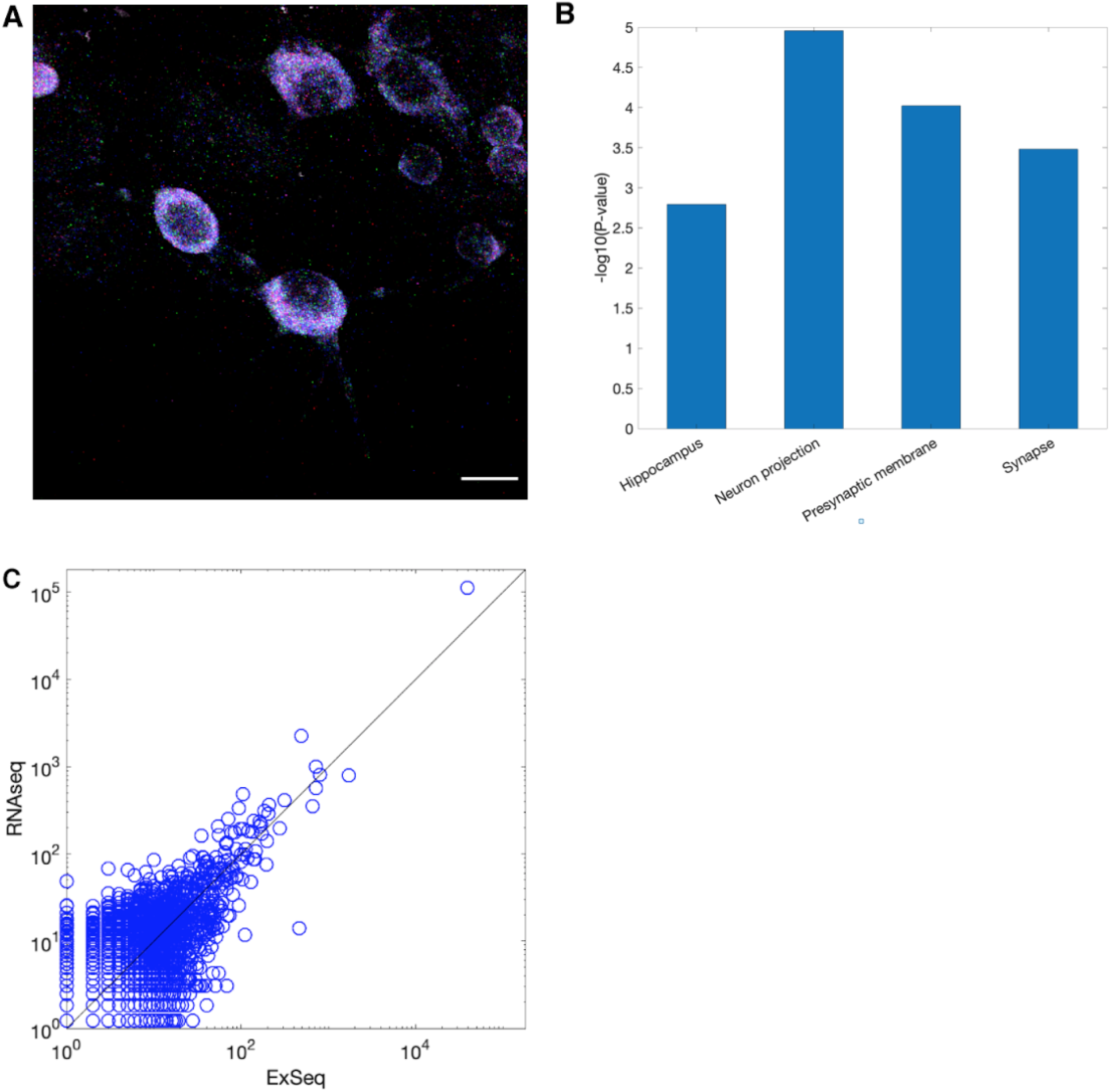
Untargeted ExSeq of hippocampal neurons in culture. (A) Maximum intensity projection of the first in situ sequencing round in a hippocampal culture. The different colors (blue, magenta, green, and red) reveal the current base of the amplified cDNA, in a SOLiD sequencing step. (B) Gene ontology analysis using the software DAVID (*110*) on all 127 expressed genes detected with ExSeq using 10 acquired FoVs (each of the size shown in A), in a hippocampal culture. Ontologies with low p-values are presented, revealing the expected functional enrichments for hippocampal neurons. See below the list of enriched GO terms as obtained by DAVID. (C) Agreement between the normalized expression levels of all well-annotated genes (RefSeq genes) using RNAseq, and ExSeq with full *ex situ* sequencing data, from the same hippocampal culture. The Pearson’s correlation between the log-transformed expression of RefSeq genes using ExSeq and using RNAseq is 0.621. Scale bar: (A) 13µm, pre-expansion.

For panel (B), the first few functional enrichment groups (i.e. the functions with the lowest p-values) as obtained by DAVID are given below.<colcnt=3>

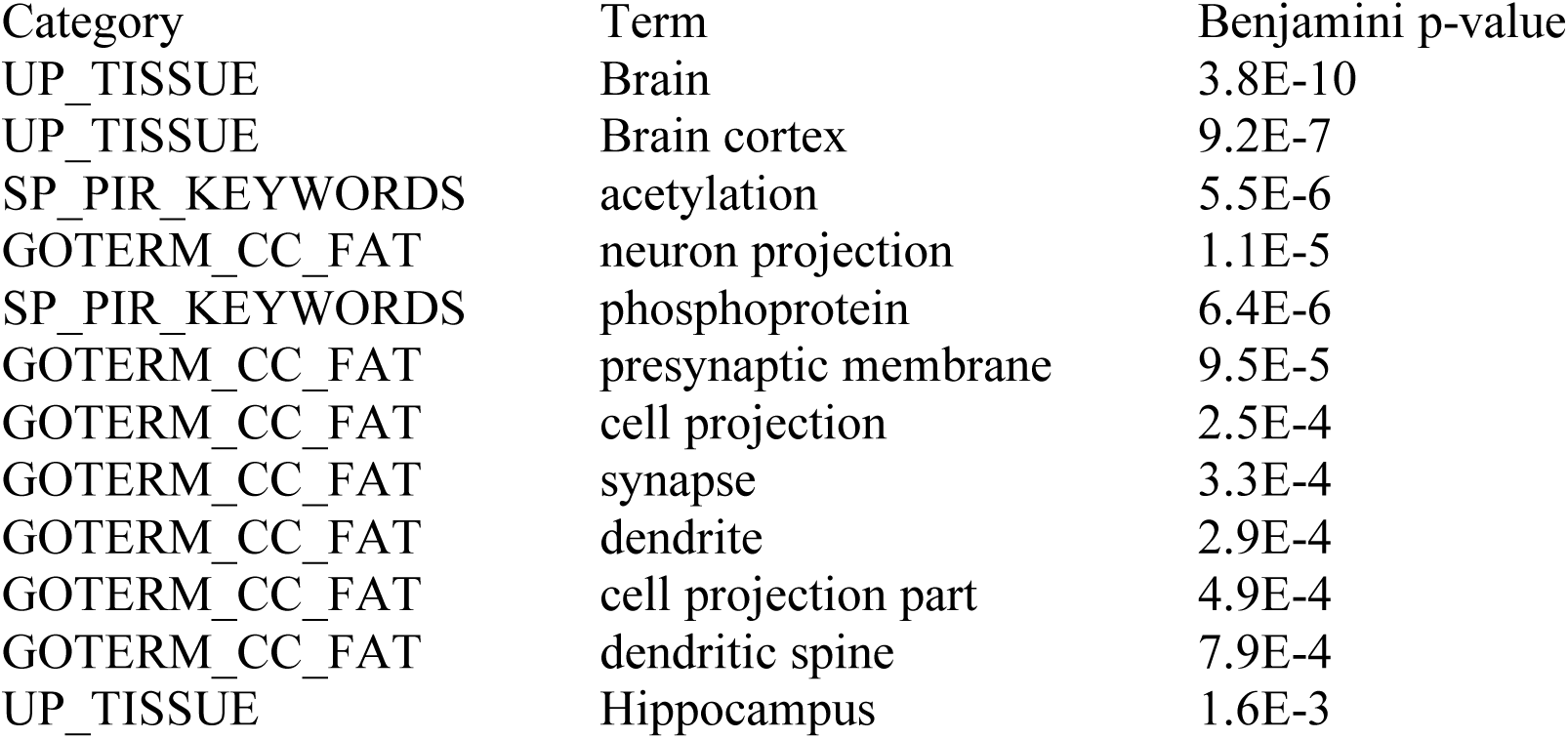

**Fig. S9.**
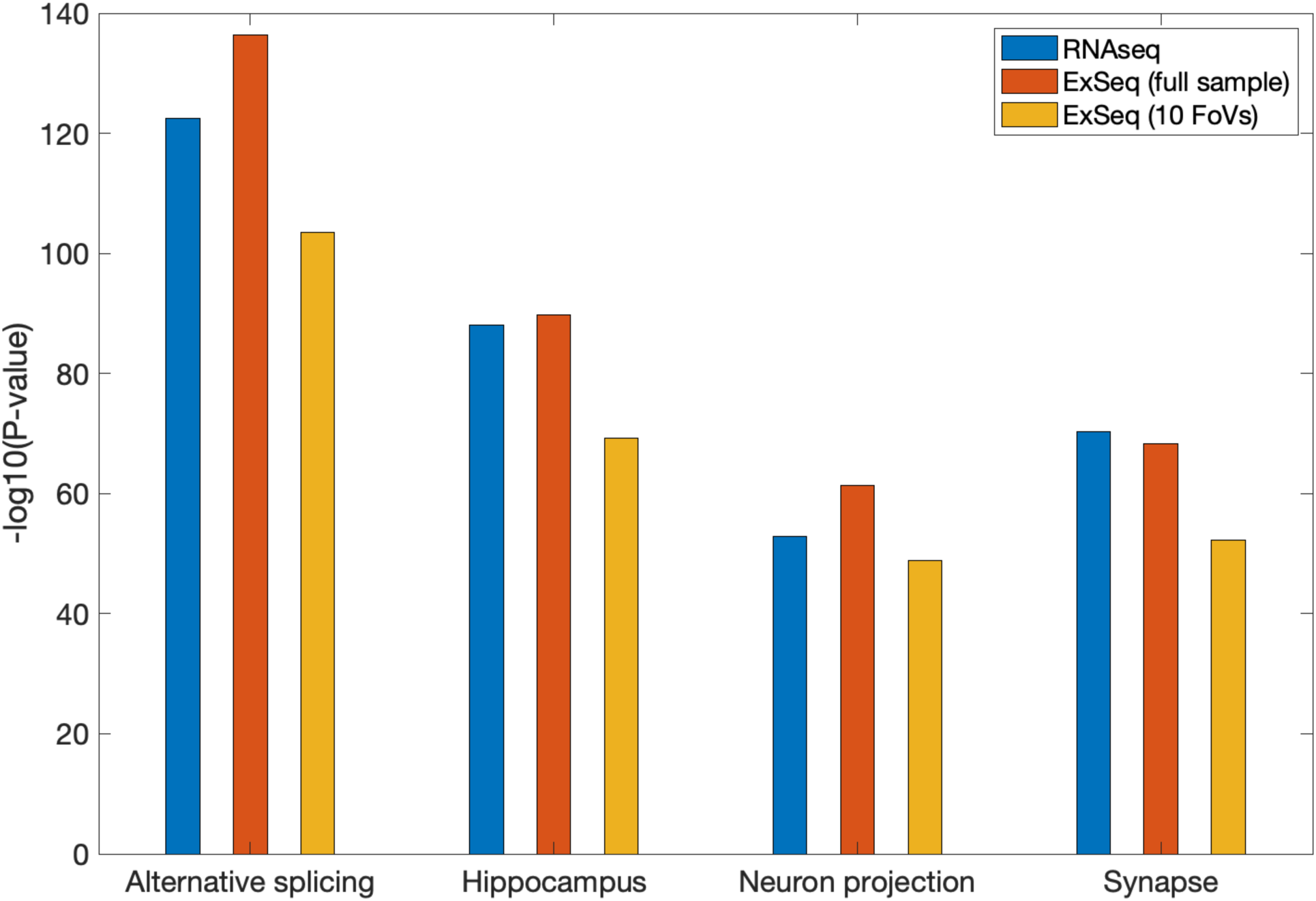
Gene ontology analysis using the software package DAVID (*110*) on the highly expressed genes detected with RNAseq, untargeted ExSeq with full *ex situ* sequencing data, and untargeted ExSeq using the *ex situ* data corresponding to the 10 acquired FoVs, in a 50 micron thick hippocampus slice. 3,039 genes were detected via ExSeq using the 10 acquired FoVs, and therefore the top 3,039 genes (sorted according to expression level) were analyzed in each one of the three analysis styles shown. Ontologies with low p-values are presented, revealing the expected functional enrichments for the hippocampus region. Importantly, the functionally enriched groups are common to the three tested conditions. The full list of the first 25 functional enrichment groups (i.e. the functions with the lowest p-values) as obtained by DAVID for each dataset is given in **Table S6**.

**Fig. S10.**
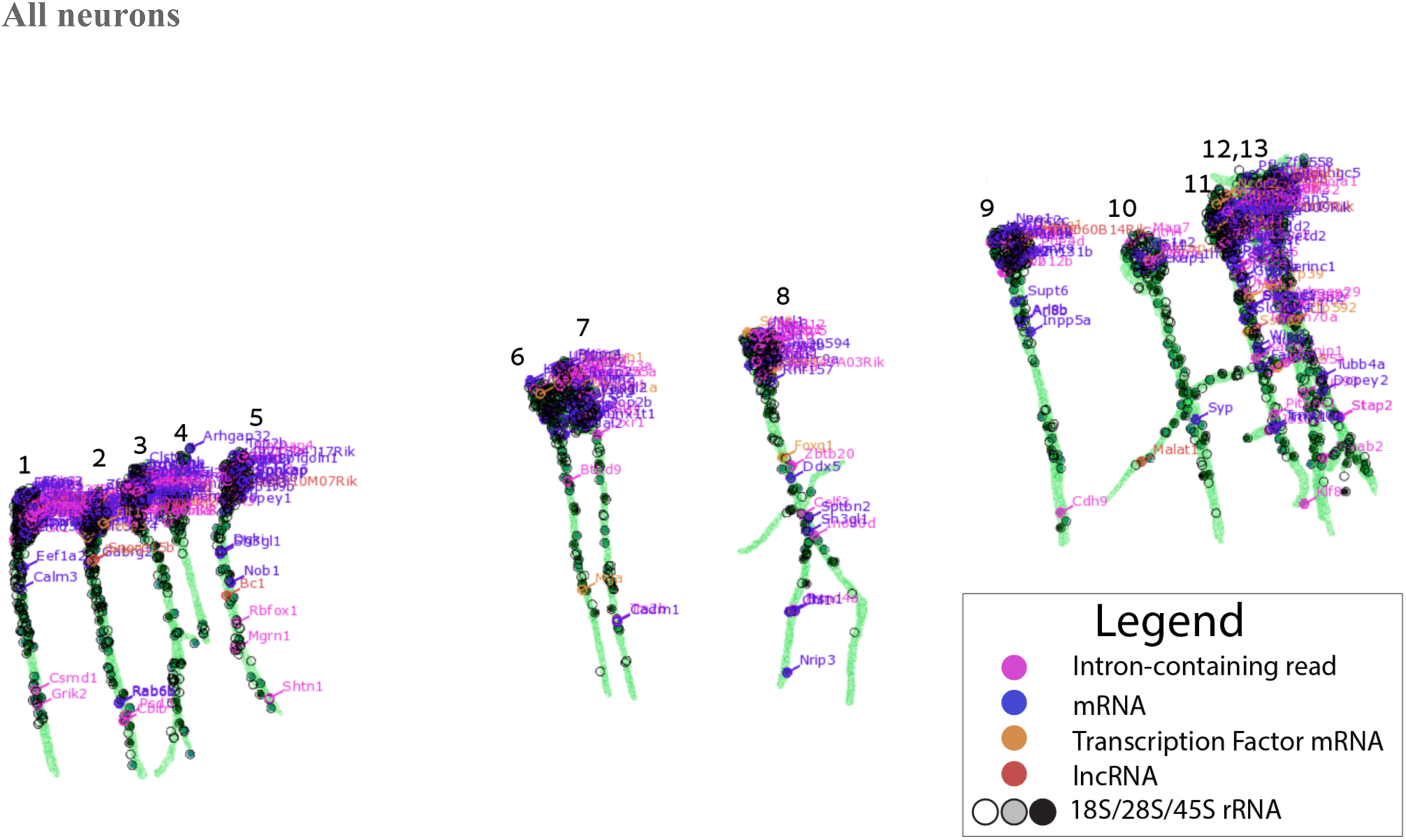
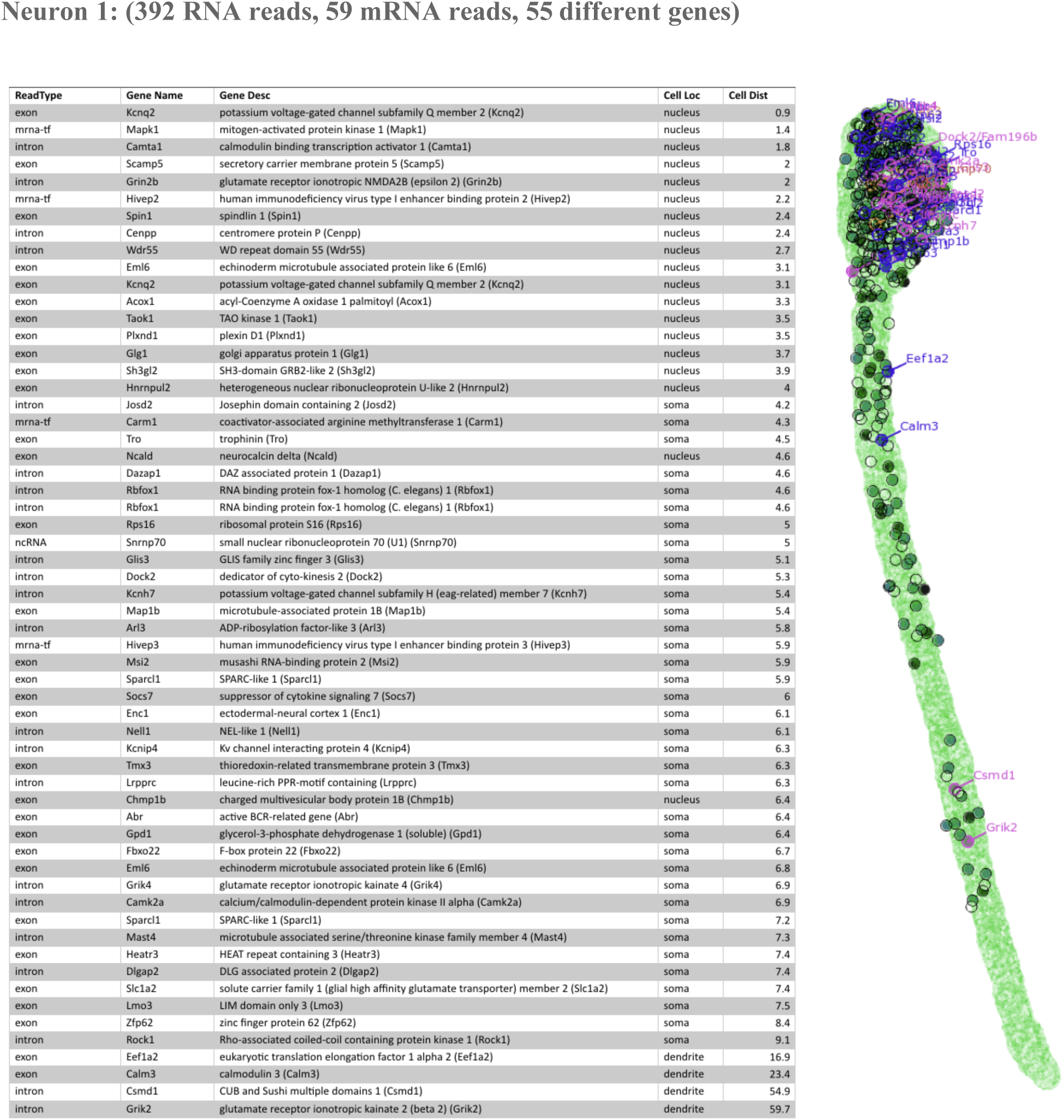
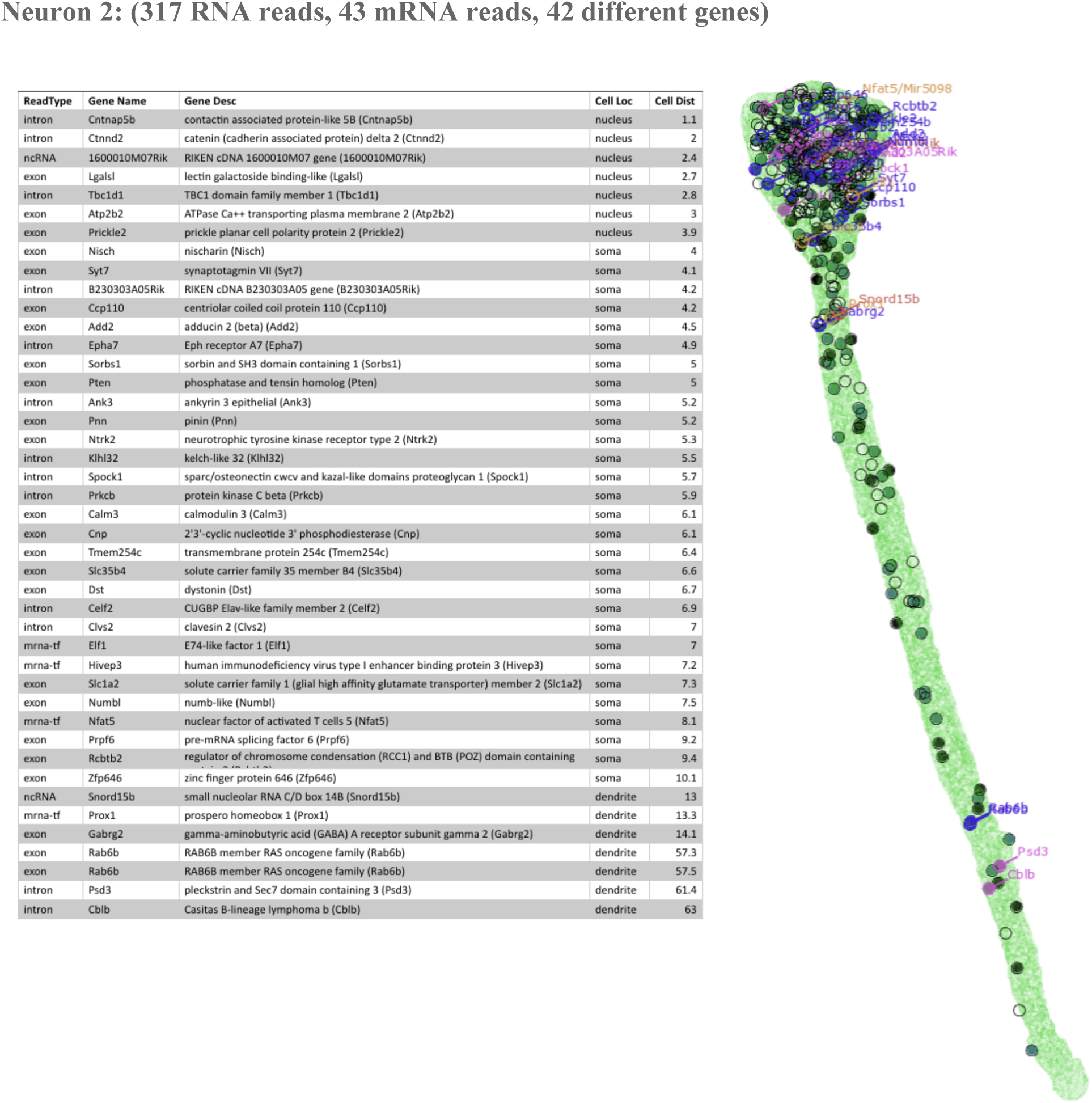
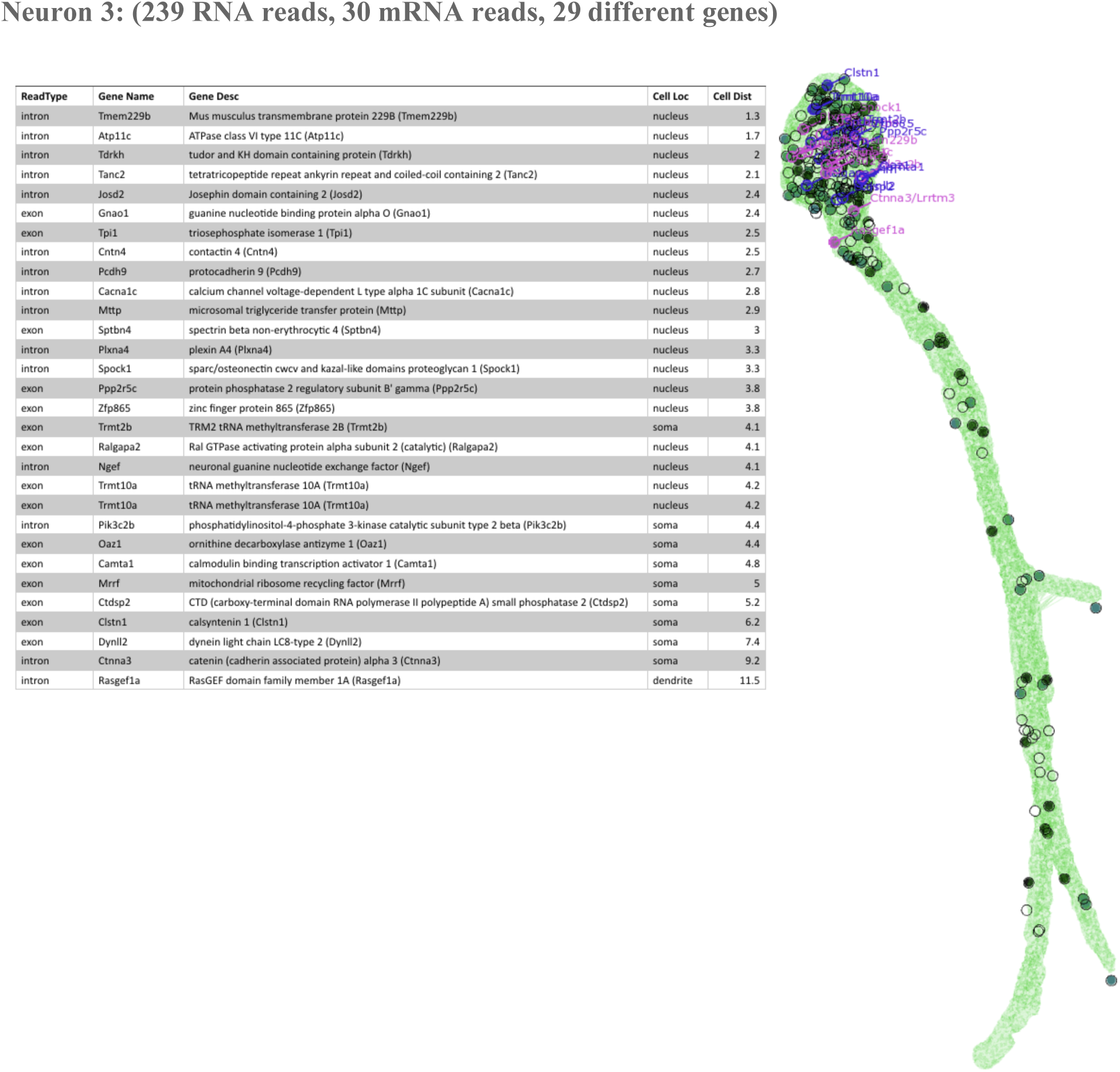
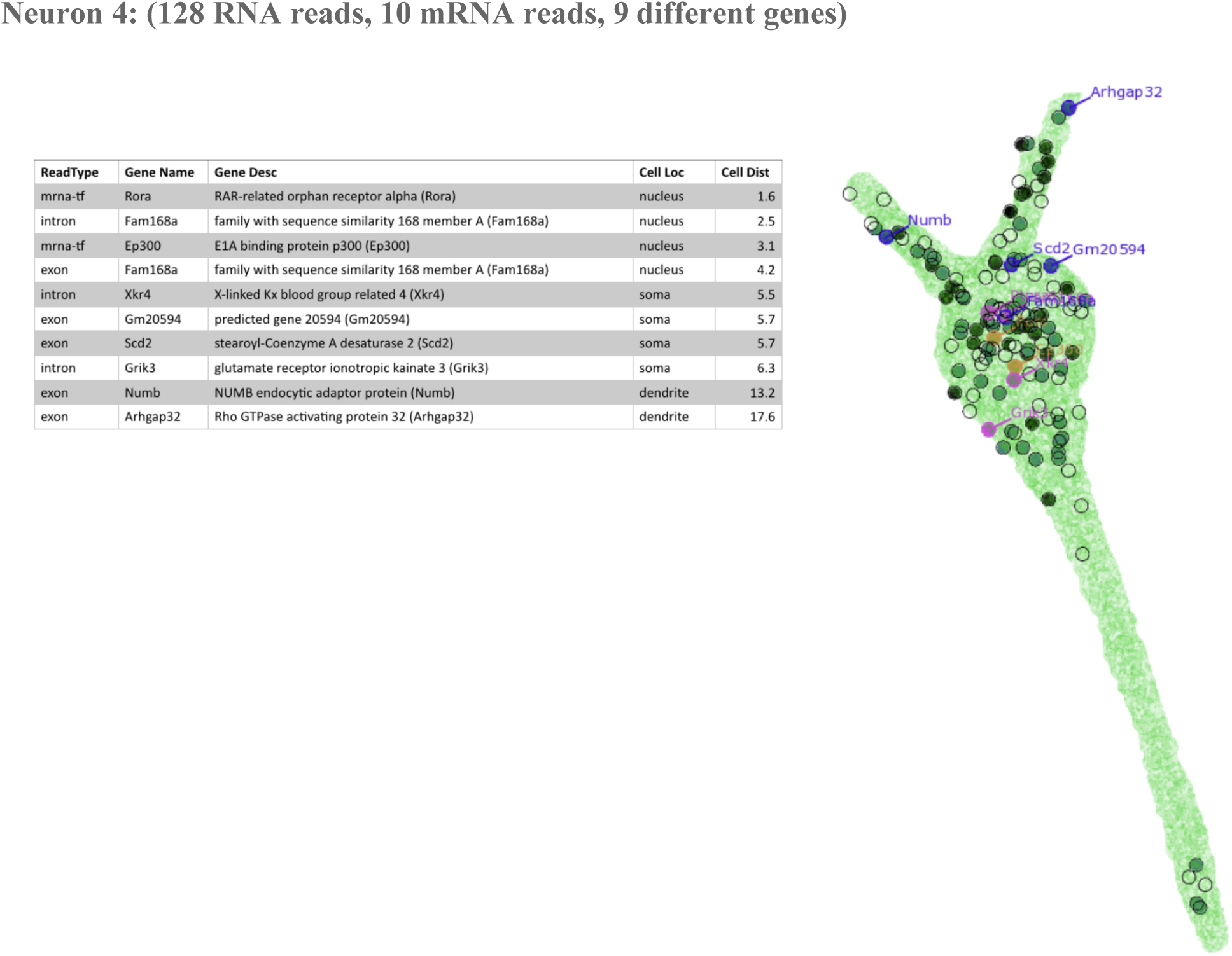
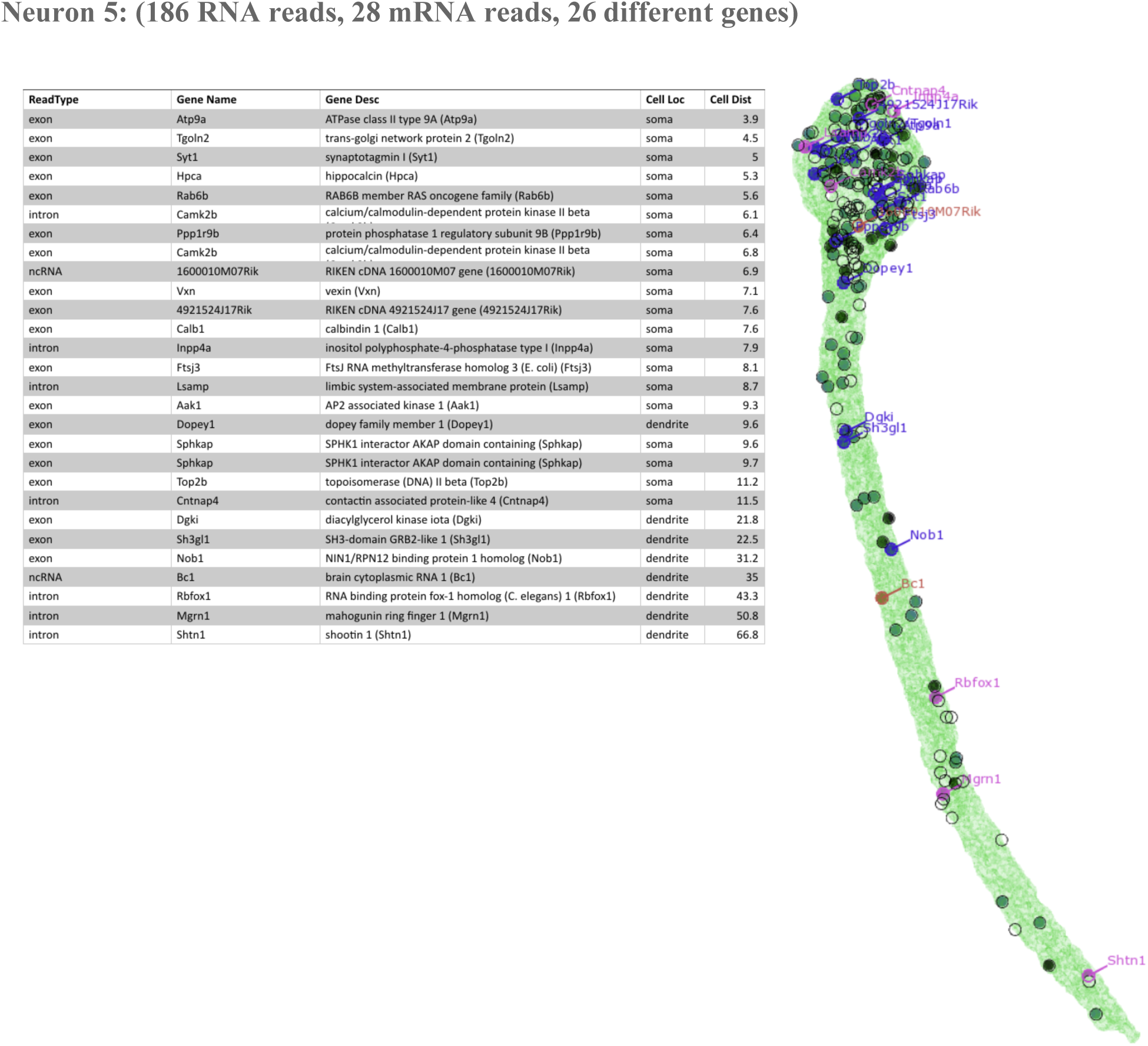
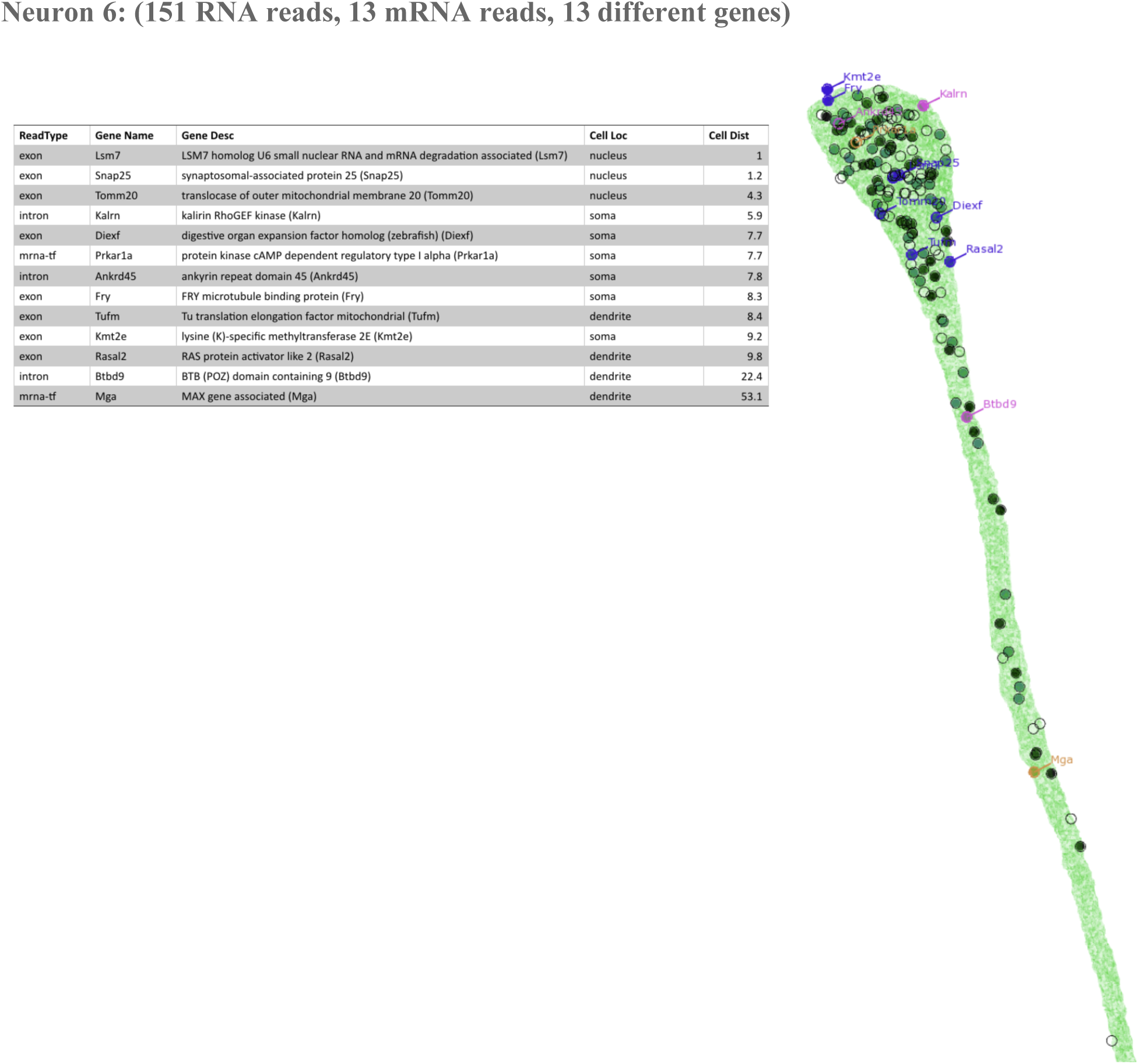
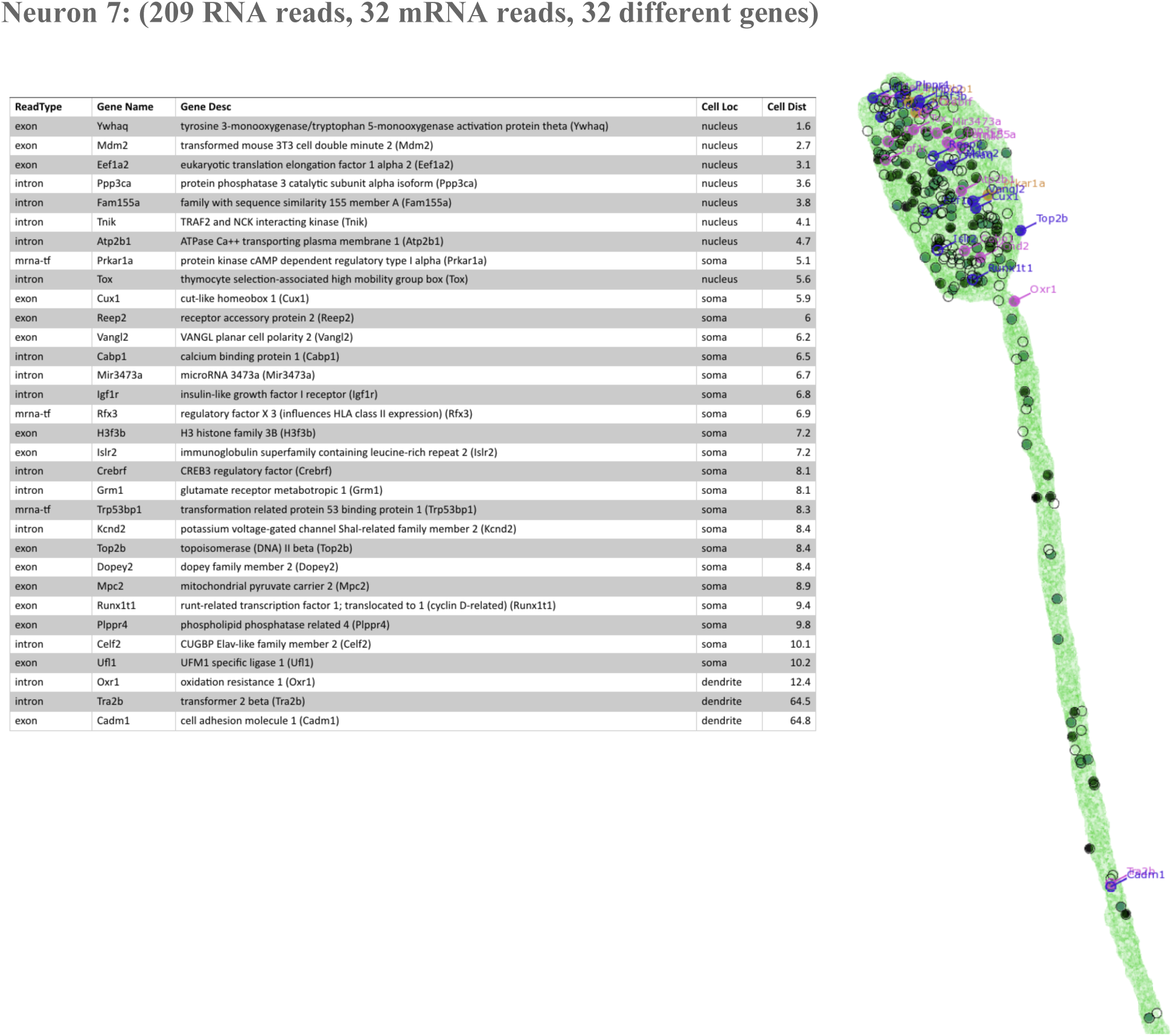
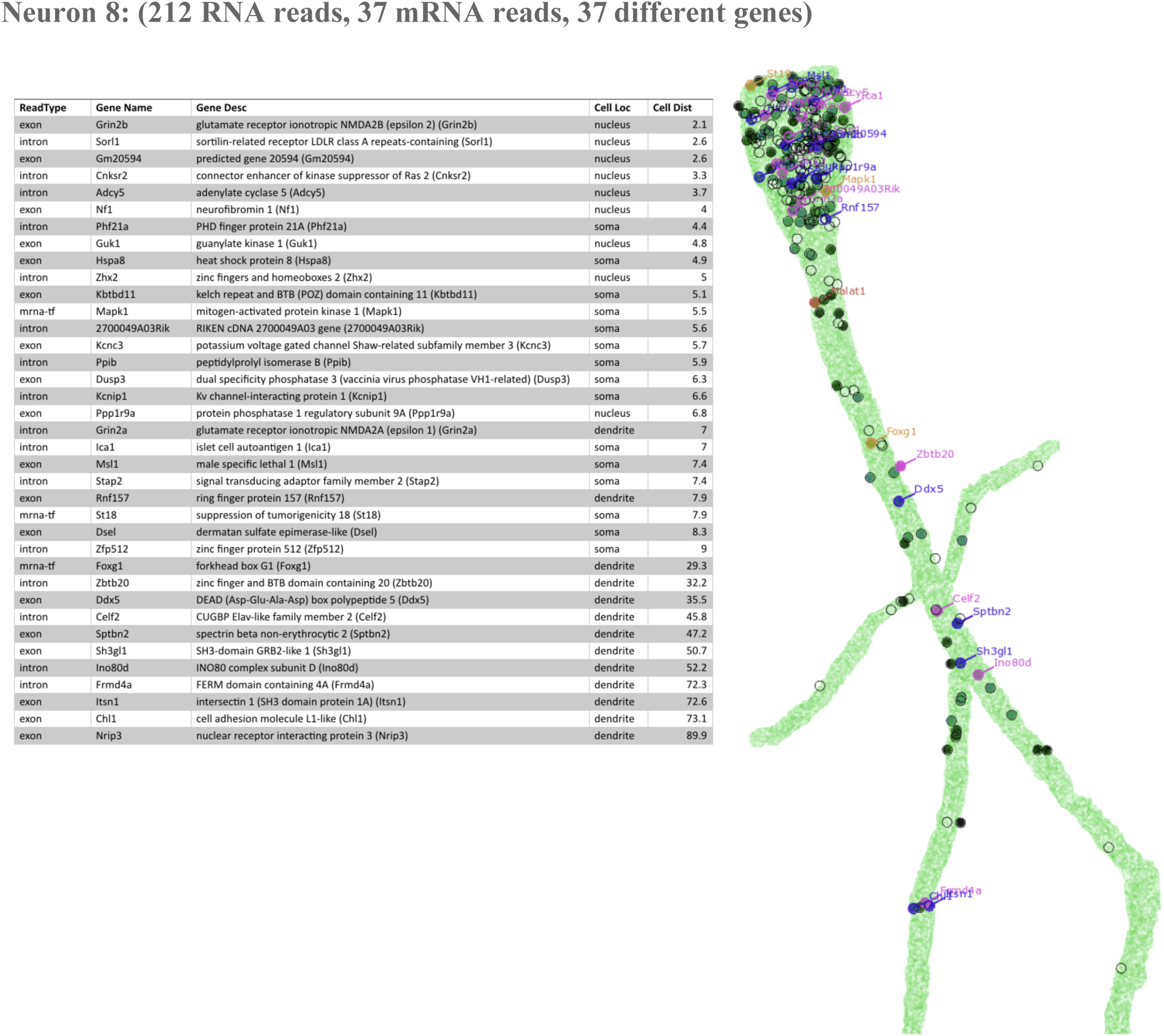
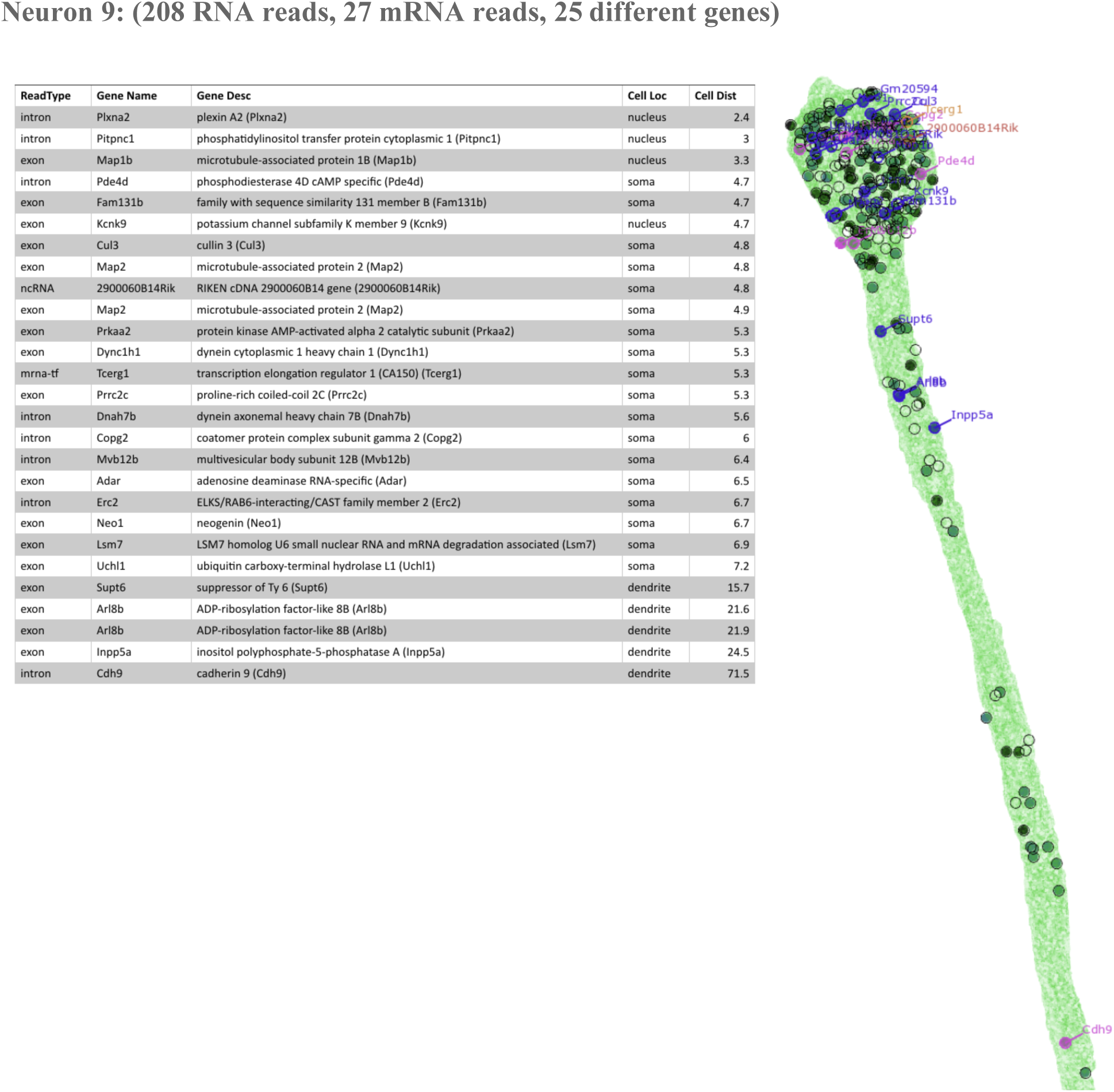
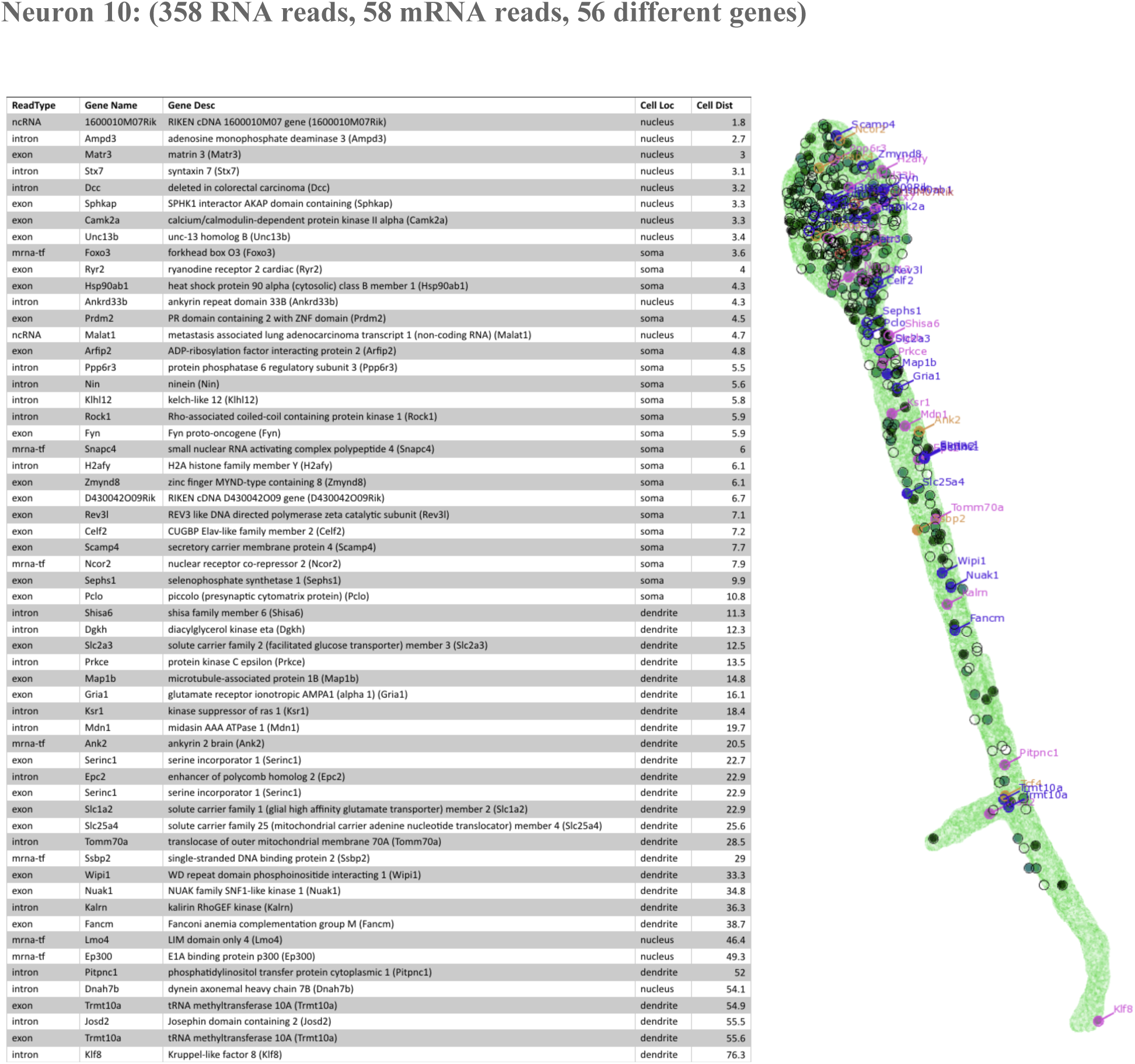
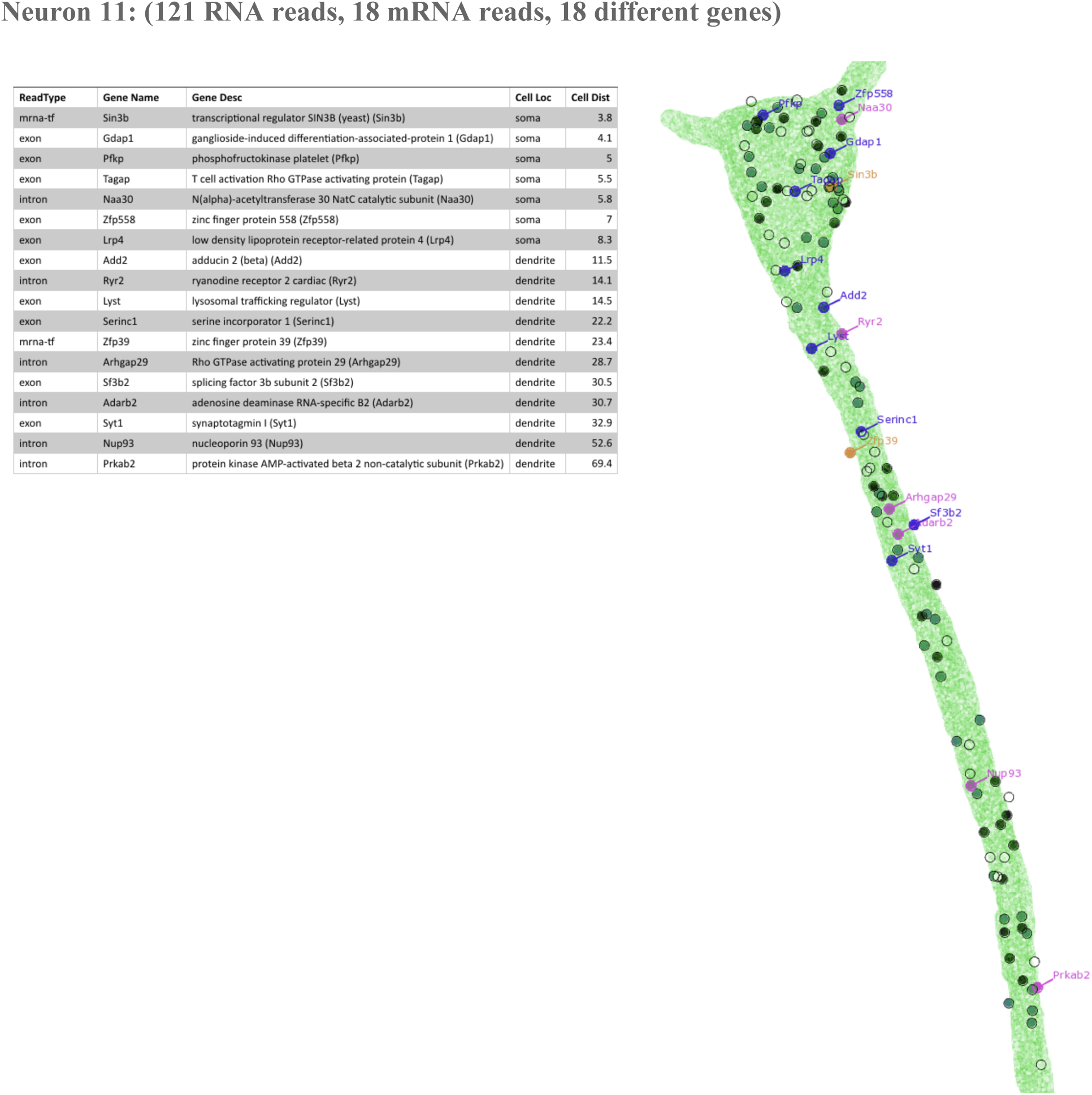
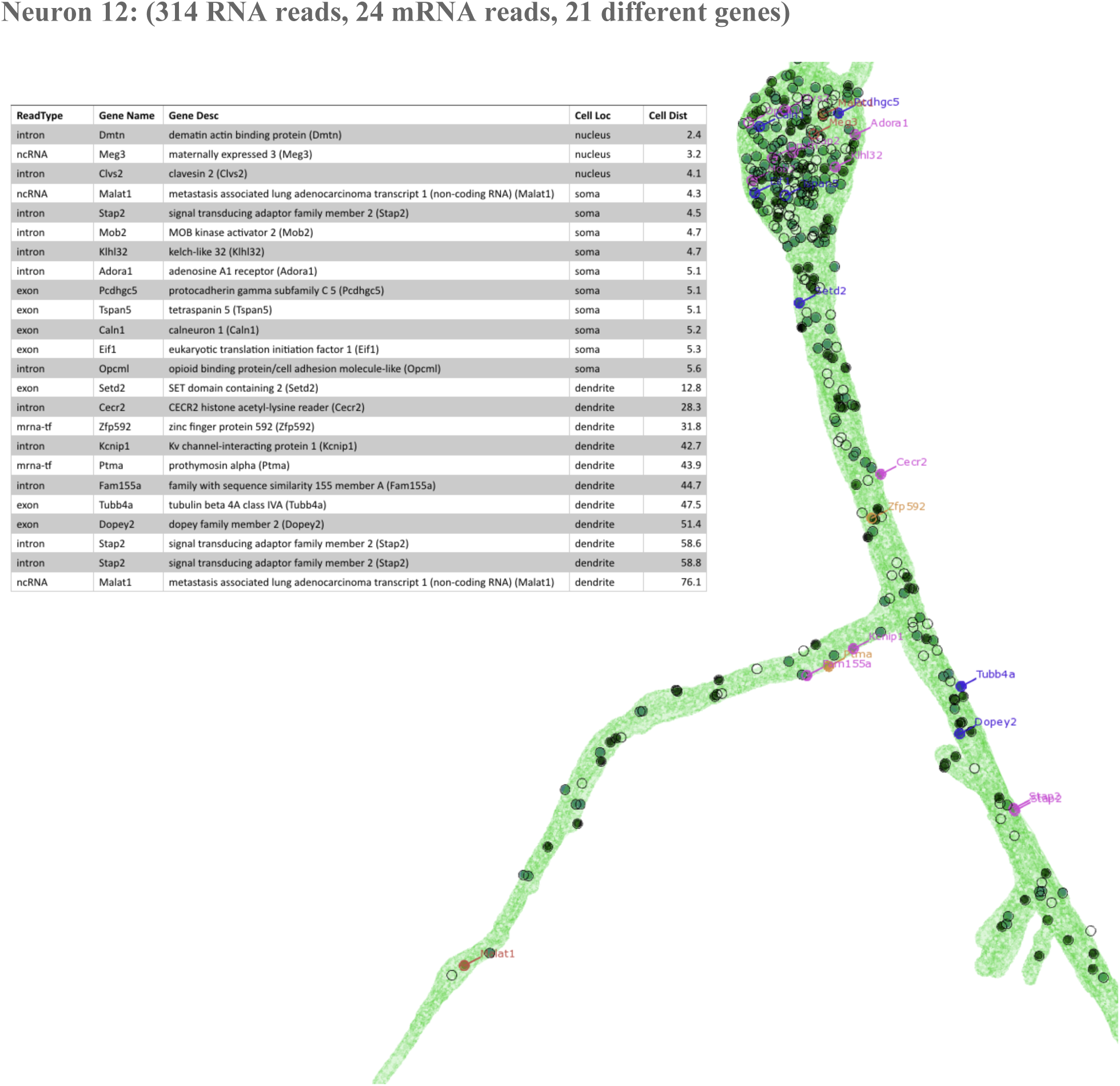
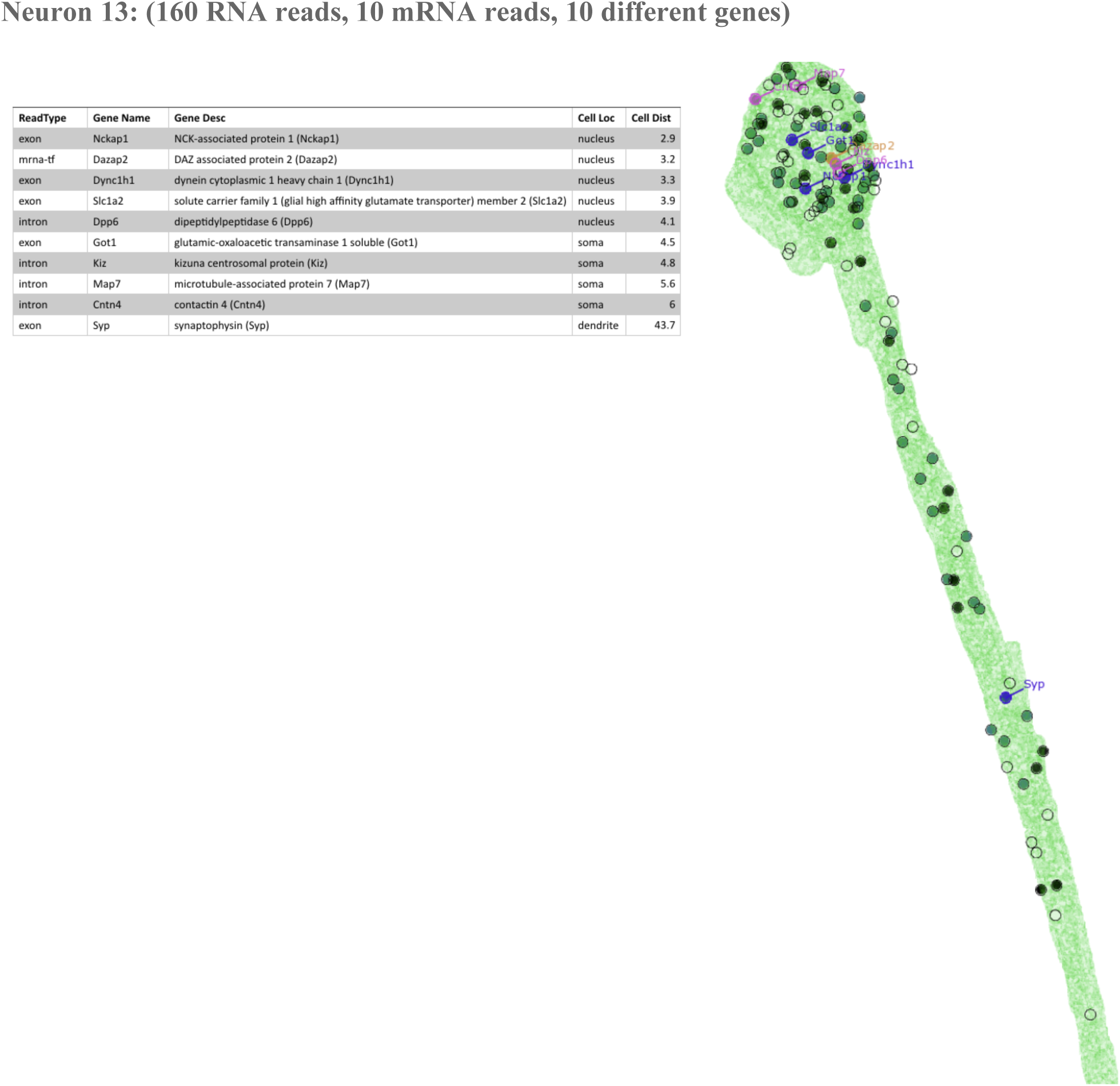
Complete list of the transcripts detected in each hand-traced neuron, not including rRNA. Each neuron is presented separately, numbered from left to right. For each neuron, the transcripts are listed according to their position from top (soma) to bottom (end of dendrites). Each ExSeq read is assigned (‘Cell Loc’ below) into the dendrite, soma, or nucleus (see Methods section ‘Image Processing - 3D Tracing’), and an Euclidean distance (‘Cell Dist’ below) is measured from the centroid of the nucleus (in microns, pre-expansion).

**Fig. S11.**
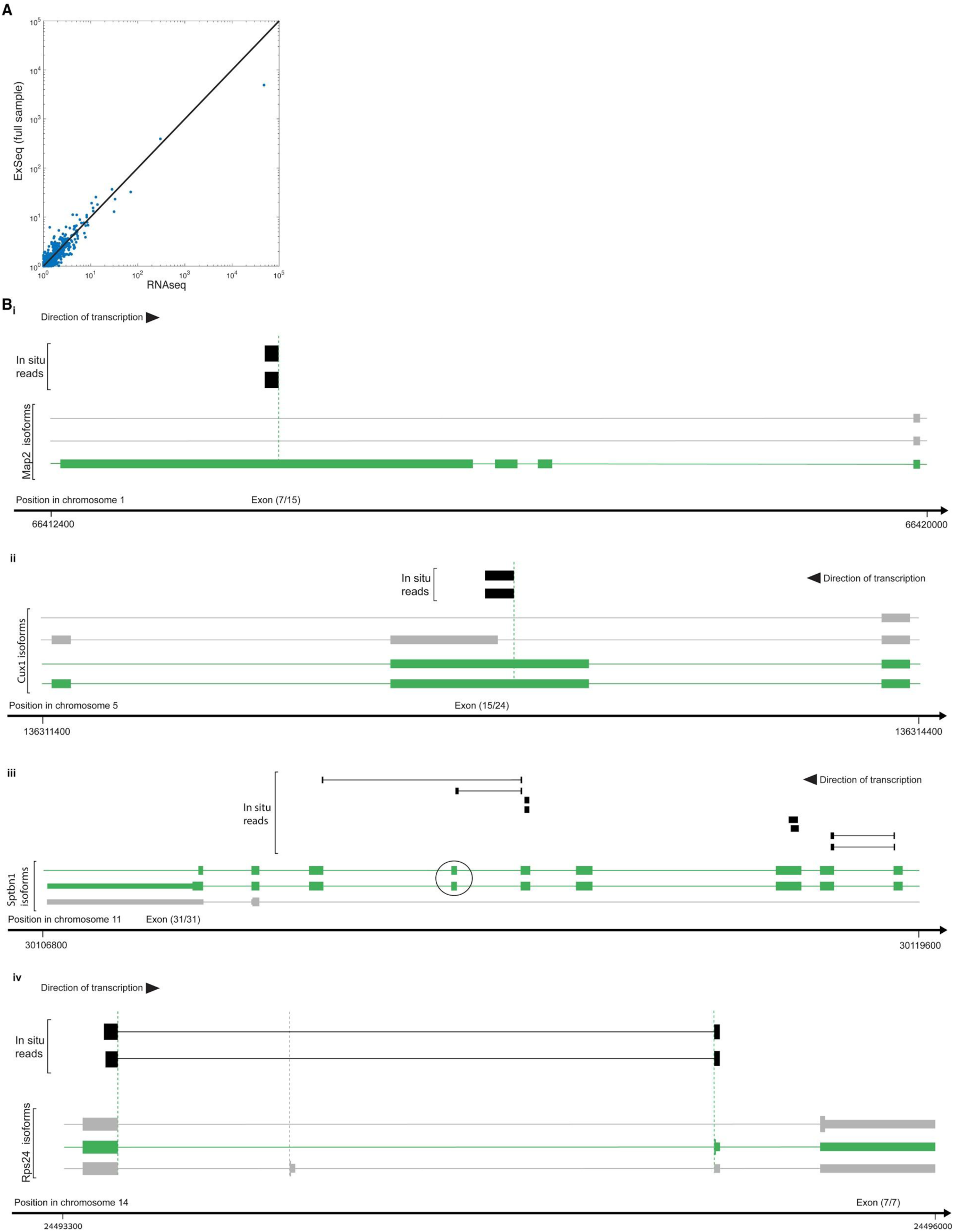
Untargeted ExSeq mapping of alternative splicing events in neurons. (A) ExSeq resolves alternative splicing isoforms with high correlation to those seen with standard RNAseq. For each RefSeq gene identified in untargeted ExSeq, the counts of the highest frequency isoform divided by the counts of the second highest frequency isoform are calculated for ExSeq (y-axis) and for RNAseq of the adjacent slice (x-axis). (B) Examples of alternative splice events detected *in situ*, shown for the genes Microtubule Associated Protein 2 (Map2) (i), Cut Like Homeobox 1 (Cux1) (ii), Spectrin Beta, Non-Erythrocytic 1 (*Sptbn1*) (iii) and Ribosomal Protein S24 (Rps24) (iv). For each sub-figure the genomic coordinates are displayed at the bottom, the known gene isoforms (RefSeq annotations) that align to that genomic region are in the middle (one row for each isoform), and the aligned *in situ* reads are at the top (each row corresponds to one sequenced puncta). For each sub-figure, the relative position of one of the exons is displayed in the following format: x/y, i.e. exon number (starting from the 5’ of the mRNA) / total number of exons. Thick lines for the known gene isoforms and *in situ* reads represent the sequences matching to the genomic coordinates at the bottom, whereas thin lines represent gaps in the alignment due to intronic regions. For the mRNA, thin lines represent introns, medium-thick lines represent untranslated regions, and fully thick lines represent coding regions. For all four genes, the *in situ* reads are matching the structure of specific mRNA isoforms (shown in green) and not other isoforms (grey), therefore revealing the expressed alternative splicing isoforms. In addition, for *Sptbn1*, the top *in situ* read possibly reveals a previously uncharacterized exon skipping splicing event, as the exon marked in a circle is not present in the read whereas the two flanking exons are present.

**Fig. S12.**
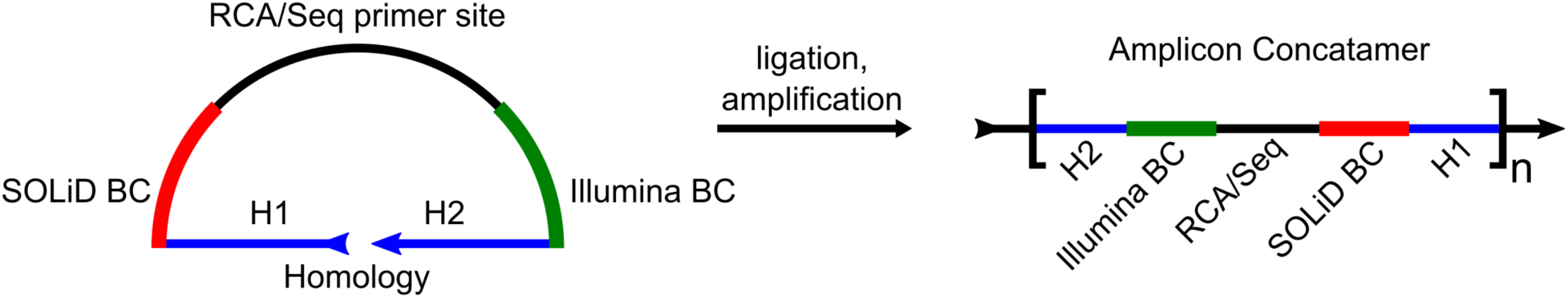
Architecture of padlock probes for targeted ExSeq. Targeted ExSeq padlock probes consist of four key elements: (1) a 32 nt homology region, split into two 16 nt halves (H1, H2) on the 5’ and 3’ ends of the padlock probe; (2) a constant RCA/Sequencing primer site on the backbone; (3) a barcode (‘BC’) sequence for readout with SOLiD sequencing chemistry; and (4) a barcode sequence for readout with Illumina chemistry. After ligation and rolling circle amplification, the amplicon concatamer is formed.

**Fig. S13.**
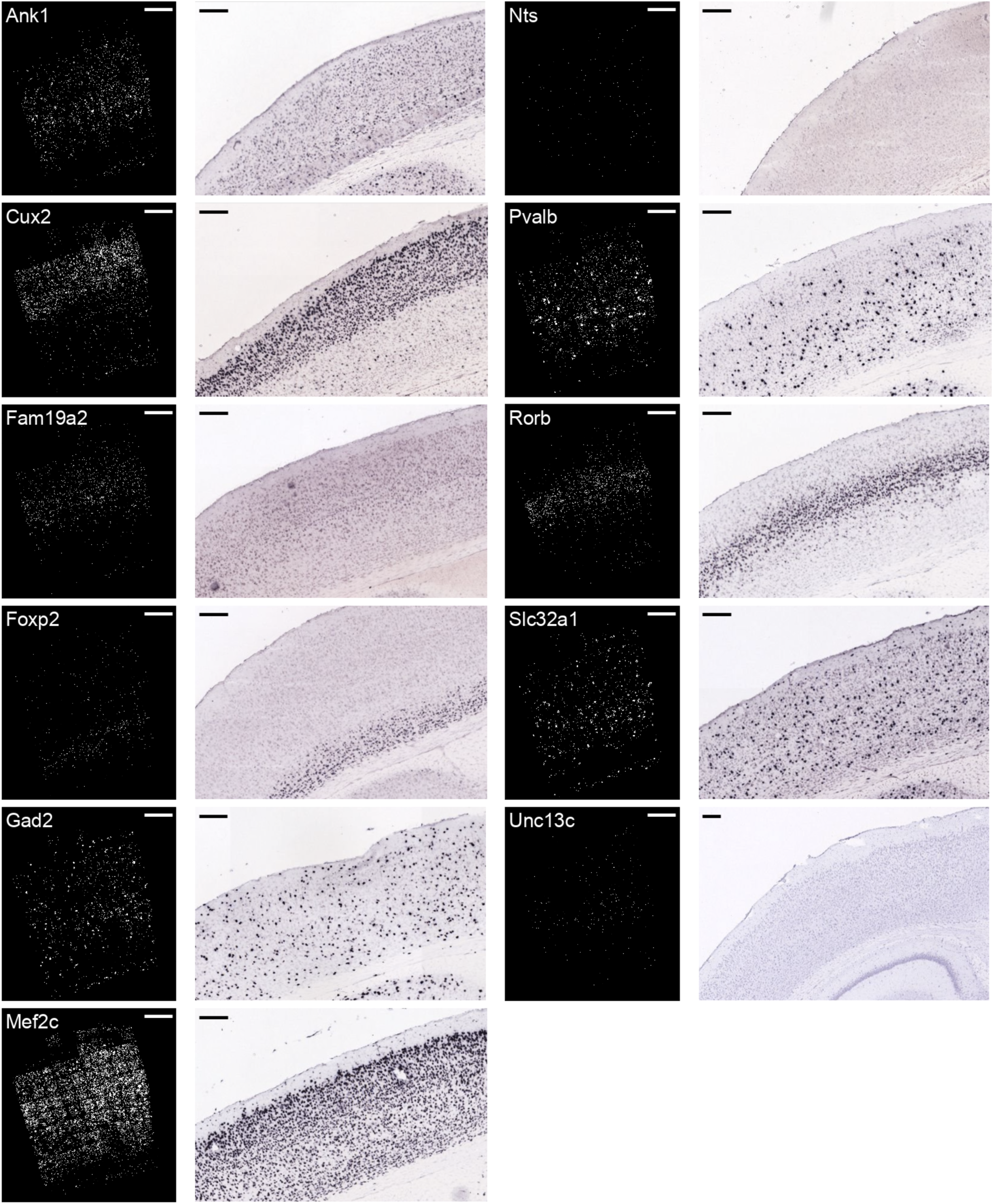
Comparison between targeted ExSeq and Allen Institute ISH atlas (https://mouse.brain-map.org/) for selected marker genes in the primary visual cortex. Scale bars: targeted ExSeq, 200 microns pre-expansion; Allen Institute ISH, 200 microns.

**Fig. S14.**
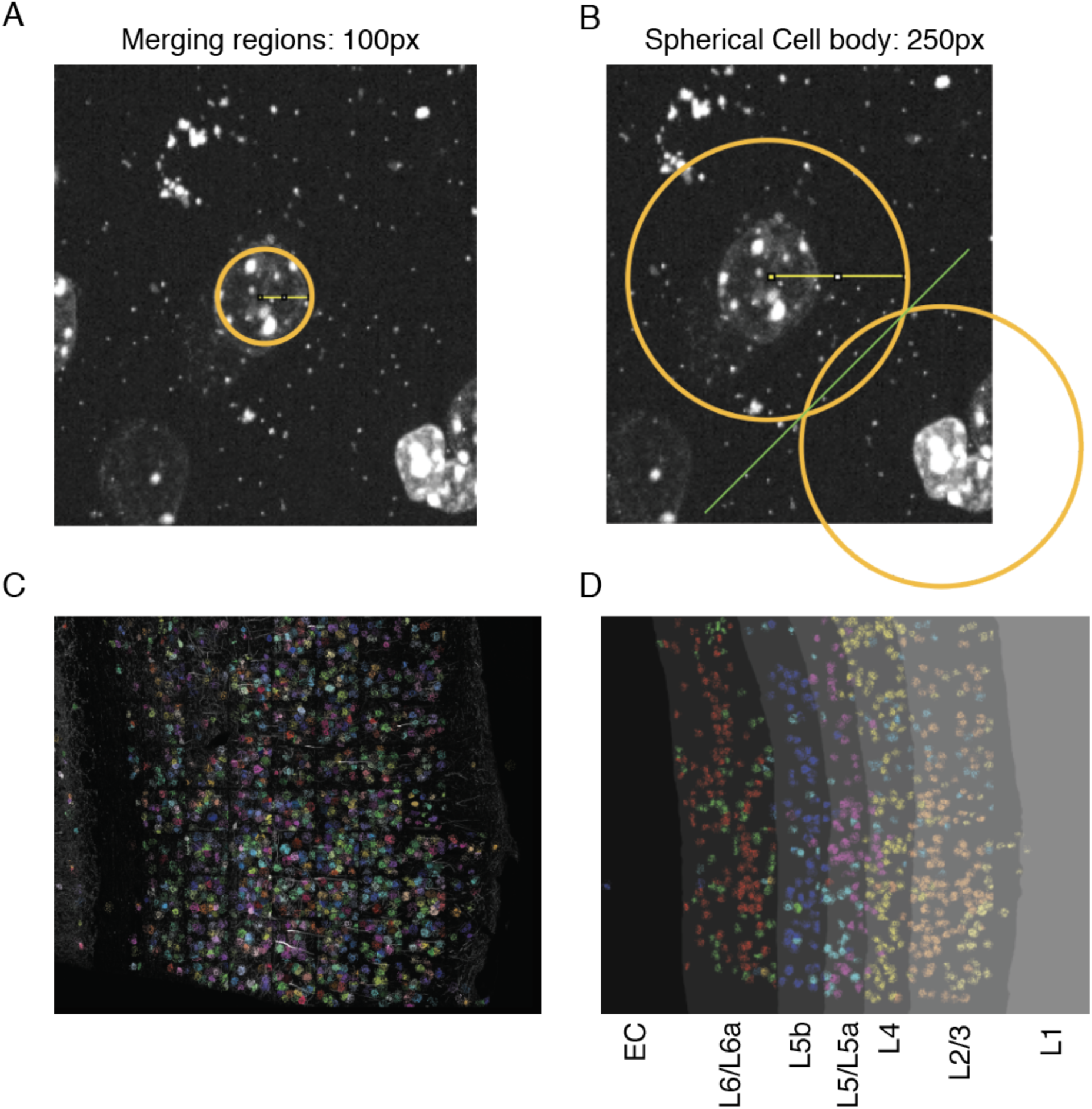
Cell and layer segmentation in the visual cortex. (A) DAPI staining marks putative nuclei objects, which were merged together if the centroids of the objects were within a radius of 5 microns (pre-expansion, 100 px) (shown). (B) Reads within 12.5 microns (pre-expansion) of a centroid were ascribed to that cell; if two centroids were within 12.5 microns of a read (as shown), the reads were ascribed to their nearest neighbor. (C) Demonstration of the cell segmentation pipeline by coloring all reads assigned to an individual cell with a random color. (D) The segmentation of the visual cortex into layers. The excitatory neuron clusters from Fig. 4D**-E** are shown, alongside the layer segmentation in gradations of grey.

**Fig. S15.**
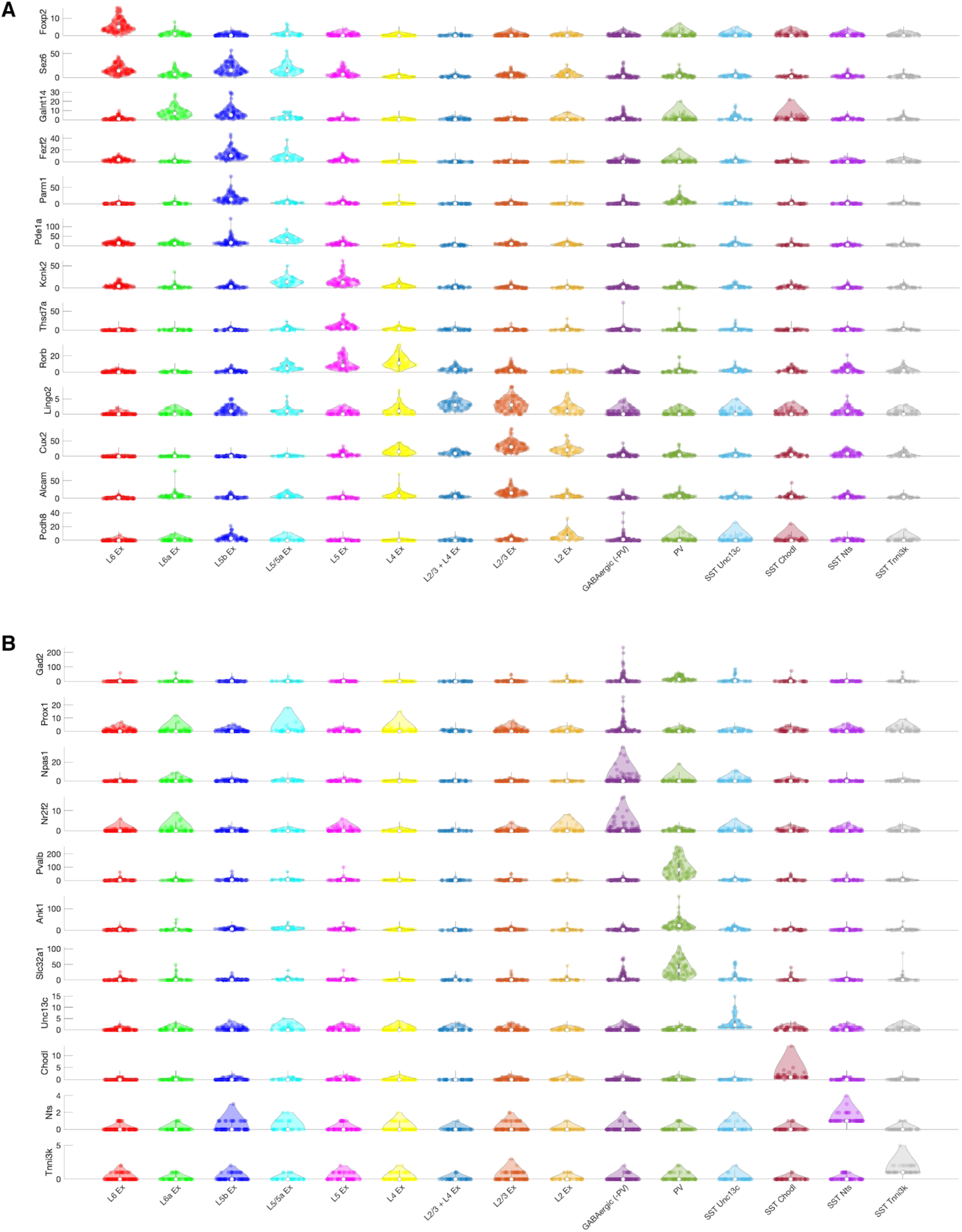
Expression of marker genes in the visual cortex clusters. (A) Violin plots of markers for excitatory neurons in Fig. 4D, showing the number of reads for each marker gene in each cell within a cluster. (B) Violin plots of markers for inhibitory neurons identified in Fig. 4D, showing the number of reads for each marker gene in each cell within a cluster. These results are consistent with previous studies (including (*58*)). For example: Foxp2 in layer 6 excitatory neurons (L6 Ex) (*129*), Sez6 in layer 5 and 6 excitatory neurons (L6 Ex, L6a Ex, L5b Ex, L5/L5a Ex, L5 Ex) (*130, 131*); Kcnk2 in layer 5 excitatory neurons (L5/L5a Ex, L5 Ex) (*132, 133*); Lingo2 in layer 2/3 excitatory neurons (L2 Ex, L2/3 Ex, L2/3 + L4 Ex) (*58*); Cux2 in layer 2/3 excitatory neurons (L2/3 Ex, L2 Ex) (*38*); Pvalb in PV inhibitory neurons (PV) (*134*); Prox1 in VIP inhibitory neurons (GABAergic (-PV)) (*58*); and Chodl in a subset of SST neurons (SST Chodl) (*58*).

**Fig. S16.**
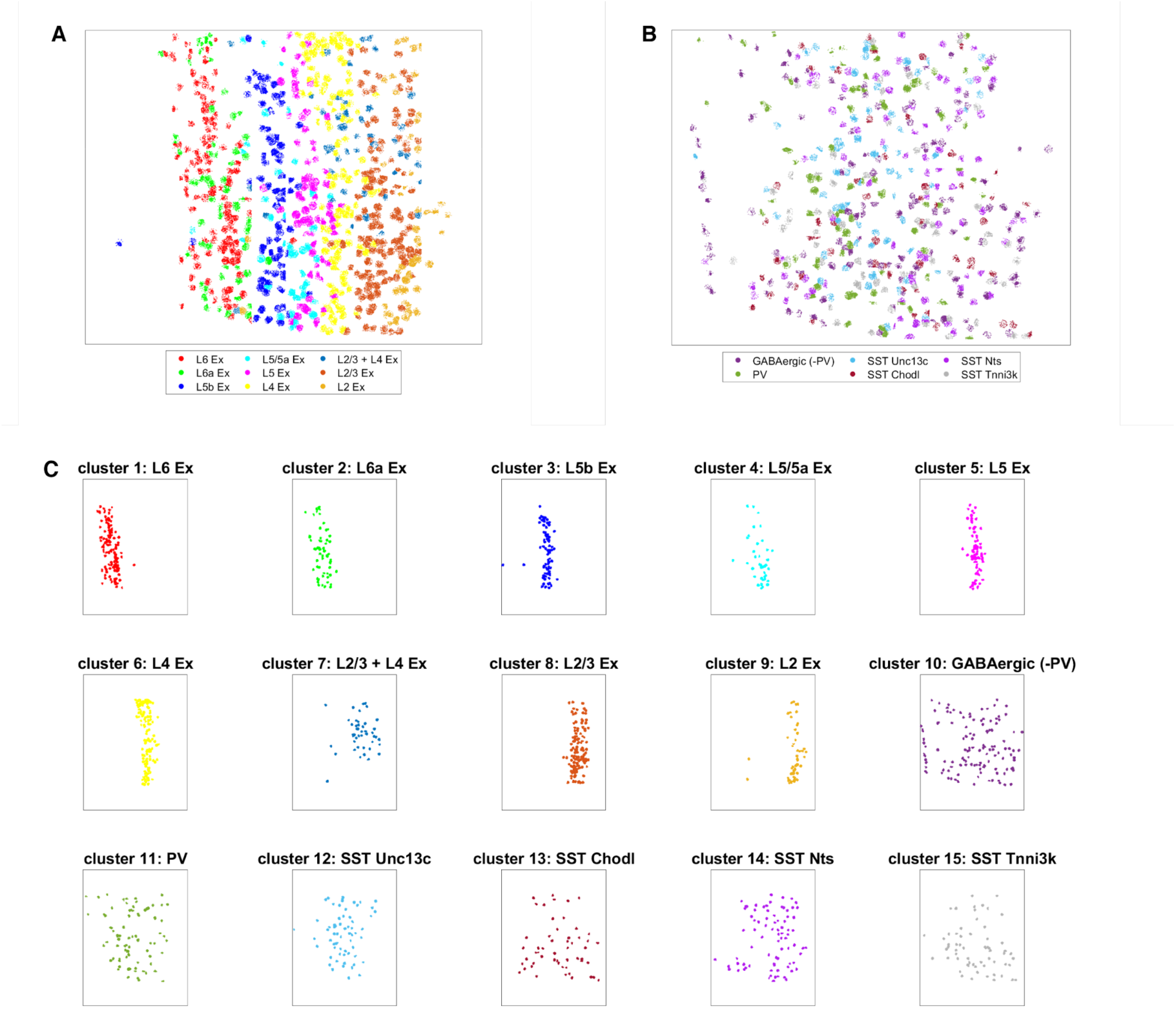
Spatial organization of clusters in the visual cortex. (A) The reads corresponding to the excitatory neuron clusters are colored by their cluster identity and plotted in their spatial locations in the visual cortex slice. (B) Reads corresponding to inhibitory neurons are colored by their cluster identity and plotted in their spatial locations in the visual cortex slice. (C) Reads within cells of each cluster are shown independently, plotted in their spatial locations in the visual cortex slice.

**Fig. S17.**
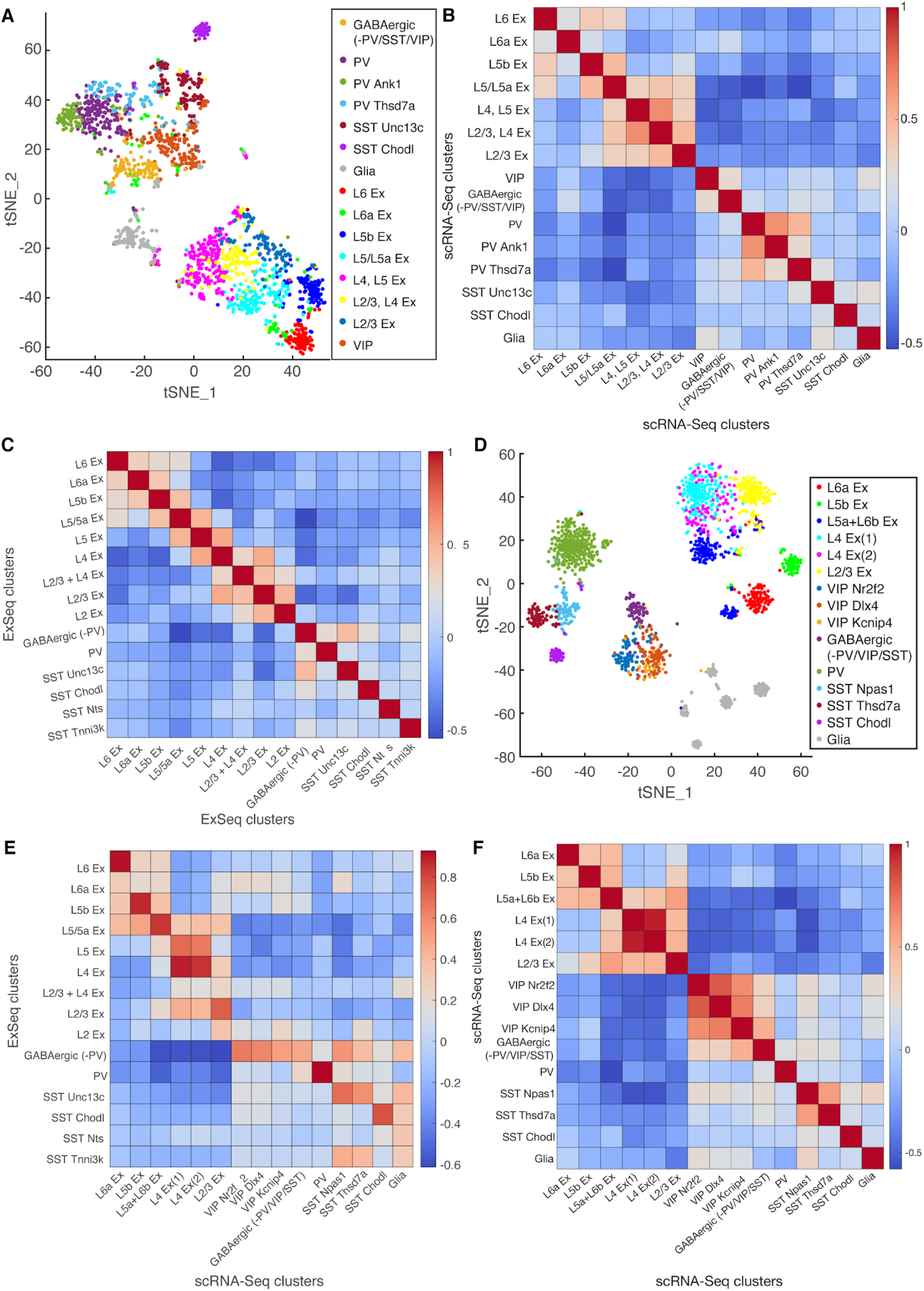
Additional validation of single-cell clustering approach. (A) t-SNE plot of the scRNA-Seq single-cell dataset used in the clustering analysis (*58*), restricted to the 42 genes utilized for the targeted ExSeq of visual cortex. (B) Heatmap showing Pearson’s correlation between pairs of single-cell clusters from A. (C) Heatmap showing Pearson’s correlation between pairs of ExSeq clusters. (D) t-SNE plot of the scRNA-Seq single-cell dataset used in the clustering analysis, clustered using variable genes (see **Methods**). (E) Heatmap showing Pearson’s correlation between ExSeq clusters and single-cell clusters generated using variable genes. (F) Heatmap showing Pearson’s correlation between pairs of scRNA-Seq single-cell clusters, when they are clustered using highly variable genes, in D.

**Fig. S18.**
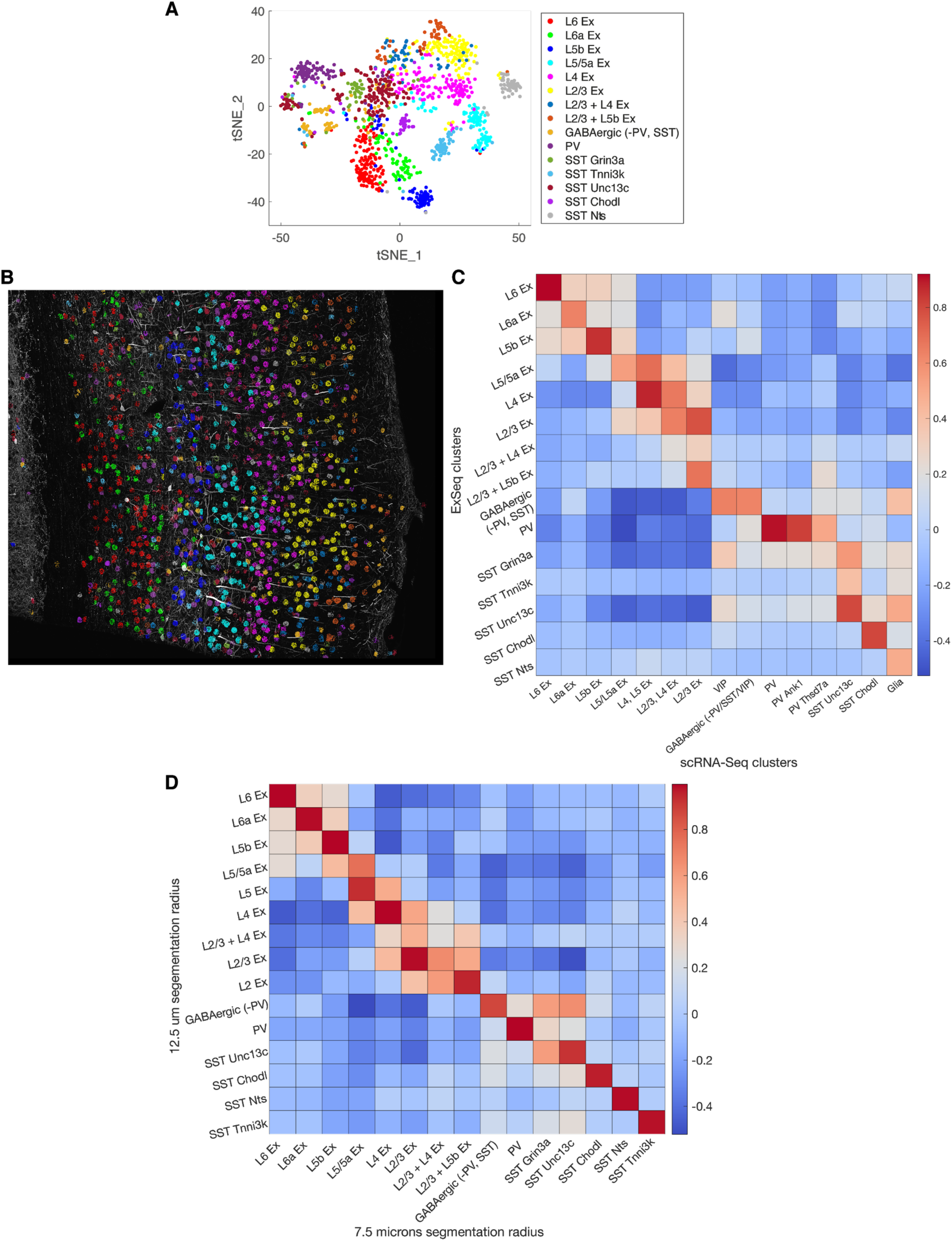
Robustness of cell segmentation. (A) t-SNE plot of targeted ExSeq gene expression profiles when cells were processed with a segmentation radius of 7.5 microns (compared to 12.5 microns in Fig. 4D). (B) Spatial organization of cell types identified in (A). Cell-segmented reads are shown, colored by cluster assignment, and overlaid on the YFP morphology (white). (C) Heatmap of Pearson’s correlation between targeted ExSeq clusters with physical radius 7.5 microns, as in A, and clusters identified in the single-cell RNA-Seq dataset (*58*). (D) Heatmap of Pearson’s correlation between targeted ExSeq clusters using a physical radius of 12.5 microns for cell segmentation (as in Fig. 4D**-G**) and targeted ExSeq clusters using a radius of 7.5 microns for cell segmentation, as in A.

**Fig. S19.**
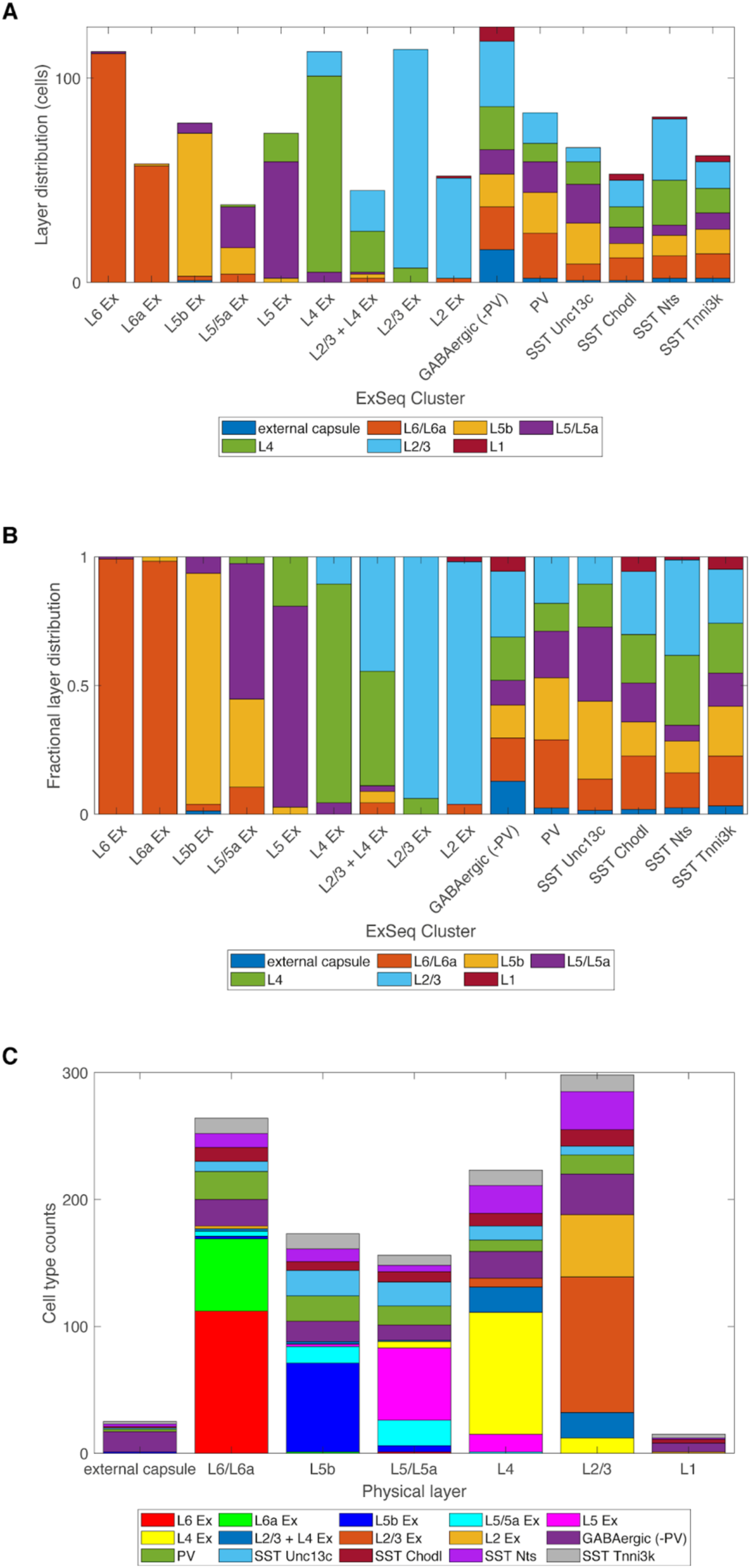
Organization of cell type clusters into layers. (A) Physical layer distribution of clusters by absolute number of cells. (B) Fractional layer distribution of clusters. (C) Cell type counts by physical layer. Whereas the excitatory neuron clusters are each preferentially found in one layer (not surprisingly, because these clusters define the layers), the inhibitory neuron clusters (‘GABAergic (-PV)’, ‘PV’, ‘SST Unc13c’, ‘SST Chodl’, ‘SST Nts’, ‘SST Tnni3k’) are more evenly distributed. For example, cells in the “GABAergic (-PV)” cluster were relatively evenly distributed across layers L2-L6.

**Fig. S20.**
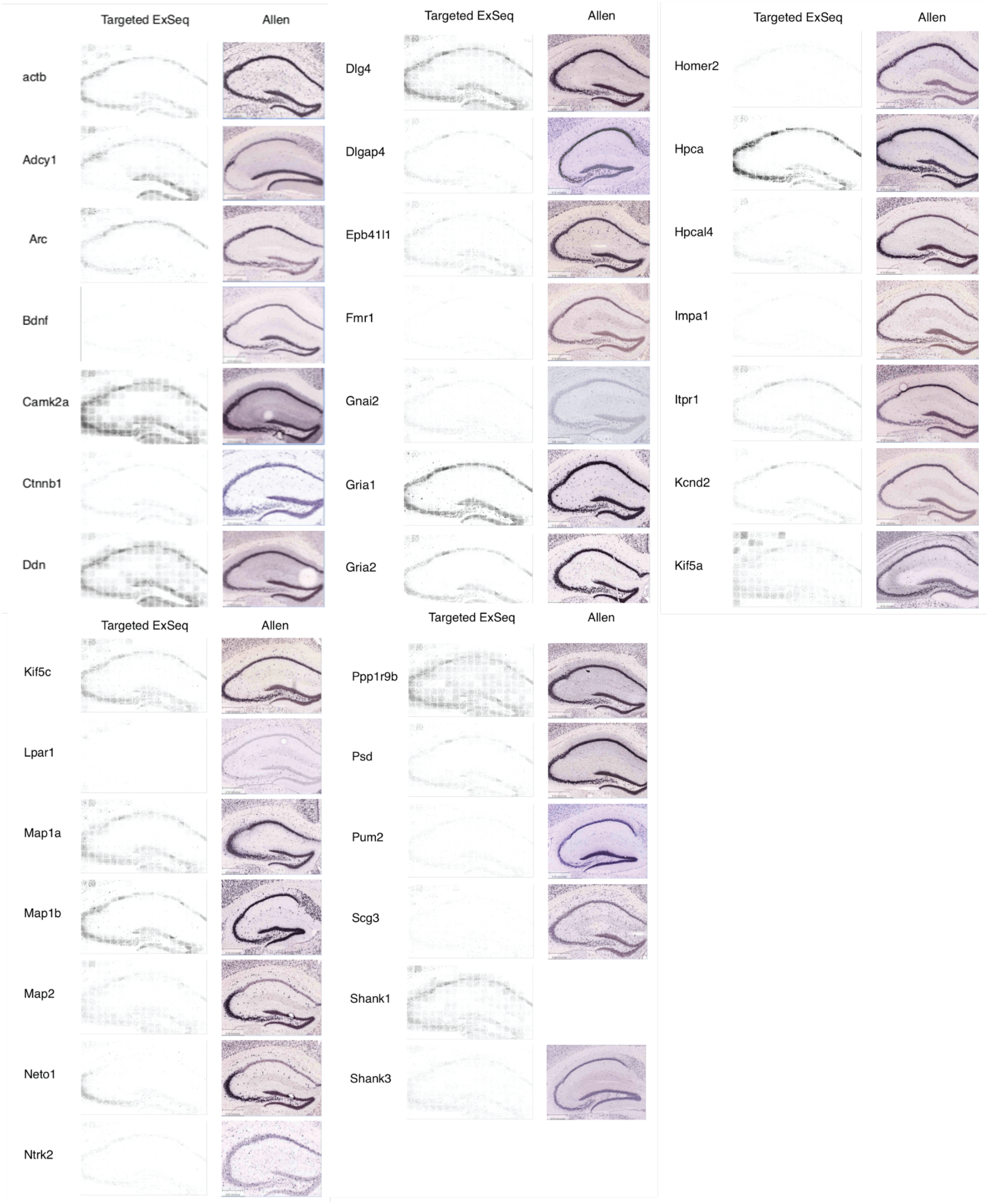
Gene expression patterns in the mouse hippocampus identified via targeted ExSeq are similar to those observed in the Allen Institute Brain Atlas (https://mouse.brain-map.org/). These series of images show transcript localization in the hippocampus via targeted ExSeq (left), comparing them to their respective coronal slice images in the Allen Institute Brain Atlas in situ hybridization dataset (right).

**Fig. S21.**
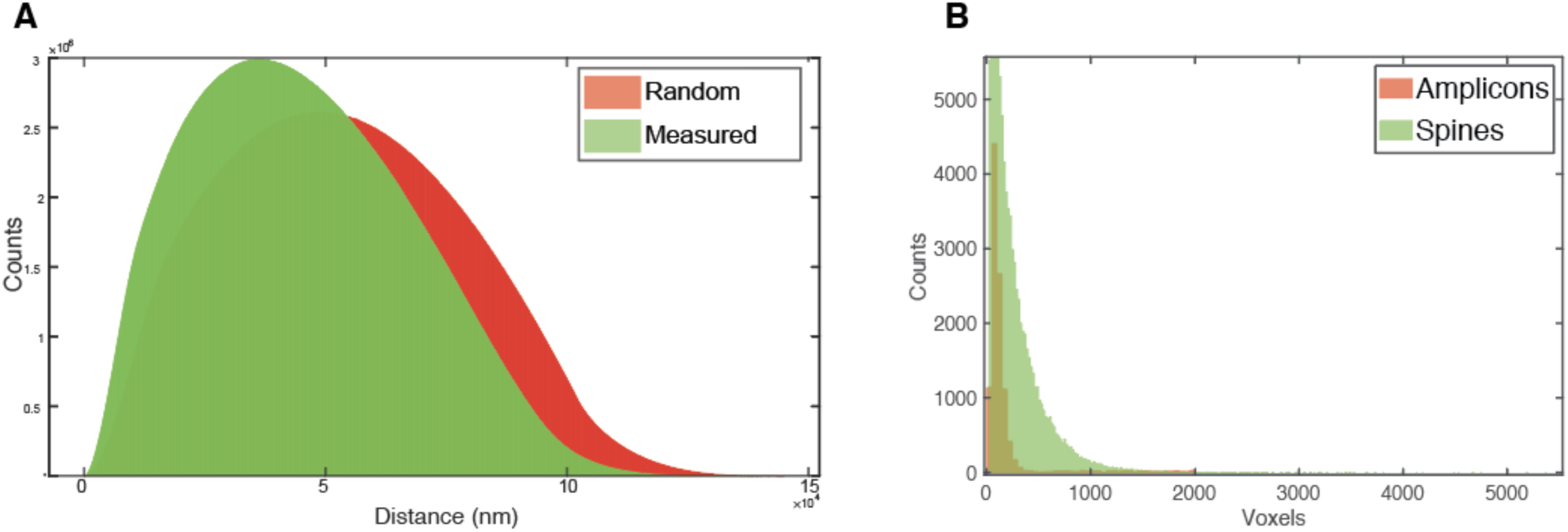
Analysis of amplicon formation density and volume, in the hippocampus targeted ExSeq dataset. (A) Histogram showing measured pairwise distances between targeted ExSeq amplicons in the hippocampal dataset (green) (associated with Fig. 5), and histogram showing the distribution of pairwise distances when the same amplicons were randomly placed within the same volume (red). The inability of amplicons to form close to one another (e.g., if forming one amplicon “laterally inhibits” the formation of a nearby amplicon) would have resulted in the measured pairwise distances tapering off quickly closer to 0, which we do not see here. (B) Histograms showing the distribution of volumes for spine heads (green) in the hippocampal dataset (Fig. 5) and amplicons (red). Based on these volumes, it is possible for multiple amplicons to occupy the same spine head, indicating that our observation of one amplicon per spine reflects the infrequent presence of the transcripts studied here within spines.

**Fig. S22.**
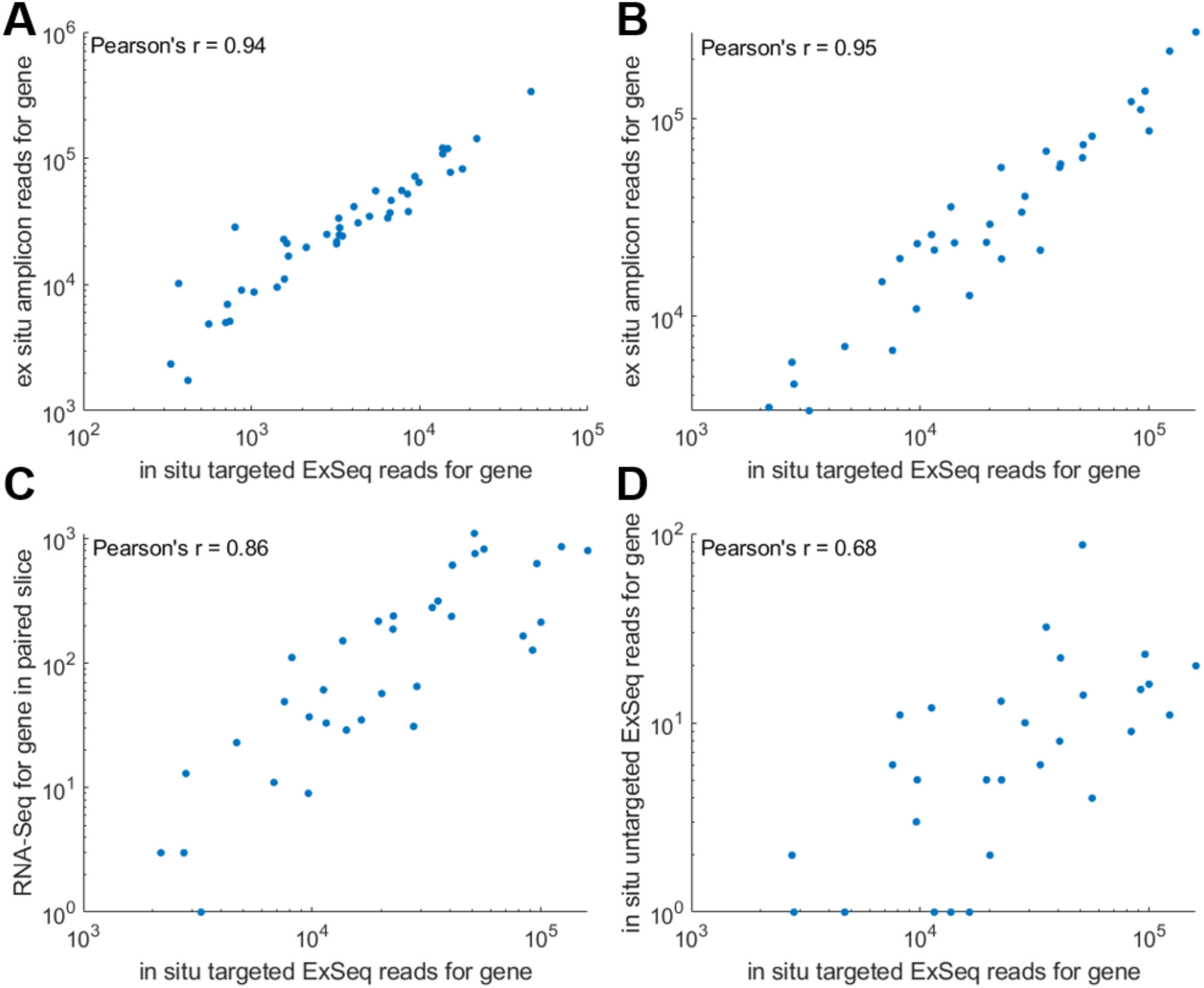
Targeted ExSeq reads are highly correlated with *ex situ* amplicon sequencing, RNAseq, and untargeted ExSeq datasets. (A) Scatterplot showing targeted ExSeq counts for transcripts studied in Fig. 4 (mouse visual cortex tissue) along with their abundance upon *ex situ* extraction and barcode sequencing of amplicons from the same sample (see **Methods** section ‘Comparison of targeted ExSeq to *ex situ* sequencing and bulk RNAseq’). Each point is a gene among the 42 genes examined in the visual cortex dataset (Pearson’s r = 0.94, p-value = 8.68 × 10^-21^; counts in **Table S12**). (B) Scatterplot showing targeted ExSeq counts for transcripts studied in Fig. 5 (mouse hippocampus tissue) along with their abundance upon *ex situ* extraction and barcode sequencing of amplicons from the same sample. Each point is a gene among the 34 genes examined in the hippocampal dataset (Pearson’s r = 0.95, p-value = 2.85 × 10^-18^; counts in **Table S13**). (C) Scatter plot showing targeted ExSeq counts for transcripts studied in Fig. 5 against their expression with RNAseq using a paired (i.e., adjacent) hippocampal slice (Pearson’s r = 0.86, p-value = 9.38 × 10^-11^; counts in **Table S13**). (D) Scatter plot showing targeted vs untargeted ExSeq counts for the genes studied in Fig. 5 in the mouse hippocampus, taken from two different mice but from identical coordinates (Pearson’s r = 0.68, p-value = 9.75 × 10^-6^; counts in **Table S13**).

**Fig. S23.**
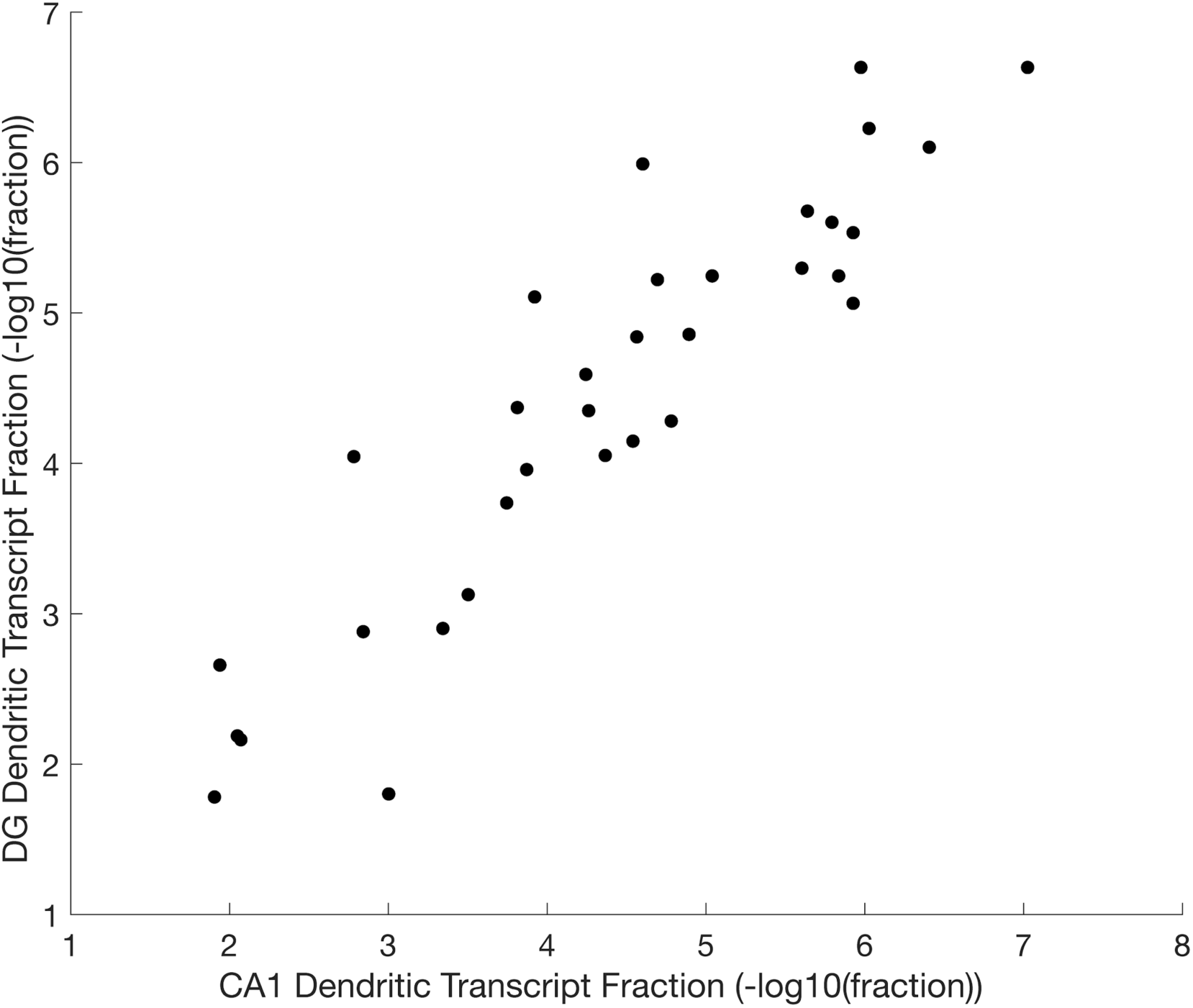
High correlation between the dendritic expression of genes for granule cells of the dentate gyrus (DG) vs. pyramidal neurons of CA1 (Pearson’s r 0.91; p-value 1.2 × 10^-13^). Scatter plot showing the ‘dendritic transcript fraction’, defined as the number of reads of a given gene that are located in the dendrites divided by the total number of reads found in dendrites (for all examined genes combined), in the dentate gyrus vs CA1 for the genes studied in Fig. 5 using targeted ExSeq.

**Fig. S24.**
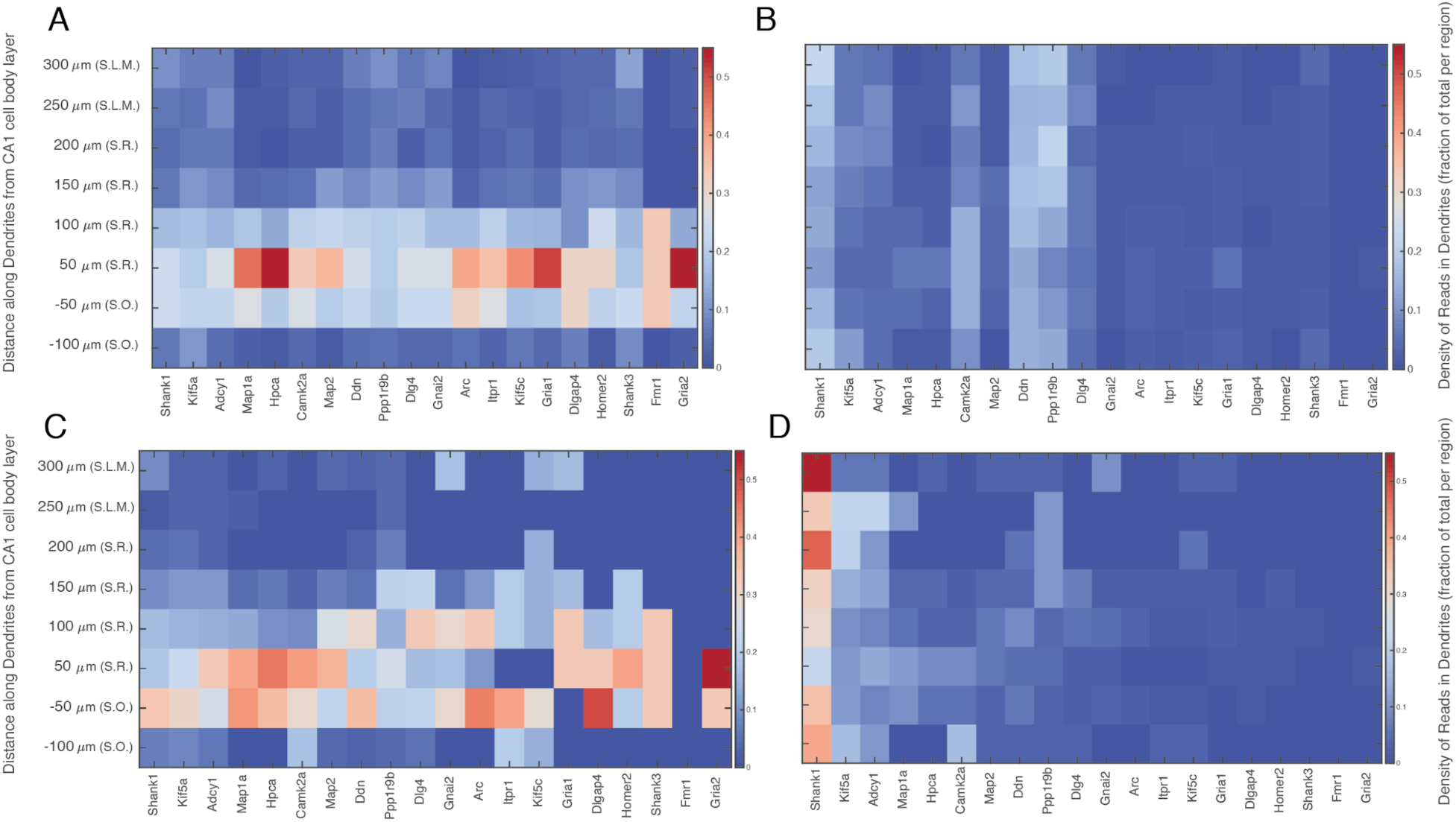
Heatmaps showing the density of transcripts in the dendrites (A,B) and spines (C,D) of CA1 pyramidal neurons corresponding to Fig. 5E along the apical-basal axis (Euclidean distance) of hippocampal area CA1, normalized by either the total transcript counts per gene (for A and C), or by the total transcript counts at a given distance from the cell body layer (for B and D).

**Fig. S25.**
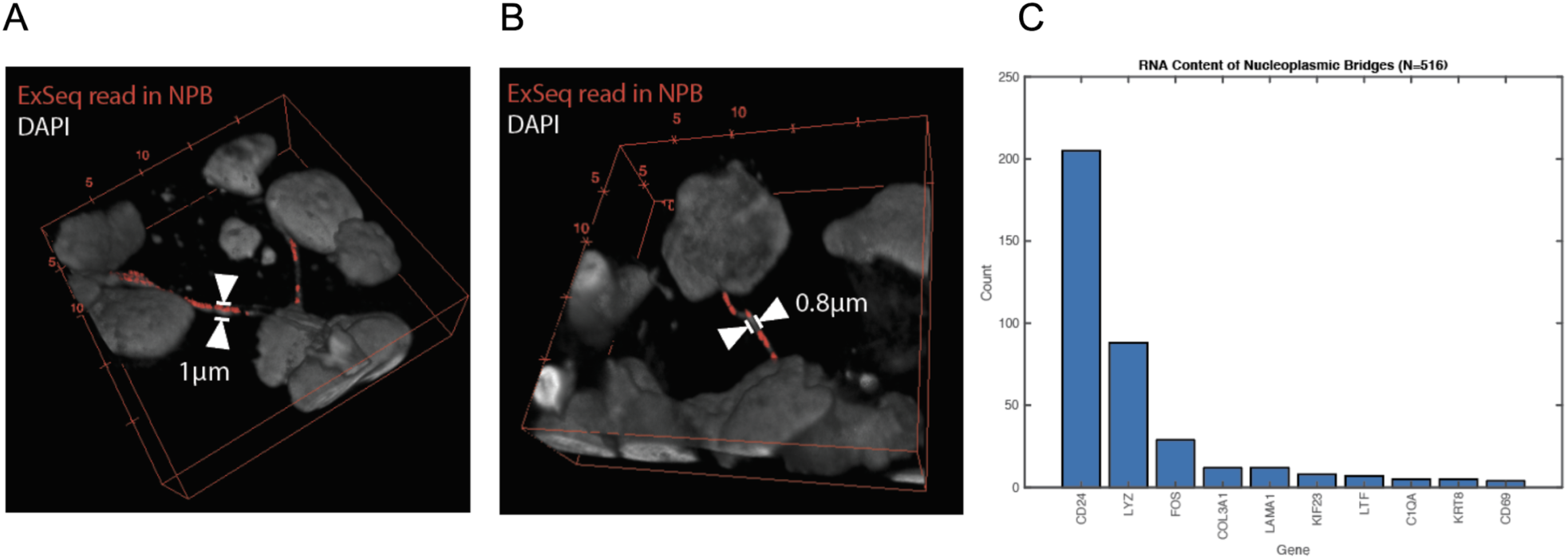
Targeted ExSeq of a human metastatic breast tumor biopsy reveals transcripts within nuclear structures (possibly nucleoplasmic bridges) that are under one micron in size. (A,B) 3D renders showing ExSeq reads (red) localized within the volumes of the nuclear structures overlaid on a confocal image of DAPI nuclear staining. Arrows show the nuclear structures and their width. (C) Bar plot showing the abundance of genes localized within the nuclear structures. A total of 516 reads were identified in 83 nuclear structures. We note that the relatively small number of reads obtained in these nanoscale regions limited our ability to systematically classify these reads into specific cell types.

**Fig. S26.**
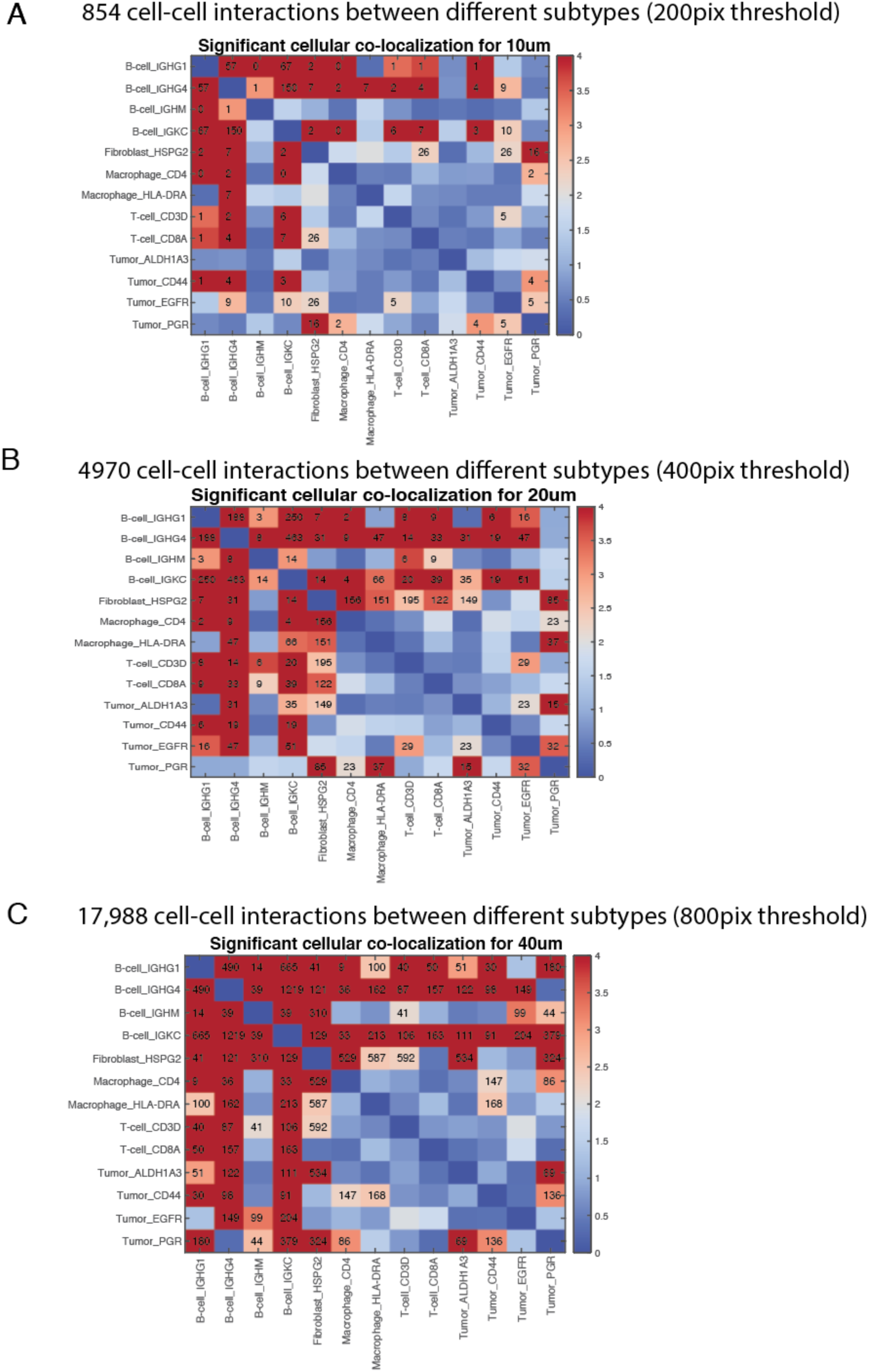
The cell clusters in metastatic breast cancer tissue exhibit robust, non-random spatial colocalizations. The adjacency matrix text values indicate the number of cell pairs across different clusters that are in close proximity as determined by a Euclidean distance threshold between cell centroids of 10 (A), 20 (B) and 40 (C) microns. The adjacency matrix heatmap shows the p-value (10,000 bootstrapping iterations) relative to obtaining the same or higher number of cells in close proximity by chance. All adjacency matrix entries with text values are statistically significant (Bejamini Hochberg false-discovery rate of 1.5%). Note that the structure of the matrix is preserved across the different thresholds.

**Table S7.**
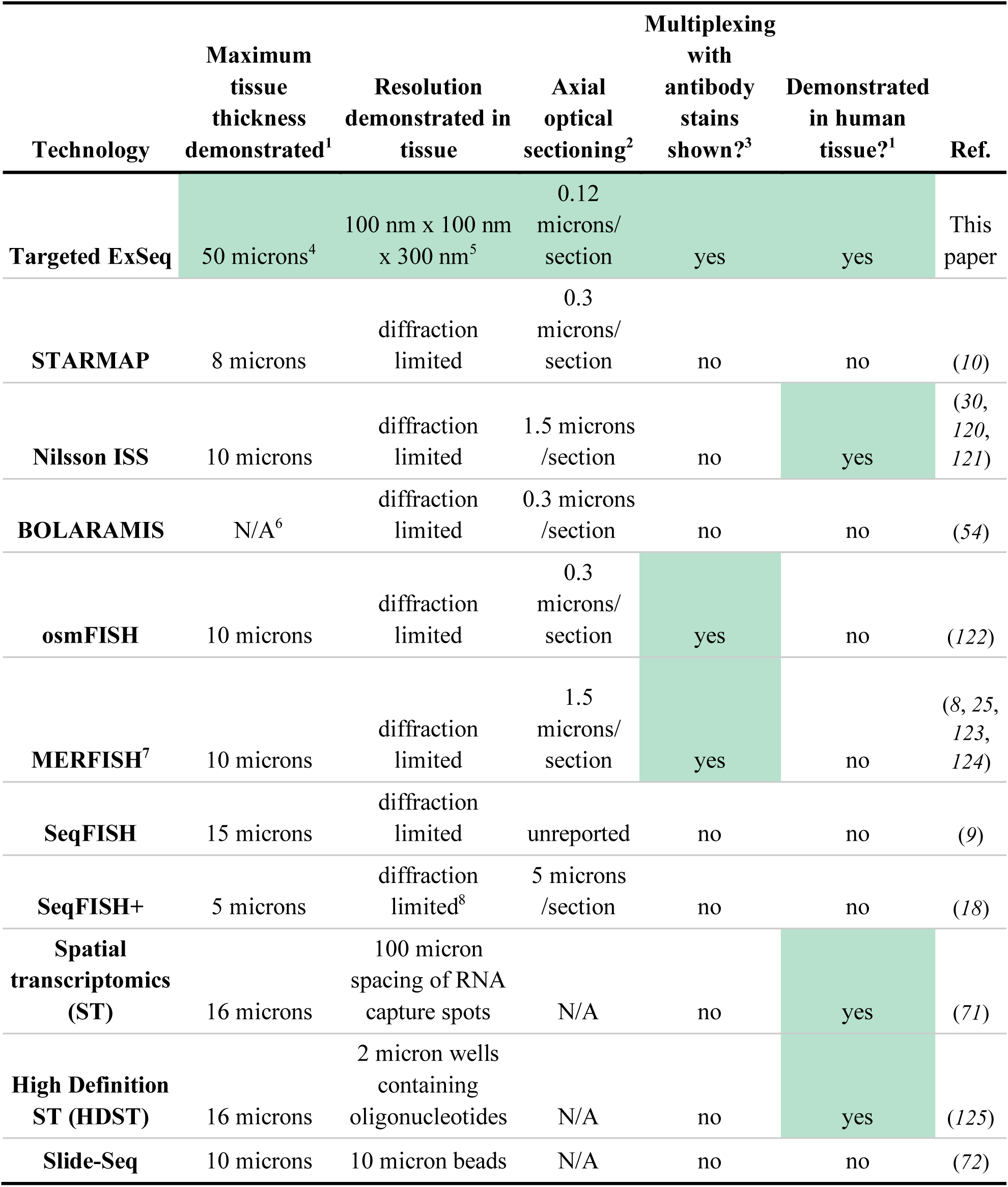
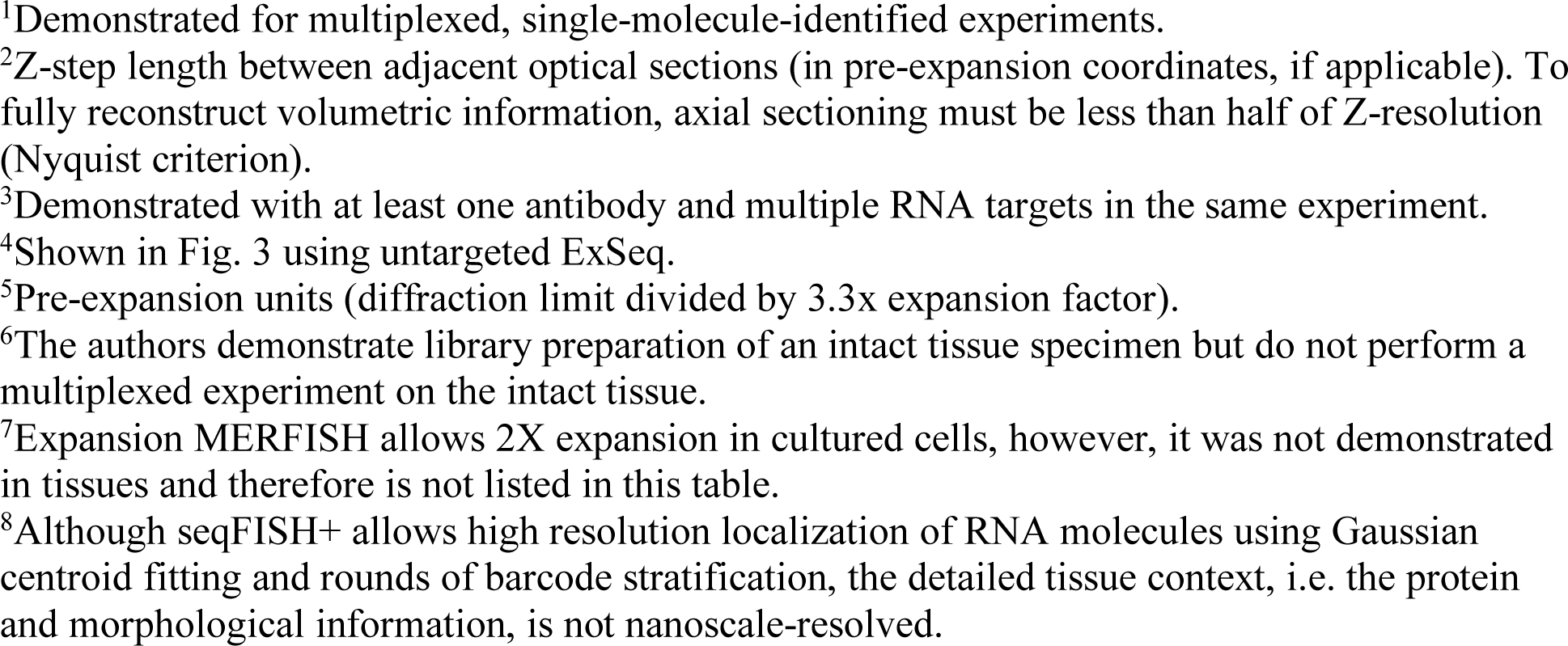
Evaluation of highly multiplexed targeted transcriptomic technologies for various criteria. Highlighted in green are the technologies with the best result for each examined criterion. Not shown in this table are untargeted ExSeq, FISSEQ (*23*) and INSTA-seq (*50*) which are untargeted technologies (as defined by their use of random primers to initialize their library preparation for *in situ* sequencing).

| Technology | Maximum tissue thickness demonstrated<sup>1</sup> | Resolution demonstrated in tissue | Axial optical sectioning<sup>2</sup> | Multiplexing with antibody stains shown?<sup>3</sup> | Demonstrated in human tissue?<sup>1</sup> | Ref. |
| --- | --- | --- | --- | --- | --- | --- |
| Targeted ExSeq | 50 microns<sup>4</sup> | 100 nm x 100 nm x 300 nm<sup>5</sup> | 0.12 microns/section | yes | yes | This paper |
| STARMAP | 8 microns | diffraction limited | 0.3 microns/section | no | no | (10) |
| Nilsson ISS | 10 microns | diffraction limited | 1.5 microns/section | no | yes | (30, 120, 121) |
| BOLARAMIS | N/A<sup>6</sup> | diffraction limited | 0.3 microns/section | no | no | (54) |
| osmFISH | 10 microns | diffraction limited | 0.3 microns/section | yes | no | (122) |
| MERFISH<sup>7</sup> | 10 microns | diffraction limited | 1.5 microns/section | yes | no | (8, 25, 123, 124) |
| SeqFISH | 15 microns | diffraction limited | unreported | no | no | (9) |
| SeqFISH+ | 5 microns | diffraction limited<sup>8</sup> | 5 microns/section | no | no | (18) |
| Spatial transcriptomics (ST) | 16 microns | 100 micron spacing of RNA capture spots | N/A | no | yes | (71) |
| High Definition ST (HDST) | 16 microns | 2 micron wells containing oligonucleotides | N/A | no | yes | (125) |
| Slide-Seq | 10 microns | 10 micron beads | N/A | no | no | (72) |

<sup>1</sup>Demonstrated for multiplexed, single-molecule-identified experiments.
<sup>2</sup>Z-step length between adjacent optical sections (in pre-expansion coordinates, if applicable). To fully reconstruct volumetric information, axial sectioning must be less than half of Z-resolution (Nyquist criterion).
<sup>3</sup>Demonstrated with at least one antibody and multiple RNA targets in the same experiment.
<sup>4</sup>Shown in Fig. 3 using untargeted ExSeq.
<sup>5</sup>Pre-expansion units (diffraction limit divided by 3.3x expansion factor).
<sup>6</sup>The authors demonstrate library preparation of an intact tissue specimen but do not perform a multiplexed experiment on the intact tissue.
<sup>7</sup>Expansion MERFISH allows 2X expansion in cultured cells, however, it was not demonstrated in tissues and therefore is not listed in this table.
<sup>8</sup>Although seqFISH+ allows high resolution localization of RNA molecules using Gaussian centroid fitting and rounds of barcode stratification, the detailed tissue context, i.e. the protein and morphological information, is not nanoscale-resolved.

**Table S8.**
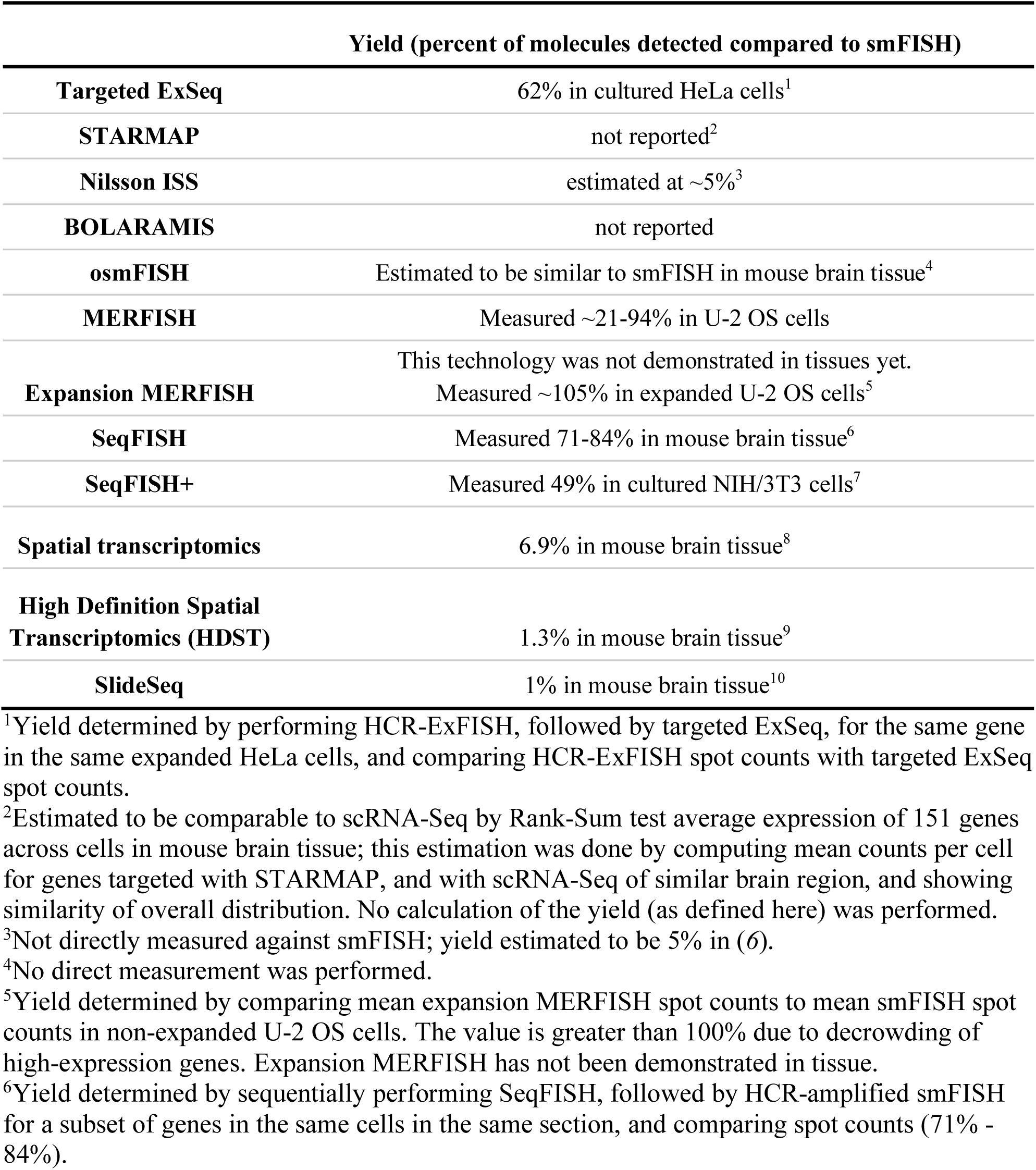
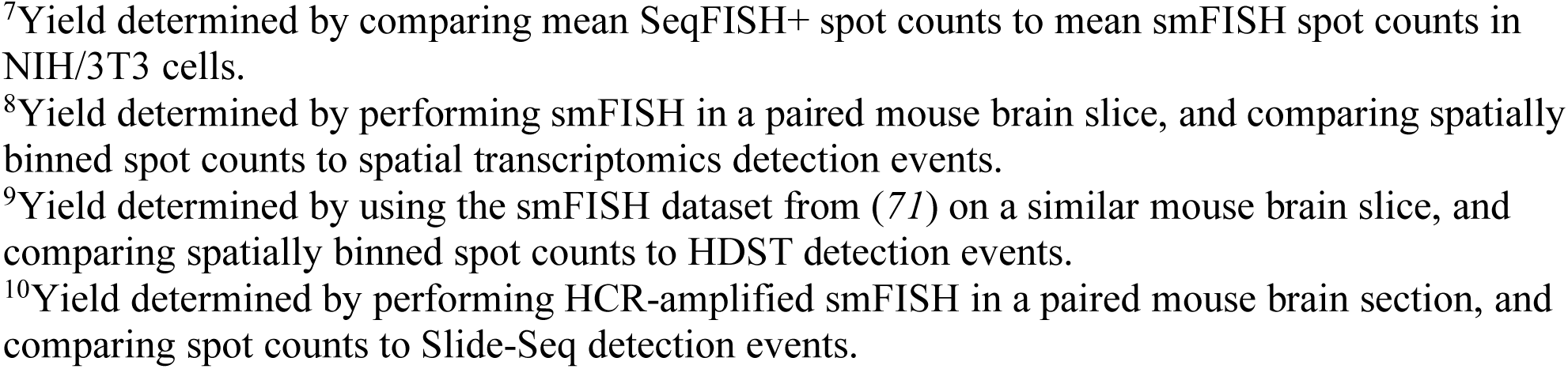
Comparison of yield, i.e. percent of molecules detected compared to single-molecule FISH (smFISH), and methods of determining yield for highly multiplexed, targeted, RNA localization technologies.

